# Transgene-Free Ex Utero Derivation of A Human Post-Implantation Embryo Model Solely from Genetically Unmodified Naïve PSCs

**DOI:** 10.1101/2023.06.14.544922

**Authors:** Bernardo Oldak, Emilie Wildschutz, Vladyslav Bondarenko, Alejandro Aguilera-Castrejon, Cheng Zhao, Shadi Tarazi, Mehmet-Yunus Comar, Shahd Ashouokhi, Dmitry Lokshtanov, Francesco Roncato, Sergey Viukov, Eitan Ariel, Max Rose, Nir Livnat, Tom Shani, Carine Joubran, Roni Cohen, Yoseph Addadi, Merav Kedmi, Hadas Keren-Shaul, Sophie Petropoulos, Fredrik Lanner, Noa Novershtern, Jacob H. Hanna

**Author notes:** These authors contributed equally (Bernardo Oldak, Emilie Wildschutz, Vladyslav Bondarenko). Correspondence: Jacob H. Hanna.

## Abstract

Our ability to study early human post-implantation development remains highly limited due to the ethical and technical challenges associated with intrauterine development of the human embryo after implantation. Despite the great progress made on human gastruloids, axioloids and in vitro cultured blastoids, such elegant models do not constitute an integrated Stem cell-derived Embryo Models (SEMs) that includes all the key extra-embryonic tissues of the early post-implantation human conceptus (e.g., hypoblast, yolk-sac, trophoblasts, amnion, and extraembryonic mesoderm), and thus, do not recapitulate post-implantation epiblast development within the context of these extra-embryonic compartments. Mouse naïve pluripotent stem cells (PSCs) have recently been shown to give rise to embryonic and extra-embryonic stem cells capable of self-assembling into post-gastrulation mouse SEMs, while bypassing the blastocyst-like stage, and eventually initiating organogenesis *ex utero*. Here, we implement critical adaptations to extend these finding to humans, while using only genetically unmodified human naïve PSCs, thus circumventing the need for ectopic expression of lineage promoting transgenes. Such integrated human SEMs recapitulate the organization of all known compartments of early post-implantation stage human embryos, including epiblast, hypoblast, extra-embryonic mesoderm, and trophoblast surrounding the latter layers. The organized human SEMs recapitulate key hallmarks of post-implantation stage embryogenesis up to 13-14 days post-fertilization (dpf, Carnegie stage 6a), such as bilaminar disk formation, epiblast lumenogenesis, amniogenesis, anterior-posterior symmetry breaking, PGC specification, primary and secondary yolk sac formation, and extra-embryonic mesoderm expansion that defines a chorionic cavity and a connective stalk. This new platform constitutes a tractable stem cell-based model for experimentally interrogating previously inaccessible windows of human peri- and early post-implantation development.

## Introduction

Our understanding of human early post-implantation development has been limited due to difficulties in obtaining relevant embryo samples from these early stages of human gestation, and technical and ethical challenges for *in vitro* development of donated blastocysts towards the post-implantation stages while preserving the *in vivo* complexity of both embryonic and extra embryonic compartments ^1, 2^. Most of our knowledge of these stages has been gained solely from histological and anatomical descriptions in embryological collections, and the experimental models for mechanistic research are lacking ^3, 4^. Despite the important recent advances in culturing the *in vitro* attached human embryos beyond implantation, these culture systems are restricted in sample number and, overall, do not support normal development of embryos beyond the initiation of early epiblast lumenogenesis ^5–7^.

The ability to capture human pluripotent stem cells (PSCs) in different developmental states in culture ^8–10^ has opened the possibility of investigating early human embryogenesis using 3D stem-cell derived embryo models. By controlling cellular composition and differentiation conditions, the recently introduced elegant models such as gastruloids, blastoids, axioloids, or amniotic sac embryoids ^11–23^ recapitulate some aspects of early mammalian development in a modular manner. Among them, Blastocyst-like structures (blastoids) are the only currently available human integrated (i.e., comprised of all embryonic and extra-embryonic lineages) model of the human pre-implantation embryo, containing epiblast, primitive endoderm (PrE), and trophectoderm (TE) ^22–25^. However, so far, the attempts to advance the blastoids towards post-implantation stages through *in vitro* culture did not progress beyond what has been achieved with natural human blastocysts ^5–7, 22–25^. Similarly, mouse blastoids do not develop into bona fide post-implantation embryos even when transferred *in utero* ^26^.

Recently, mouse PSCs were shown to possess the ability to be coaxed *ex utero* into post-gastrulation-stage SEMs, by co-aggregation of non-transduced naïve PSCs with naïve PSCs transiently expressing the transcription factors Cdx2 and Gata4, to promote their priming towards TE and PrE lineages, respectively ^27, 28^. Such SEMs do not go through a blastocyst or an *in vitro* implantation-like stages but develop directly into egg-cylinder shaped SEMs within complex extra-embryonic compartments and are able to advance beyond gastrulation and reach early organogenesis stages of development as late as day E8.5 ^27, 28^. These findings establish that naïve pluripotent cells ^29–31^ can serve as the sole source of embryonic and extra-embryonic tissues in advanced bona fide “organ-filled” embryo models, and thus may enable the generation of integrated SEMs from other mammalian species from which naïve PSCs have been stabilized, including humans. Given that conditions for human naive pluripotency have been continuingly improved and optimized ^32–38^, here we sought to test whether and how an equivalent approach could be implemented with human naive PSCs to obtain human post-implantation SEMs that unequivocally acquire anatomical compartments of pre-gastrulating human embryo with lineage marker expression and an adequate emergence of the key developmental milestones.

### Optimizing human naive PSC priming towards extra-embryonic lineages competent for post-implantation SEM formation

In mouse, deriving SEMs that contain critical embryonic and extraembryonic compartments requires optimal conditions and high-quality rapid induction of Primitive endoderm (PrE) and Trophectoderm (TE) from naïve pluripotent stem cells (PSC), which was successfully achieved by ectopic expression of Gata4 and Cdx2, respectively ^27, 28, 39, 40^. The latter feat^27^ offered the crucial and exclusive platform for us to demonstrate and unleash the self-organizing capacity of mouse stem cells to generate post-gastrulation bona fide synthetic embryos with both organs and extra-embryonic compartments in a unique ex utero set-up and only by starting from naive PSCs. This latter fact is of added importance, since it showed that the establishment of pluripotent stem cells only, may be sufficient to make advanced embryo-like structures in the petri dish from many mammals, including humans, without the need to derive TSC or PRE/XEN lines from human embryo derived material, some of which are not yet established in humans or did not prove competent to generate post-implantation mouse SEMs.

Hence, we first set out to establish a similar platform to obtain extraembryonic lineages through transient expression of these transgenes in human naïve PSCs (**Fig. 1a**), while bearing in mind that the early post-implantation pre-gastrulation human, but not mouse, embryo already contains an extra embryonic mesoderm compartment (ExEM) ^41–47^. To generate DOX (Doxycycline) inducible human PSCs for GATA4 or GATA6, regulators of PrE in mouse ^48, 49^ and of Pre and ExEM lineages in humans ^50^, we used a PiggyBac system carrying M2Rtta and transcription factor of interest (**Extended Data Fig. 1a**). Monoclonal and polyclonal human ESC and iPSCs lines were validated for GATA4 or GATA6 expression upon DOX addition (**Extended Data Fig. 1b**). We aimed to use FACS staining for cell surface expression of PDGFRa, which marks both PrE and ExEM lineages in humans ^50^, as an initial screening point for identifying optimal conditions to rapidly and efficiently induced PrE and/or ExEM cells, which will be followed by secondary characterization and validation. While Gata4 induction (iGata4) in mouse naïve 2i/LIF conditions yielded dramatic upregulation of PDGFRa+ cell fraction after 48 hours of DOX supplementation (**Extended Data Fig. 1c**) ^51^, induction of GATA4 and GATA6 expression in human naive PSCs cultured in HENSM conditions, resulted <10% PDGFRa+ cells even after 6 days of follow-up (**Fig. 1b, Extended Data Fig. 1d**). Since WNT stimulation by GSK3B inhibition via CHIR99021 (CH) has been shown to be the key stimulant for mouse and human PrE induction ^52–54^, and given that CH is included in mouse, but not in recent human naïve media conditions ^35, 55, 56^, we thought that HENSM conditions during the induction phase might not be suitable for human cells. Hence, we screened for other conditions to facilitate the induction of PDGFRa+ cells from naive human PSCs (**Fig. 1a**). The recently described mouse PrE-derivation conditions (termed C10F4PDGF) ^54^, resulted in very low yield of PDGFRa induction (Fig.1c). RACL induction media (**R**PMI based medium supplemented with **A**CTIVIN A, **C**HIR99021 and **L**IF) ^57^ which has been used to prime human naïve PSCs towards PrE and ExEM state ^50, 58^, or NACL media (DMEM/F12/Neurobasal **N**2B27-based) that stabilizes naïve Endoderm (nEnd) cells generated in RACL conditions ^57^, also led to low levels of PDGFRa+ cell fraction even after six days of induction in RACL (**Fig. 1c, middle**) or NACL (**Extended Data Fig. 1e**).

**Figure 1.**
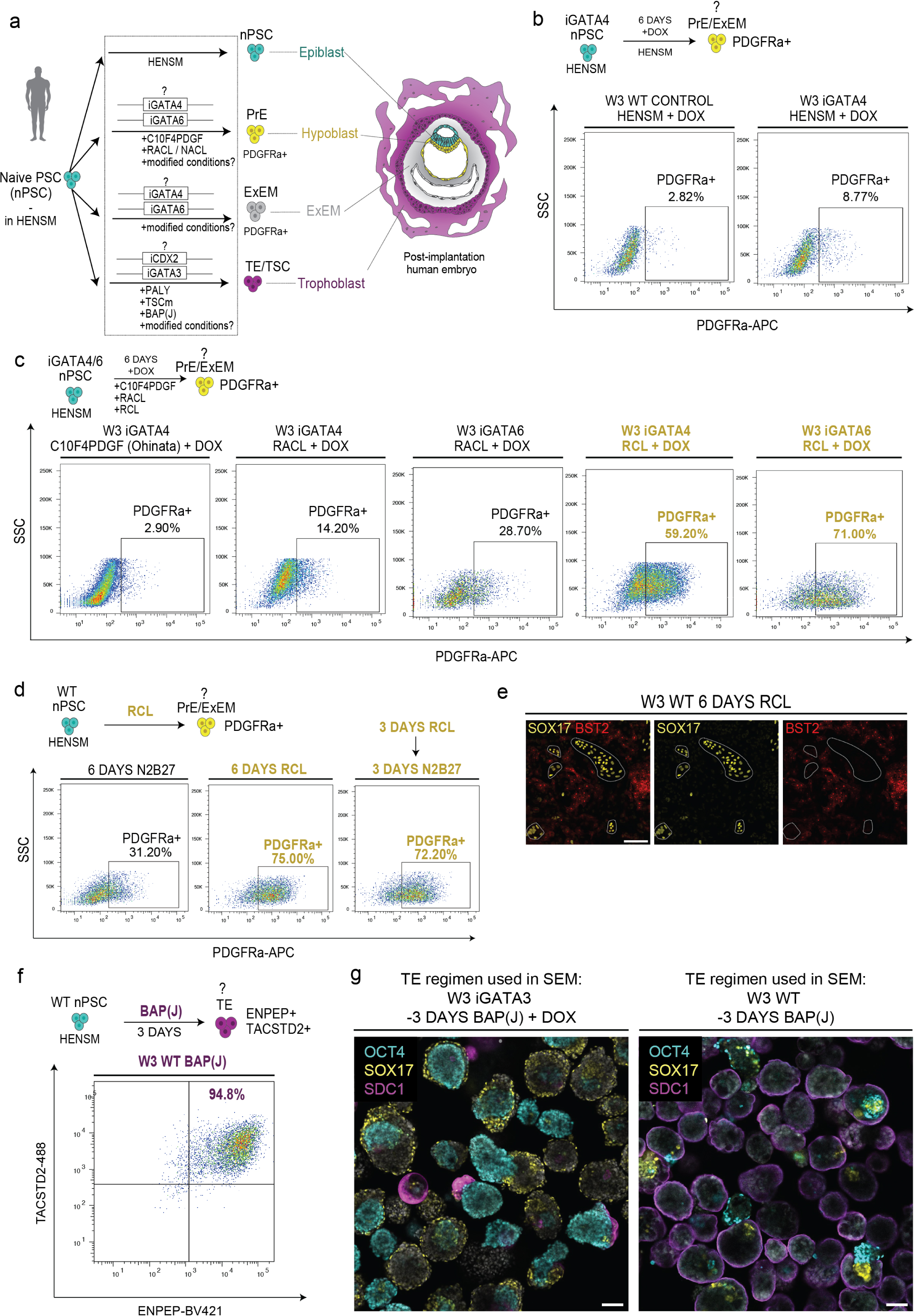
Optimizing human naive PSC differentiation towards extra-embryonic lineages competent for early post-implantation SEM generation. **a,** left, scheme of the tested induction (iGATA4, iGATA6, iCDX2, iGATA3) and media conditions for generating the three different needed extraembryonic lineages from naïve pluripotent stem cells (nPSC). Right, scheme of the early human post-implantation embryo with the epiblast (cyan), hypoblast (yellow), ExEM (grey), and trophoblast (magenta) compartments. **b,** Flow Cytometry (FACS) plots of PDGFRa-APC versus SSC for PrE/ExEM induction using iGATA4 with DOX in HENSM after 6 days (right) and the control condition of naïve cells (WT cells without iGATA4, left). **c,** from left to right, FACS plots of PDGFRa-APC for PrE/ExEM induction using iGATA4 and iGATA6 with DOX for 6 days in different media conditions as indicated (C10F4PDGF, RACL, RCL). **d,** from left to right, FACS plots of PDGFRa-APC for PrE/ExEM using WT naïve PSCs induced 6 days in different media conditions (N2B27, RCL, and RCL for 3 days followed by 3 days of N2B27). **e,** immunofluorescence images of WT nPSCs induced in RCL media for SOX17 (yellow) and BST2 (red). Two distinct cell populations emerge with mutually exclusive expression pattern (outlined); scale bar, 100 µm. **f,** FACS plots of ENPEP against TACSTD2 for trophectoderm (TE) lineage induction of HENSM naïve PSCs in BAP(J) regimen for 3 days. **g,** immunofluorescence images of day 6 SEM aggregates stained for OCT4 (cyan), SOX17 (yellow) and SDC1 (magenta). Aggregates made with iGATA3 in DOX for 72h in BAP(J), showed no outer surrounding trophoblasts in the obtained aggregates (left). In contrast, WT nPSCs in BAP(J) showed very high efficiency in yielding aggregates with outer layer of surrounding trophoblast (marked by SDC1 – magenta). Scale bars 100 µm. RACL **R**PMI based medium with **C**HIR99021, ACTIVIN, and **L**IF; RCL, **R**PMI based medium with **C**HIR99021 and **L**IF without ACTIVIN; **H**ENSM, **H**uman **E**nhanced **N**aïve **S**tem cell **M**edia; BAP(J), DMEM/F12 based medium with ALK4/5/7 inhibitor **A**83-01, ERKi/MEKi PD0325901, and **B**MP4 substituted with **J**AK inhibitor I after 24h.

Since ACTIVIN A is not required for priming cells towards PrE in mouse ^54^ and inhibits human naïve PSC in vitro differentiation into ExEM cells ^50^, we omitted it from RACL condition (now termed RCL). Remarkably, RCL media resulted in up to 60% PDGFRa+ cells in iGATA4 cells and up to 80% PDGFRa+ from iGATA6 cells within 3-6 days (**Fig. 1c, right**). The slightly improved outcome when using Gata6 overexpression is consistent with previous reports placing Gata6 as the most upstream regulator of PrE identity in mouse and human cells ^48, 59^. Remarkably however, high efficiencies of PDGFRa+ cell formation was evident in RCL conditions from isogenic wild type (WT) cells without exogenous expression of GATA6 (**Fig. 1d**), indicating that the transient transgene expression is not required for efficient PDGFRa+ induction in human naïve PSC. Further optimization showed that 3 days of induction in RCL condition followed by 3 days incubation in basal N2B27 conditions yielded comparable results (**Fig 1d – right, Extended Data Fig. 1g)**). Notably, incubating naïve PSCs in N2B27 media also yielded PDGFRa+ cells albeit at significantly lower levels (**Fig 1d, left**), and thus we focused on using RCL conditions for further characterization analysis. The B27 supplement devoid of insulin in RCL was utilized herein as it yielded slightly higher efficiency in PDGFRa+ cell induction compared to B27 with insulin (**Extended Data Fig. 1f**), consistent with previous reports ^57^.

Our next step was to test for existence of PDGFRa+ PrE and/or ExEM cells in our RCL induction protocol and to distinguish between them. Both immunostaining and RT-PCR analysis validated endogenous expression of PrE marker genes in RCL conditions, including SOX17 which marks only the PrE fraction, alongside GATA4, GATA6, and NID2 genes (**Extended Data Fig. 2 a, b)** ^57^. Markers of definitive endoderm like GSC or HHEX were not induced from naïve PSCs in RCL conditions (**Extended Data Fig. 3a**) thus excluding definitive endoderm cell formation ^57^. Applying RCL conditions on human isogenic primed PSCs yield predominant definitive endoderm marker expression (GSC or HHEX) (**Extended Data Fig. 1h**), consistent with previous findings ^57^. Immunostaining results for SOX17 and BST2 markers that distinguish between PrE or ExEM lineages ^52, 60^, respectively, confirmed the co-emergence of PrE and ExEM from naïve PSCs under the same RCL induction protocol (with or without GATA6 overexpression) (**Fig. 1e, Extended Data Fig. 3b**). GATA4 positively marked both SOX17+ PrE and BST2+ ExEM populations as expected (**Extended Data Fig. 3c**) ^50^. BST2+ ExEM cell identity was further validated by upregulation of FOXF1 protein which marks ExEM cells, but not PrE cells or residual PSCs (**Extended Data Fig. 3d**) ^61, 62^. Thus, we adopted RCL pre-treatment of naïve PSCs for co-aggregation experiments with other lineages (**Fig. 2b**).

**Figure 2.**
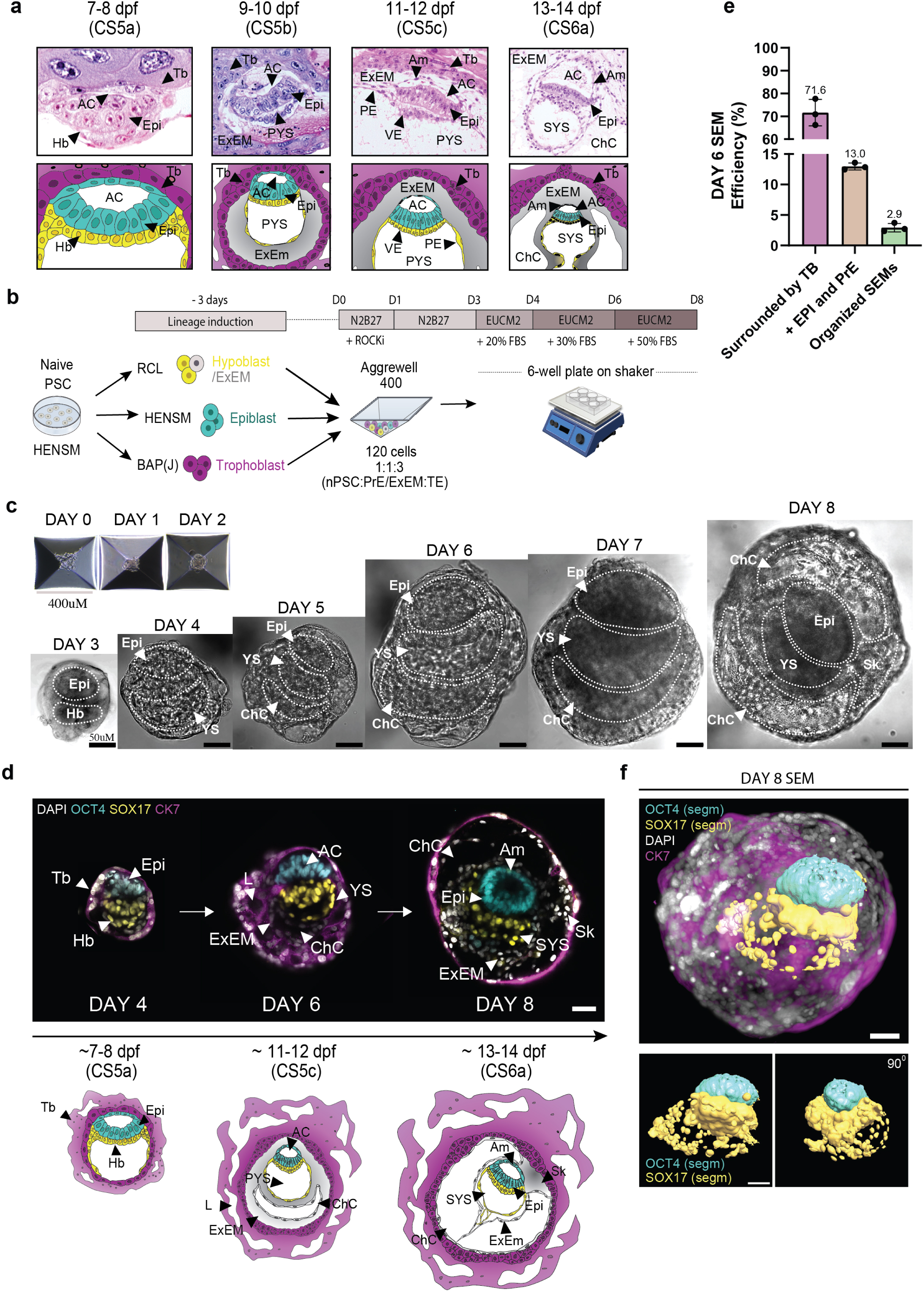
Self-assembly of human Post-Implantation SEM exclusively from non-transgenic naïve PSCs. **a**, anatomical sections taken from the Carnegie collection and schemes of the human embryo development between 7 and 14 days post-fertilization (dpf); left to right, Carnegie Stages (CS) CS5a (7-8 dpf), CS5b (9-10 dpf), CS5c (11-12 dpf), and CS6a (13-14 dpf). **b**, scheme of the protocol for generating human early post-implantation SEMs. Hypoblast/ExEM (yellow/grey), epiblast (cyan), and trophoblast (magenta) lineage priming for 3 days from naïve hPSCs in HENSM media is followed by aggregation (counted as day 0) in Aggrewell 400 at the 1:1:3 nPSC: PrE/ExEM: TE ratio in N2B27. Starting from day 3, SEMs are transferred to non-adherent 6-well plates on a shaker and cultured in *Ex-Utero* Culture Medium 2 (EUCM2) with increasing concentrations of FBS until day 8 (see Methods). **c**, representative brightfield images of SEMs growing between days 0 and 8 showing formation of the inner embryonic structures, which were defined based on the lineage-specific immunofluorescence (see **Extended Data Figure 12b**). Scale was maintained to show the growth. **d**, from left to right, immunofluorescence images of SEMs from days 4, 6, and 8 of the protocol, showing OCT4 (cyan), SOX17 (yellow), CK7 (magenta), and nuclei (DAPI, white), labeling different embryonic structures (marked with arrows). **e**, quantification of the SEM derivation efficiency across three independent experimental replicates. **f**, 3D reconstruction of the day 8 SEM shown in d (right) with segmented epiblast, (OCT4, cyan), hypoblast (SOX17, yellow), CK7 (magenta) and nuclei (DAPI). Bottom, segmentation of the epiblast and hypoblast shown in 0 and 90^0^ degrees of rotation. See also **Video S1.** Epi, epiblast; AC, amniotic cavity; Am, amnion; TE, trophectoderm; Tb, trophoblast; PrE, primitive endoderm; Hb, hypoblast; PYS, primary yolk sac; VE, visceral endoderm; PE, parietal endoderm; SYS, secondary yolk sac; ExEM, extraembryonic mesoderm; ChC, chorionic cavity; Sk, stalk; L, lacunae. Scale bars, 50 µm.

In parallel, we set out to find optimal conditions for deriving cells form the trophectoderm (TE) lineage that can adequately aggregate with the other lineages and develop into post-implantation stage human SEMs ^63^. Overexpression of Cdx2, a master regulator of TE lineage, can lead to formation of TSC lines in mice ^39, 64^. Moreover, its overexpression in mouse naive PSCs has been shown to be efficient in generating TSCs that remained viable and correctly integrated upon coaggregation with mouse naïve PSCs and iGata4 cells and generated both chorionic and ectoplacental cone placental lineages ^27, 28^ (**Extended Data Figure 4a-b**). Cdx2 has also been shown to be transiently expressed in the TE of the human blastocysts and blastoids ^6, 7, 22, 65^. Therefore, we generated DOX inducible CDX2 human ESC and iPSC lines using a similar PiggyBac approach (**Extended Data Fig. 4c, d**), which were also subjected to lentiviral labeling with a constitutively expressed tdTomato marker to better track cell viability and integration patterns (**Extended Data Fig. 4e**). Remarkably, human iCDX2 cells pre-treated with DOX for 1 – 4 days in either naïve HENSM or in different validated human TE/TSC induction conditions: PALY medium (N2B27 supplemented with PD0325901, A83-01, and Y-27632) ^22^, BAP(J) induction medium (DMEM/F12 based medium with ALK4/5/7 inhibitor A83-01, FGF2 inhibitor PD173074, and BMP4 substituted with JAK inhibitor I after 24h) ^36, 66^, or TSC maintenance medium (TSCm) ^67^, did not meaningfully contribute and expand within the aggregates generated with induced PrE/ExEM and PSCs. This can be judged by the loss of CK7+ or tdTomato-positive cells in early aggregates (**Extended Data Fig. 4f, h**), likely due to a drastically reduced viability of iCDX2 cells upon treatment with DOX in different conditions and clones used (**Extended Data Fig. 4i**), which was not encountered in mouse cells ^27, 28, 68^. iCDX2 cells also expressed low levels of trophectoderm markers when compared to WT non-transgenic cells (**Extended Data Fig. 4g**)

In the mouse SEM model, established embryo derived TSC lines can substitute the iCdx2-induced PSCs during aggregations without compromising the outcome ^27, 69^. Therefore, we then proceeded to test the established TSC lines derived from both naïve and primed hPSC ^70, 71^. To be able to trace cells after aggregation, TSC lines were either labelled with tdTomato via lentiviral transduction (**Extended Data Fig. 5a**) or stained for CK7 expression. In all employed aggregation conditions with either primed (pTSC) (**Extended Data Fig. 5**) or naïve PSC derived TSC lines (nTSC) (**Extended Data Fig. 6**), the TSCs did not generate an outer layer surrounding the aggregate, but rather formed focal clumps (**Extended Data Fig. 5b – e; Extended Data Fig. 6a – d**). The inability of human TSCs to integrate in putative SEMs, as opposed to mouse TSCs, might stem from the fact that mouse TSC lines are Cdx2+ and correspond to earlier stages of trophoblast development, than those isolated from human naïve or primed TSCs which are CDX2-negative ^70–74^.

Recent studies suggest that GATA3 might have an important role for human TE induction ^75^, thereby, we also established and tested induced GATA3 (iGATA3) human PSCs lines (**Extended Data Fig. 7a–c**) and compared them to differentiation conditions using isogenic genetically unmodified cells. RT-qPCR analysis showed higher expression levels of CDX2 and TACSTD2 in the non-transduced group under BAP(J) conditions (**Extended Data Fig. 7d**). Immunofluorescence staining revealed that although GATA3 overexpression induced endogenous expression of GATA2 TE marker, the cells did not uniformly express TFAP2C, CDX2, and CK7, while in the absence of transgene overexpression we observed higher and uniform expression of TFAP2C, CDX2, and higher occurrence of CK7 positive cells under BAP(J) conditions (**Extended Data Fig. 7e, f**). Flow cytometry analysis for TACSTD2 (marker of early and late TE) and ENPEP (expressed only in the late TE) showed the highest percentage of a double positive population in non-transduced cells under BAP(J) conditions (**Fig. 1f; Extended Data Fig. 7g, right**) ^73^. Addition of BMP4 on the first day of induction showed homogeneous double positive population while exclusion of BMP4 showed that almost half of the population was ENPEP negative (**Extended Data Fig. 7g, right**), consistent with a previous report ^36^. Notably, following the induction and aggregation with naïve PSCs and PrE/ExEM, iGATA3 cells remained viable but did not surround the aggregates (**Fig. 1g, left**). In contrast, TE derived from genetically unmodified naïve PSC under the same BAP(J) protocol, uniformly surrounded the aggregates (**Fig. 1g, right**), which is a critical criterion expected to be fulfilled in integrated SEM ^76^. It is possible that the high ectopic expression produced by the PiggyBac system of GATA3 or CDX2, in addition to the endogenous levels induced by the media used, leads to unfavorably high levels of such factors which might derail the differentiation outcome, and is consistent with the findings in mouse SEMs that cautioned from using highly expressing Cdx2 transgenic clones when generating mouse TE/TSCs ^76, 77^. The ability to derive relevant extra-embryonic lineages from genetically unmodified WT human naive PSCs are without the need for transgene overexpression are in line with the recent studies demonstrating that human naïve pluripotent cells can be more easily coaxed to give rise to early progenitors of PrE cells and TSCs when compared to mouse PSCs, which so far require ectopic transcription factor overexpression ^35, 36, 58, 74^. The latter is consistent with the observation that enhancers ^78^ of key TE and PrE regulators (GATA3, GATA6, GATA4) are accessible in human, but not mouse naïve PSCs while being transcriptionally inactive in both (**Extended Data Figure 8**).

### Human Post-Implantation SEM self-assembled exclusively from non-transgenic naïve PSCs

Having established fast and efficient *in vitro* induction of the three extraembryonic lineages comprising the pre-gastrulating natural human embryo, we proceeded to test their capacity to form embryo-like structures solely from naïve pluripotent stem cells (nPSC) (**Fig. 2a**). To this end, we calibrated the aggregation conditions, such as cell numbers needed and ratios within cell mixtures (**Extended Data Fig. 9a-c**), and the aggregation and growth media composition for different stages of SEM development (**Extended Data Fig. 10**). Unlike for mouse where FBS presence in the first 3 days of SEM formation was critical ^27, 79^, serum free conditions were found favorable for human SEM (**Extended Data Fig. 10b, c**). The protocol starting with 120 cells aggregated at the ratio 1:1:3 (nPSC: PrE/ExEM: TE) in basal N2B27 conditions supplemented with BSA, which was found critical to reduce human aggregate stickiness (**Extended Data Fig 11a**)^80^, for three days resulted in optimal aggregation, validated by the presence of epiblast and extra-embryonic lineages in the aggregates via immunostaining (**Extended Data Fig. 12a**). To further support unrestricted growth of the SEM and prevent TE attachment to the 2D surface, which disrupts the morphology after day 3 of the protocol, we had to continue our culture using orbital shaking conditions and optimal medium volume per well (**Extended Data Fig. 11b-d**) ^27^. The composition of the Ex-Utero Culture Media 2 (EUCM2) was adapted from the mouse SEMs protocol ^27^, while using here incremental FBS concentrations, ranging from 20% at day 3, 30% at day 4-6, and 50% at day 6-8, which were found most optimal for human SEM growth (**Fig. 2b, Extended Data Fig. 12a-b**). We did not observe an improved outcome when using roller-culture applied after day 6 with the same media composition, compared to orbital shaking placed in static incubators (**Extended Data Fig. 11e**) ^27^, which is consistent with observations in mice where roller culture platform was needed particularly for late gastrulation and organogenesis stages ^81^.

To study the development of our human SEMs in further detail and with an adequate reference, we mostly relied on the data from the Virtual Human Embryo Project based on the Embryo Carnegie Stages as the most detailed and relevant source available to date ^82, 83^ (see Methods). During eight days of *ex utero* culture, the aggregates grew extensively, forming a 3D spherical structure with evident tissue compartments and inner cavities (**Fig. 2c**), which we then aimed to characterize in more detail by immunofluorescence (**Video S1**). Strikingly, human SEM did not only express the respective lineage markers, but also established the structures morphologically characteristic of *in utero* implanted embryos (**Fig. 2d, f**). From the beginning of the *ex utero* culture, SEMs became surrounded by the Trophoblast (Tb) compartment, marked by GATA3, CK7, and SDC1 that marks syncytiotrophoblast (STB) (Days 3, 4; **Fig. 2d, left; Extended Data Fig. 12a – c**), which is reminiscent of human early *in utero* development by 8 dpf (day post-fertilization) or Carnegie Stage 5 (CS5) (**Fig. 2a, left**). At this stage, the implanting embryo starts to become surrounded by a layer of syncytiotrophoblast (STB), which directly invades maternal endometrium and supports future histotrophic nutrition of the embryo. Notably, the co-aggregation protocol devised herein does not result in a blastocoel cavity formation nor in a condensed ICM-like structure, as opposed to human and mouse blastoid models (**Extended Data Fig. 13a-h**), indicating that human SEMs do not go through a blastocyst-like stage, in agreement with observations in mouse SEM protocol (**Extended Data Fig. 13i-j**) ^27, 79, 84, 85^. The efficiency of correctly organized human SEM was estimated at day 6 to be 2.9% of the starting aggregates as judged by co-immunofluorescence for lineage markers and meeting morphology criteria (see Methods) (**Fig. 2e, Extended Data fig 12c-d**). We note that the human SEMs show a large degree of asynchrony, with up to 2 days difference in developmental staging for SEMs found at day 6-8, leading to some more advanced and some earlier structures when evaluating SEMs at a specific time point. Starting the human SEM protocol with human primed, rather than naïve, PSCs did not generate equivalent SEMs (**Extended Data Fig. 9d**), as also seen in mouse models ^27, 86^.

In human and non-human primates (NHP), the inner cell mass segregates into two lineages, the epiblast (Epi), formed by columnar cells expressing OCT4 ^5–7, 87^, and the hypoblast located underneath the Epi (Hyp, comprised of cuboidal cells expressing SOX17, GATA6, GATA4, PDGFRa) ^88–90^ (CS4-5a, **Fig. 2a**). Importantly, in our human SEM, Epi and PrE cells segregated into two distinct compartments, differentially expressing the respective lineage marker genes (OCT4 and SOX17) (**Fig. 2d**). Both Epi and Hyp were surrounded by the Tb, marked by CK7 starting from day 3 (**Fig. 2d, Extended Data Fig. 12a**), and SEMs drastically advanced morphologically by day 6 (**Extended Data Fig. 12b**). In particular, the Epi initiated formation of the amniotic cavity (AC), whilst the hypoblast formed a yolk sac (YS) cavity, establishing a bilaminar structure in between (**Fig. 2d, middle; Extended Data Fig. 12b**) reminiscent of the 9-10 dpf human embryo (CS5b, **Fig. 2a**).

The early post-implantation pre-gastrulating human embryo already contains some extraembryonic mesoderm (ExEM) ^41–47, 91^. The ExEM contributes to all fetal membranes and participates in formation of blood and placental vasculature ^91, 92^. ExEM tissue becomes abundant between the primary yolk sac (PYS) and the Tb by CS5a, forming a chorionic cavity (ChC) underneath the PYS by CS5c (11 dpf, **Fig. 2d – middle scheme**). The latter was also observed upon close examination of day 6 SEMs, revealing the presence of a cavity formed by an additional tissue layer between the YS and the Tb, that corresponds to ChC, similar to what has been characterized in natural human embryos corresponding to these stages (**Fig. 2c, d**). Later on, the ExEM expands allowing the remodeling of the PYS into becoming what is termed the secondary yolk sac (SYS) and formation of a connective stalk (St), the structure that crosses through the chorionic cavity and holds the bilaminar disc to the chorion and later forms the umbilical cord (Florian, 1930). In human SEMs, we also observed 3D expansion of all the above mentioned lumina and growth of the extraembryonic tissues (**Fig. 2c-d, f**), suggesting differentiation and remodeling of the PYS into SYS alongside ExEM expansion, and formation of a connective stalk (**Fig. 2d, f**; **Video S1; Extended Data Fig. 14**).

By 11-12 dpf (CS5c) in human embryos, the embryonic disk segregates into a ventral pseudostratified epiblast and a dorsal squamous amnion (Am) (**Fig. 2a, right panel**). The bilaminar epiblast will give rise to the embryo proper, whereas the amnion will comprise the protective membrane surrounding the fetus until birth ^94^. Strikingly, starting from day 6 and until day 8 of the *ex utero* culture (D6 – D8 SEMs), the dorsal segment of the epiblast acquired squamous morphology resembling the amnion-like layer, and the bilaminar structure adapted a disk shape (**Fig. 2d, f; Extended Data Fig. 14; Video S1**), resembling the key hallmark of the *in utero* human post-implantation development in preparation for gastrulation. Altogether, our approach exploits the developmental plasticity of genetically unmodified transgene-free human naïve PSCs, demonstrating their capacity to self-assemble into early post-implantation human embryo models that comprise both embryonic and extraembryonic compartments, as we characterize and validate further below.

### Human SEMs undergo epiblast morphogenesis and form a bilaminar disk, amnion and PGCs

We then aimed to characterize development of key lineages in SEMs in more detail. As the epiblast is derived from initially naïve PSCs, we checked whether they undergo priming after co-aggregation with other lineages and being placed in N2B27 basal conditions. Indeed, shortly after co-aggregation, we noted loss of expression of naïve marker genes (DNMT3 and STELLA) that were originally expressed in the starting naïve PSCs in HENSM conditions (**Fig. 3a**), and upregulation of the primed pluripotency marker OTX2 by day 3, alongside maintaining the expression of OCT4 and SOX2 pluripotency factors that mark both naïve and primed pluripotency states (**Fig. 3a, b**) ^95, 96^. Consistent with developmental progression, by day 4, PSCs formed an evident epiblast tissue inside the SEM which considerably grew during subsequent development (**Fig. 3c**, **Video S2**). Electron micrographs of the *in utero* rhesus monkey embryos indicate Epi polarization approximately at the time of implantation ^97^. *In vitro* attached and cultured (IVC) human blastocysts showed Epi polarization and lumenogenesis (evident by apical Podocalaxyn localization) starting from 10 dpf ^5^, while in our SEM, the Epi established apicobasal polarity from day 6, as judged by the apical localization of phosphorylated Erzin, Podocalaxyn (**Fig. 3e**), and aPKC (**Extended Data Fig. 15a**), as well as alignment of the Epi cells towards the emerging cavity (**Extended Data Fig. 15b**). The timing of lumenogenesis at day 6 also corresponded to a significant increase in the Epi cell number (**Fig. 3d**), in agreement with the histological descriptions of the *in utero* human embryo ^82^.

**Figure 3.**
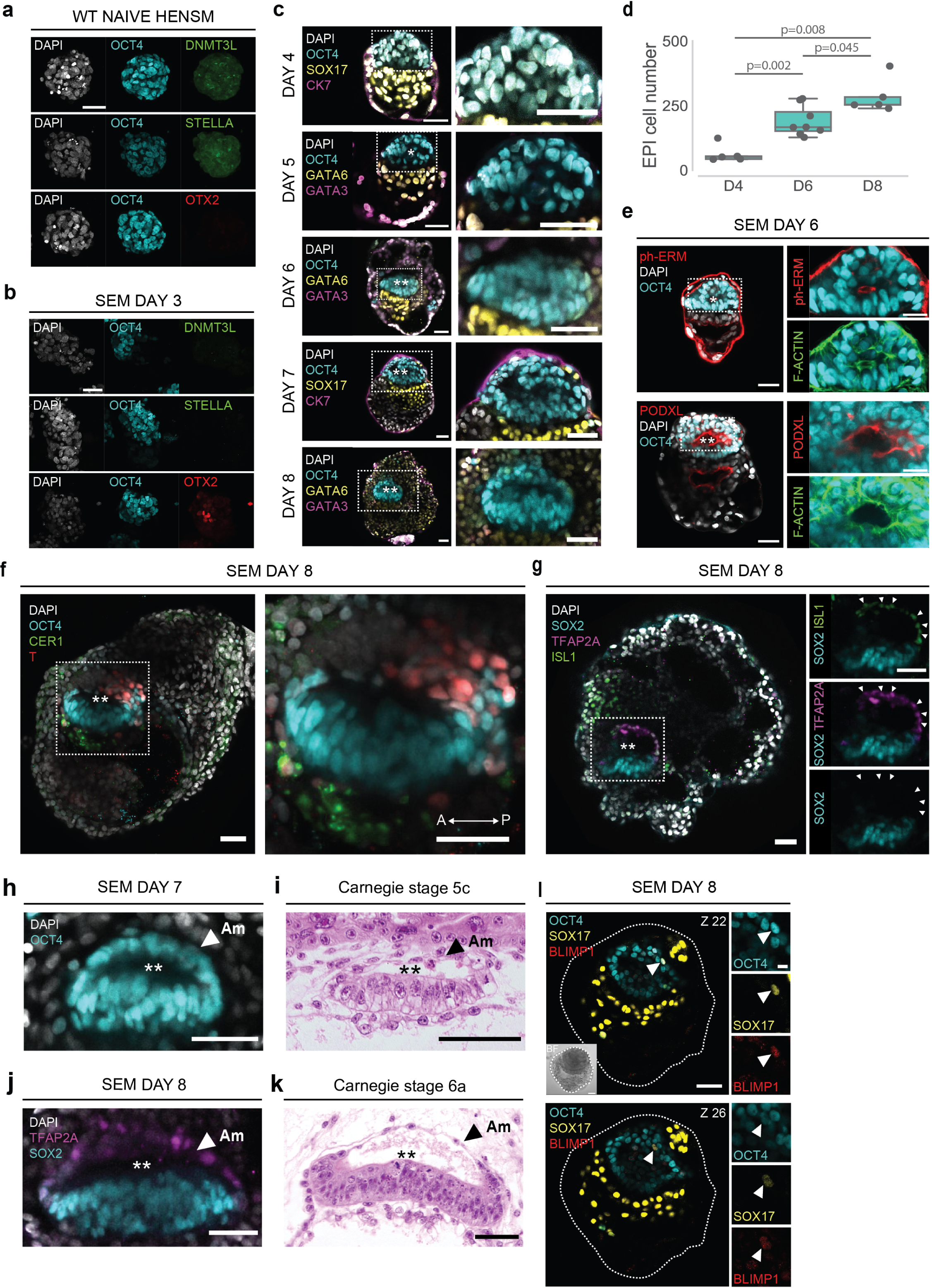
Human SEMs undergo epiblast morphogenesis and form a bilaminar disk with amnion and PGCs. **a,** immunofluorescence images of naïve PSCs in HENSM medium, showing expression of OCT4 (cyan), DNMT3L (green, top), STELLA (green, middle), but not OTX2 (red). **b**, immunofluorescence images of Day 3 SEMs in N2B27 showing expression of OCT4 (cyan) and OTX2 (red), but not DNMT3L (green, top) or STELLA (green, middle). **c**, immunofluorescence images of SEMs developing from day 4 to day 8 showing epiblast (OCT4, cyan), hypoblast (SOX17 or GATA6, yellow), and trophoblast (CK7, SDC1 or GATA3, magenta). Right, zoom into the epiblast. Single- and double-asterisks mark proamniotic and amniotic cavities, respectively. **d**, quantification of epiblast growth by counting cell numbers in day 4, 6, and 8 SEMs. p-values, two-sided Mann-Whitney U-test. **e**, immunofluorescence images of day 6 SEMs showing apical cell markers: phospho-Ezrin/Radixin/Moesin (ph-ERM, red; top), Podocalaxyn (PODXL, red; bottom), and F-actin (green). **f**, immunofluorescence image, showing Anterior-Posterior (A-P) axis, marked by expression of T/BRA (red) and CER1 (green) on the opposite sides of the SEM in day 8. Right, zoom into the epiblast (OCT4, cyan). **g**, immunofluorescence image, showing amnion, marked by co-expression of TFAP2A (magenta) and ISL1(green) in SOX2-negative cells (cyan) of the day 8 SEM. Right, zoom into the epiblast region; arrows point at the amnion cells. **h**, immunofluorescence image, showing putative amnion in day 7 SEM, characterized by squamous OCT4-positive cells; OCT4 (cyan), DAPI (white). **i**, histological sections from the Carnegie collection of the human embryo at Carnegie Stage (CS) CS5c highlighting the start of amnion (Am) formation. **j**, immunofluorescence image, showing amnion in day 8 SEMs, characterized by squamous cells expressing TFAP2A (magenta), but not SOX2 (cyan); nuclei (DAPI, white). See also **Video S2. k**, histological sections taken from the Carnegie collection of the human embryo at Carnegie Stage (CS) CS6 highlighting amnion (Am) located dorsally to the bilaminar disk. **l**, immunofluorescence images (Z slices 22 and 26, top and bottom, respectively) of day 8 SEM, showing progenitor germ cells (PGCs) with co-expression of OCT4 (cyan), SOX17 (yellow), and BLIMP1 (red). Right, zoom into the individual PGCs, marked with arrows. Inset, brightfield image, embryo perimeter is outlined. Single- and double-asterisks mark proamniotic and amniotic cavities, respectively. Scale bars, 50 µm, 25 µm (e, right), 10 µm (l, right).

Early emergence of the anterior-posterior (AP) axis is prevalent in mammals when a proportion of Epi cells initiates expression of BRACHYURY (T) at the prospective posterior side of the embryonic disk ^98^. We also checked for the expression of BRACHYURY (T) in human SEMs and identified a T/BRACHYURY-positive population of Epi cells in the posterior part of the SEM epiblast (**Fig. 3f, Extended Data Fig 15d**). The emergence of the anterior visceral endoderm (AVE), which constitutes the anterior signaling center for epiblast patterning, was seen by the expression of CER1 in the Epi-adjacent part of the VE from day 6 (**Extended Data Fig. 15c, d**). The previously known vesicular staining pattern of CER1 ^99, 100^ was evident in human SEMs (**Extended Data Fig. 15c**). Co-immunostaining for these markers further supported establishment of the AP axis and symmetry breaking by day 8 in human SEMs, where T/BRA-positive Epi cells were found in the region opposite to Cer1-positive AVE (**Fig. 3f**).

From day 6 on, the Epi exhibited patterning of early amniotic sac formation with a dorsal squamous cell population corresponding to the putative amnion, and ventral columnar pseudostratified epiblast cells (**Extended Data Fig. 15e, f**). Co-immunostaining for TFAP2A and ISL1 in day 8 SEMs, revealed their co-localization in multiple squamous dorsal cells, also depleted of SOX2 expression, confirming their amnion identity by localization, morphology, and gene expression (**Fig. 3g; Video S3**) ^101–104^. Based on the cell morphology and localization, amnion can be distinguished from the rest of the Epi between 10 and 12 dpf of human *in utero* development ^82^. Moreover, by day 8 in SEM, the Epi acquired an apparent disk shape with an enlarged amniotic cavity, while the amnion formed a thinner squamous shaped epithelium highly resembling *in utero* embryo morphology as documented in Carnegie collection at CS6a (**Fig. 3h – k, Video S3**). Lastly, we asked whether advanced *in utero*-like development of the embryonic disk would also lead to induction of early primordial germ cells (PGCs) ^105, 106^, as was seen in mouse SEMs derived from naïve PSCs ^27^. Co-immunostaining for several PGC markers identified a rare population of cells positive for OCT4, SOX17, and BLIMP1 in the dorsal Epi region of the SEM at day 8 (**Fig. 3l; Extended Data Fig. 15g**), which unequivocally constitutes a specific signature of human PGCs ^107–109^.

### Human SEMs recapitulate development of yolk sac and scaffolding of the chorion cavity by the extra-embryonic mesoderm

The yolk sac starts forming from the hypoblast between CS4 and CS5 in human. The part of hypoblast located underneath the epiblast (forming a characteristic bilaminar disc), belongs to the visceral endoderm (VE) and is a dynamic signaling center for epiblast patterning. It is connected to the parietal endoderm (PE), forming the inner cavity, primary yolk sac (PYS), which becomes reorganized during development ^42, 46^. Eventually, yolk sac serves multiple functions for the growing embryo, suppling nutrients, and the blood cell progenitors during the embryonic period until the placenta takes over ^42^. Formation of the yolk sac cavity was frequently seen in structures with all segregated lineages and became more prominent at day 6 of the *ex utero* development (**Fig. 4**). Once formed, SOX17-positive yolk sac was comprised of the columnal VE in the Epi proximity, and the squamous PE lining the opposing side of the cavity (**Fig. 4a – c**), recapitulating VE and PE cell morphology in PYS of CS5c natural embryos (**Fig. 4c**). Both VE and PE acquired apicobasal polarity, as judged by the apical localization of aPKC (**Extended Data Fig. 16a; Video S4**), which agrees with the electron micrographs of the hypoblast cell morphology in implantation stage rhesus monkey embryos ^97^. Consistent with polarity, cell shape, and localization in natural embryos, SOX17-positive hypoblast also co-expressed primitive endoderm markers GATA6 and GATA4 (**Fig. 4d, e; Extended Data Fig. 16d**).

**Figure 4.**
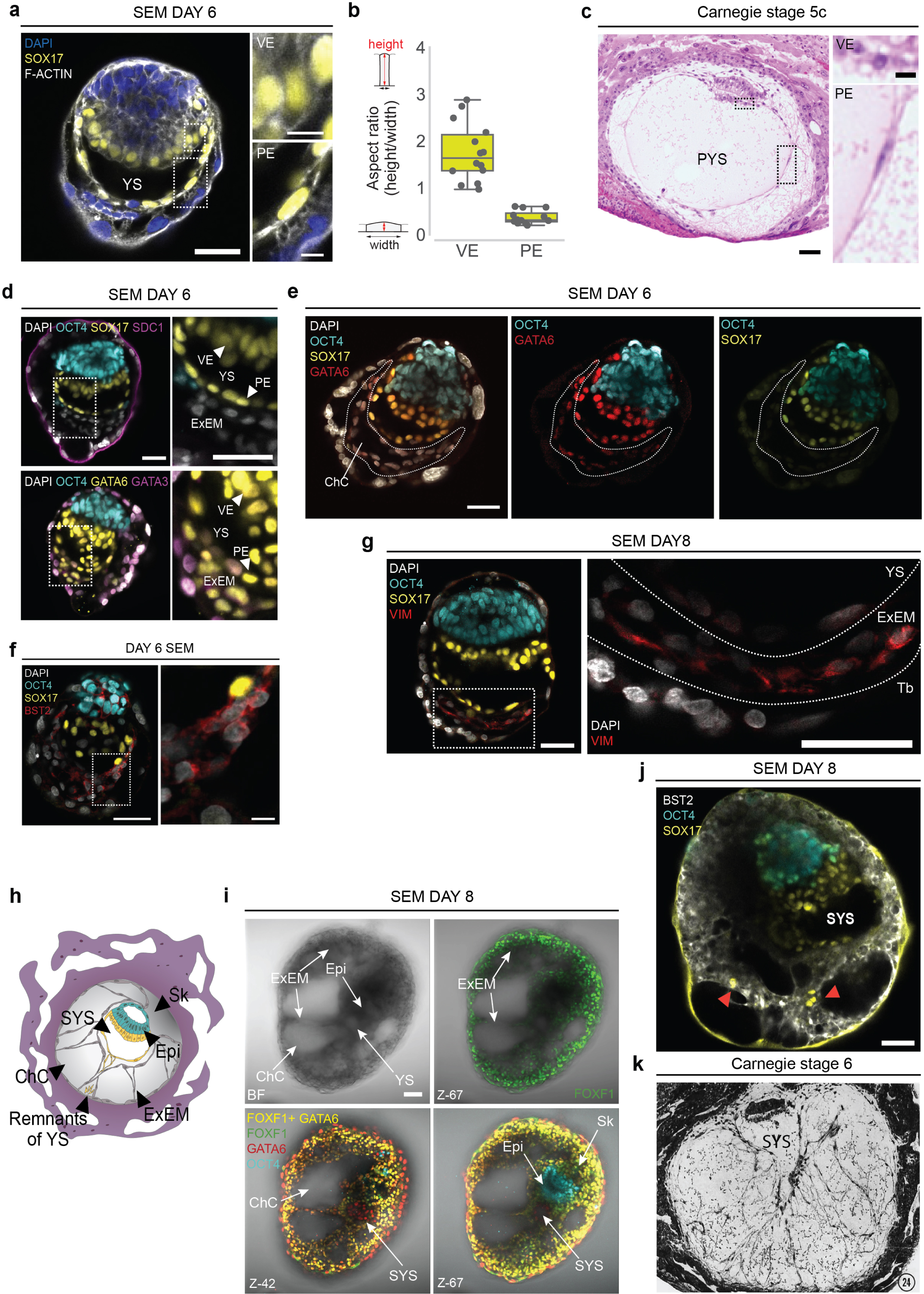
Human SEMs recapitulate development of yolk sac lumenogenesis and extra-embryonic mesoderm scaffolding. **a**, immunofluorescence image, showing yolk sac (YS) parietal (PE) and visceral (VE) endoderm (SOX17, yellow) in day 6 SEM. Nuclei (DAPI, blue), F-actin (white). Right, zoom into cuboidal VE and squamous PE cells. See also **Video S4. b**, quantification of cell aspect ratio (height/width) in VE (n = 14) and PE (n = 12) cells. **c**, Histological section taken from the Carnegie collection of the human embryo at Carnegie Stage 5c with a primitive yolk sac (PYS), cuboidal visceral endoderm (VE, top right), and squamous parietal endoderm (PE, bottom right). **d**, immunofluorescence images of day 6 SEMs showing epiblast (OCT4, cyan), hypoblast (SOX17, GATA6, yellow), and trophoblast (CK7, GATA3, magenta); nuclei (DAPI, white). Right, 2x zoom into the yolk sac (YS) region with extraembryonic mesoderm (ExEM). White arrows point at the VE and PE. **e**, immunofluorescence image of day 6 SEM showing formation of chorionic cavity (ChC) surrounded by ExEM (outlined) expressing GATA6 (red) and not SOX17 (yellow); nuclei (DAPI, white). Note lower expression levels of GATA6 in ExEM relative to YS cells. **f**, immunofluorescence image of day 6 SEM showing expression of BST2 in the cell membrane (red) in ExEM underneath the YS (SOX17, yellow); OCT4 (cyan), nuclei (DAPI, white). Right, zoom into the ExEM region. **g**, immunofluorescence image of day 8 SEM showing expression of VIM (red) in ExEM underneath the YS (SOX17, yellow); OCT4 (cyan), nuclei (DAPI, white). Right, zoom into the ExEM region. The boundaries between the YS, ExEM, and trophoblast (Tb) are outlined. See also **Video S5**. **h**, schematic of the 14 dpf human embryo showing ExEM in grey; Am, amnion; Sk, stalk; SYS, secondary yolk sac. **i**, brightfield and immunofluorescence images of day 8 SEM in different Z slices (Z 42 and 67) showing FOXF1 (green), GATA6 (red), and OCT4 (cyan). Arrows point to epiblast (Epi), ChC, ExEM, YS, Sk, and a secondary yolk sac (SYS); scale bar, 100 µm. **j**, immunofluorescence image of day 8 SEM with a cavitated ExEM marked by BTS2 (white), and SYS; OCT4 (cyan), SOX17 (yellow). Red arrows point at the SOX17+ PYS cell remnants. See also sequential sections from the same SEM in **Extended Data Fig. 17**. **k**, histological section taken from the Carnegie collection of the human embryo at Carnegie Stage 6 showing filamentous ExEM and the secondary yolk sac (SYS). Scale bars, 50 µm, 10 µm (zoom: a, c, f).

Next, we aimed to characterize the above mentioned OCT4-negative and SOX17-negative cell population beneath the yolk sac (**Fig. 2d**). Interestingly, these cells had mesenchymal rather than epithelial morphology, extending cell protrusions towards the surrounding tissues and forming an intermediate mesh-like 3D structure (**Extended Data Fig. 16b**). We then checked the expression of multiple lineage-specific transcription factors, differentially marking ExEM vs. YS during the relevant stages in marmoset embryos ^90^. The ExEM cells expressed GATA6 and GATA4, but not SOX17 (**Fig. 4e; Extended Data Fig. 16d, e**), consistent with the ExEM expression profile from marmoset embryos ^90^ (**Extended Data Fig. 16c**) and human PSC in vitro derived ExEM cells ^50^. The observed lower GATA6 level in ExEM cells compared to PrE cells (**Fig. 4e, Extended Data Fig. 16d**), is also in agreement with the expression for those lineages in the *in utero* marmoset embryos ^90^ (**Extended Data Fig. 16c**). Immunostaining for additional mesenchymal markers, such as BST2 ^50^, VIM ^87, 110^ and FOXF1 ^62^, which allow distinguishing ExEM from PrE, further validated the ExEM Identity of these cells in day 6 – 8 SEMs (**Fig. 4f-g, Extended Data Fig. 16e-f; Video S5**). Hence, we concluded that the inner cavity of the ExEM represents the chorionic cavity, which is formed by remodeling of a mesh like population of mesenchymal cells predominantly visible starting from day 6 SEMs that corresponds to the human extraembryonic mesoderm.

Histological descriptions of human and NHP embryos suggest the remodeling of the PYS cavity by the ExEM expansion with a subsequent pinching off and vesiculation of the PYS remnants, resulting in the formation of the secondary yolk sac (SYS) ^46^. In some of the day 8 SEMs, BST2 and FOXF1 positive cells formed a complex filamentous meshwork with multiple cavities, contributing to the SEM complex architecture (**Fig. 4h, i; Extended Data Fig. 17**). Closer examination of these structures revealed clusters of SOX17 positive cells entrapped between ExEM cells, suggesting residual PYS cells after the tissue remodeling (**Fig. 4j-k; Extended Data Fig. 17**). Moreover, an ExEM cell population connecting the bilaminar disk to the trophectoderm could be seen in day 8 SEMs, resembling a connective stalk structure (**Fig. 4h-I**, **Fig. 2d, Extended Data Fig. 14**), which contributes to the future umbilical cord in natural human embryos ^93^.

### Trophoblast integration and maturation in human SEMs

*In utero*, the human embryo develops surrounded by the trophoblast, which is essential for truly integrated experimental models of early implantation-stage development. The early days of trophoblast development during implantation are partially characterized in human *in vivo*. The fusion of trophoblast leads to formation of an outer syncytium, morphologically seen shortly after implantation on the earliest Carnegie data. Immunofluorescence analysis shows that the majority of human SEMs generated with our protocol are surrounded by the trophoblast with very high efficiency (79-84%), expressing multiple trophoblast marker genes, such as GATA3, CK7, and SDC1, the latter being characteristic for syncytium (**Fig. 5a; Extended Data Fig. 18a-d**) ^36^. Marker gene expression and cell morphology further indicated that the outer trophoblast layer is formed by syncytiotrophoblast, confirming the development of post-implantation trophoblast in human SEMs (**Fig. 5a; Extended Data Fig. 18a-d**). Notably, SDC1 is not expressed on the starting TE cells upon BAP(J) induction prior to the aggregation, indicating that maturation of the TE cells occurs in the aggregates (**Extended Data Fig. 7e**).

**Figure 5.**
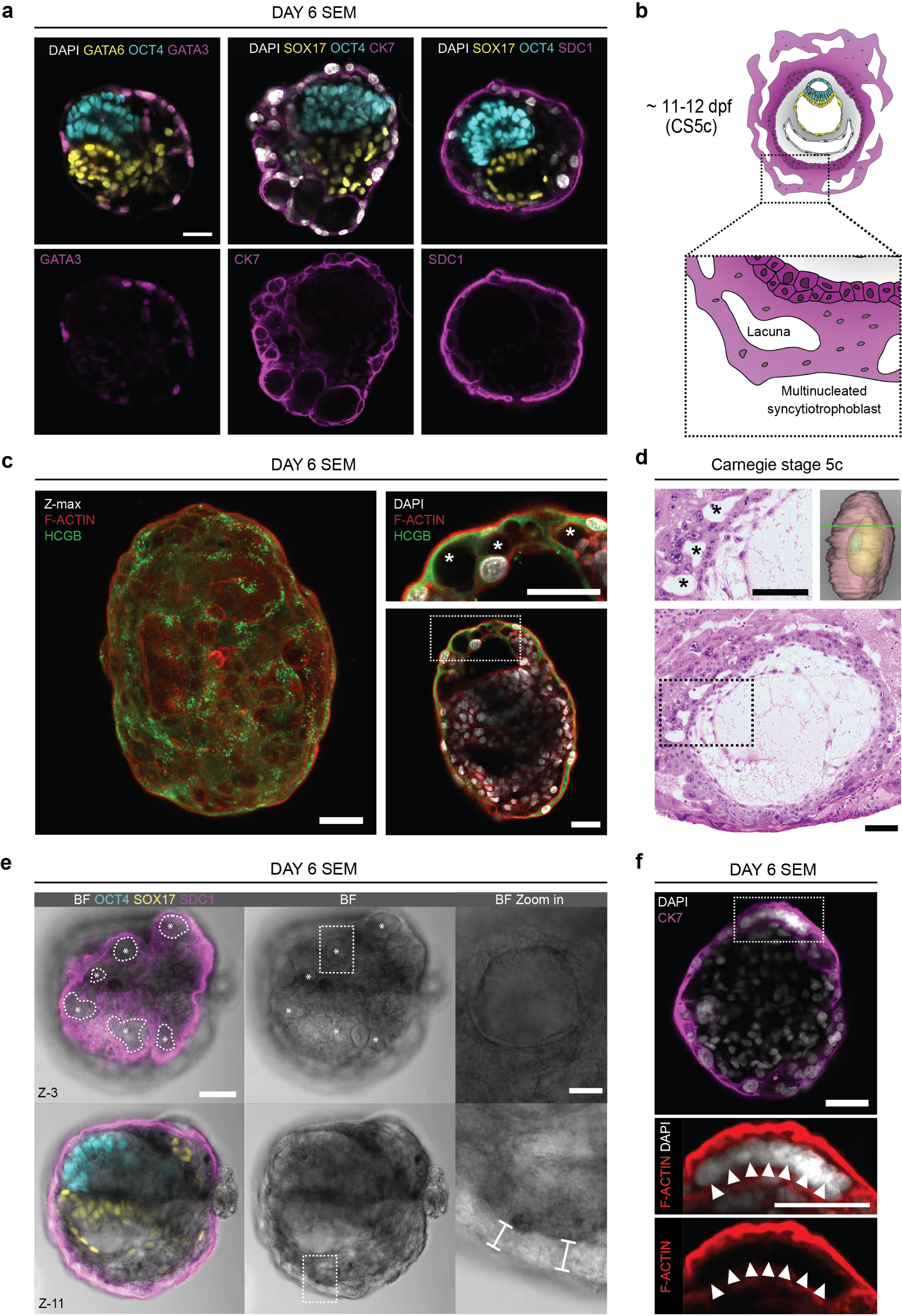
Trophoblast compartment integration and maturation in human SEMs. **a,** top, immunofluorescence images of day 6 SEMs showing epiblast (OCT4, cyan), hypoblast (SOX17 or GATA6, yellow), and trophoblast (GATA3, CK7 or SDC1, magenta). Bottom, single-channel images of the trophoblast, surrounding the SEMs. **b**, scheme of the 11-12 dpf human embryo showing multinucleated syncytiotrophoblast (magenta) with lacunae. **c**, left, maximum intensity projection image of day 6 SEMs showing HCGB expression in the outer cells; F-ACTIN (red). Right, immunofluorescence image of the same SEM showing intracellular lacunae (marked with asterisks) inside the outer syncytiotrophoblast cell layer; nuclei (DAPI, white). **d**, histological section and 3D reconstruction (top right) from the Carnegie collection of a human embryo at Carnegie Stage (CS) 5c, showing lacunae in the syncytiotrophoblast (asterisks). **e**, brightfield and immunofluorescence images of two different Z planes (number 3 and 11, top and bottom panels, respectively) of day 6 SEMs showing epiblast (OCT4, cyan), hypoblast (SOX17, or GATA6, yellow), and trophoblast (SDC1, magenta). Top row - Lacunae are outlined and marked with asterisks. Right, zoom into the lacunae (top). Bottom row - the outer syncytiotrophoblast layer (bottom), its thickness is marked with brackets. **f**, immunofluorescence image of day 6 SEM showing CK7 (magenta), F-ACTIN (red), and nuclei (DAPI, white). Bottom, zoom into the multinucleated syncytiotrophoblast cell; arrows point at multiple nuclei inside the single cell. Scale bars, 50 µm, 10 µm (e, zoom).

The lacunar phase of trophoblast development begins around 14 dpf, when the fluid-filled spaces from within the trophoblast syncytium, merging and partitioning the trophoblast into trabeculae ^111^, later contributing to the placental villi (**Fig. 5b**). Remarkably, we observed multiple cavities with variable sizes forming inside the syncytiotrophoblast and were predominantly located at the SEM periphery (**Fig 5b-d, e top panel; Video S6**), which resembled the trophoblast lacunae on the embryo periphery in utero at CS5c (**Fig. 5d**).

Another important function of the trophoblast syncytium is production of the hormones, such as human chorionic gonadotropin beta (HCGB), which maintains multiple physiological aspects of pregnancy. The immunostaining for HCGB demonstrated abundance of the hormone protein in surrounding trophoblast cells of the SEM, enriched inside the intracellular vesicles ^36^ (**Fig. 5c; Extended Data Fig. 18e**). We additionally confirmed its secretion by detection of soluble HCG in media in which day 7 SEMs were placed for 24h until day 8 (**Extended Data Fig. 18g**). Lastly, we checked whether the syncytium in our SEMs is multinuclear, as it is typically seen during early placental development ^112^. Phase-images alongside co-immunostaining of the trophoblast marker SDC1 enabled us to see clearer the trophoblast cell shape forming a layer of thick outer syncytium (**Fig. 5e, bottom**). Co-immunostaining with F-ACTIN, that helps define individual cell membrane boundaries, and DAPI, showed that the trophoblast syncytium indeed has multiple nuclei (**Fig. 5f; Extended Data Fig. 18f**). Collectively, these findings confirm proper integration, maturation, and localization of Tb cells in our SEMs.

### scRNA-seq analysis validates human SEM cellular composition and identity

To further validate and examine the milieu of cell types present in the human SEMs generated herein in a more unbiased manner, we performed a single cell transcriptomic analysis by Chromium 10X scRNA-seq on selected day 4, 6 and 8 SEMs. We analyzed ∼4,000-8000 cells from pooled high-quality SEMs manually selected at these experimental timepoints. After quality control and strict filtering, a total of 12,190 single cells were used for subsequent analyses (**Extended Data Fig. 19a-c**). Uniform Manifold Approximation and Projection (UMAP) analysis using Seurat package identified a total of 13 separate cell clusters (**Fig. 6a**). Cluster annotation was performed based on expression (+) or lack-of expression (-) of previously defined lineage-specific markers that allowed us to annotate all 13 identified cell clusters **(Fig. 6 b,c)**: Epiblast (Epi – in total four clusters: OCT4+, SOX2+), Yolk-Sac/Hypoblast (YS/Hb – in total three clusters: SOX17+, APOA1+, LINC00261 in addition to GATA6+, GATA4+, PDGFRa+), Extra embryonic mesoderm (ExEM – in total four clusters: FOXF1+, VIM+, BST2+, in addition to GATA6+, GATA4+, PDGFRa+), amnion (Am – one cluster: ISL1+, GABRP+, VTCN1+), and trophoblast (Tb - one cluster: GATA3+, SDC1+, CPM+) (**Fig. 6a,b**).

**Figure 6.**
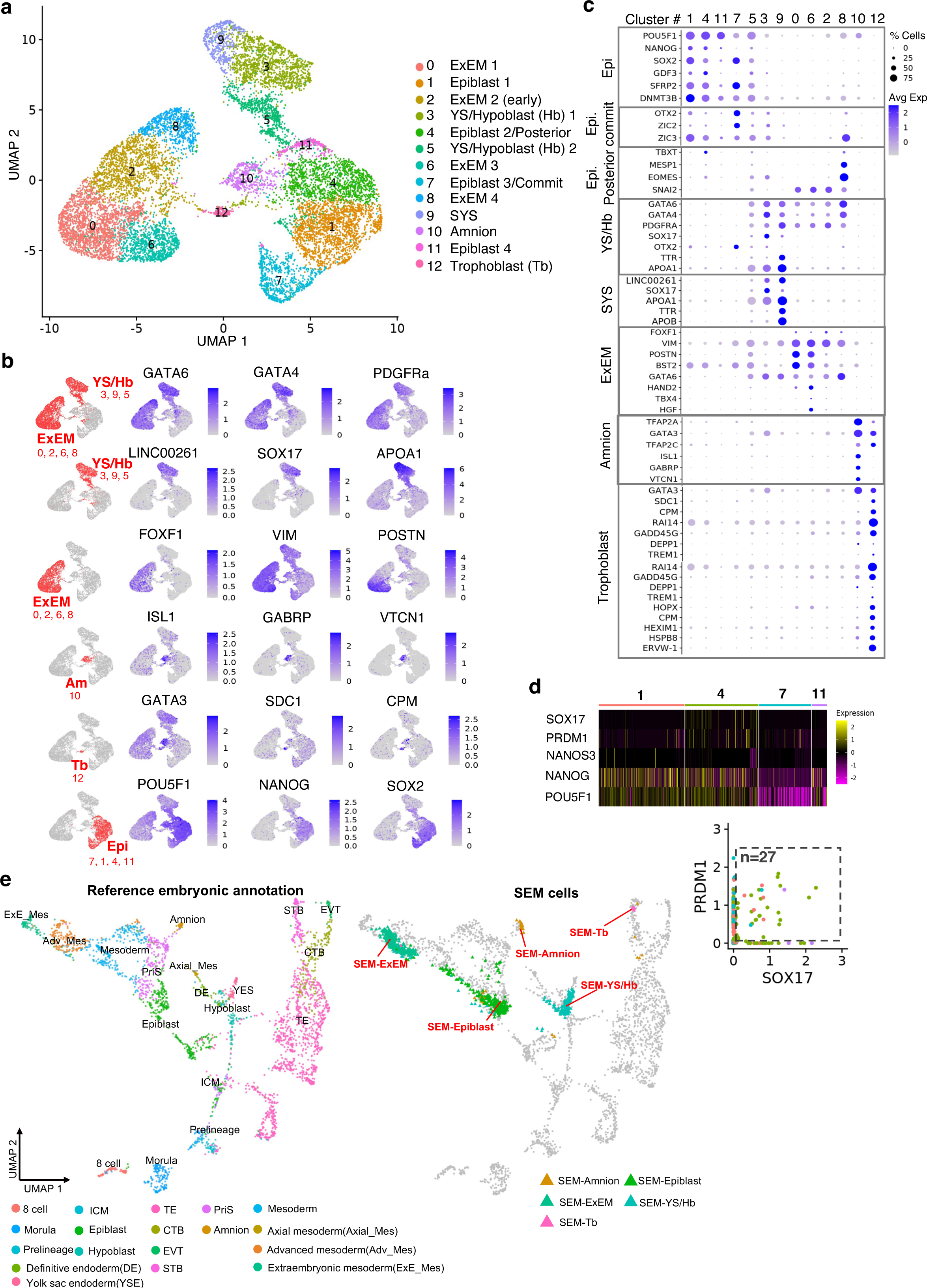
scRNA-seq analysis validates key cell type identity comprising the human SEM. **a**, UMAP plot displaying individual human SEM derived cells. Points are colored according to their assigned cell cluster. All 13 clusters representing multiple cell types were identified based on known marker genes and overlap with gene signatures, as indicated. ExEM: Extra-embryonic Mesoderm; YS: Yolk-Sac; Hb: Hypoblast; SYS: Secondary Yolk Sac; Am: Amnion; Tb: Trophoblast. **b**, Normalized expression of key marker genes projected on SEM UMAP. Cell type cluster highlighted is indicated in red, along with the corresponding cluster number. **c**, Dot-plot illustrating the expression of key marker genes across the 13 clusters. Cell types as in (A) and (B). Color shade indicates average expression. Dot size indicates the percentage of cells in the cluster which express the marker highlighted. **d**, left: POU5F1+PRDM1+SOX17+ cells (n=27) can be identified in Epi clusters, with majority located in cluster 4 (n=19). Right: Normalized expression heatmap of PGC markers (SOX17, PRDM1, NANOS3, NANOG, POU5F1). 9 cells were positive for POU5F1, PRDM1, SOX17 and NANOS3, again confirming their identity as human PGCs. **e**, Left: UMAP projection of integrated human embryonic reference ^101^ consisting of 5 human embryonic data sets spanning in vitro cultured human blastocysts ^24, 118, 119^, 3D-in vitro cultured human blastocysts until pre-gastrulation stages ^120^, and a Carnegie Stage 7 (CS7) 16-19dpf human gastrula ^117^. The color of each data point corresponds to the cell annotations retrieved from the respective publication. Right: The grey data points represent embryonic reference cells, as in the left panel. Each colored triangle represents the projection position of representing neighborhood nodes from SEM cells onto the human embryonic reference UMAP space.

From the four annotated Epi clusters (all expressing CD24 primed pluripotency marker alongside OCT4 (POU5F1) and SOX2 (**Extended Data Figure 20a-top row**), we subclassified two of them. The first one, which we termed Posterior epiblast cluster (cluster 4), was marked by upregulation of TBXT (BRACHYURY/T), co-expression of MIXL1, EOMES, MESP1 and WNT8A **(Extended Data Fig. 20a, second row)**, which are all markers of epithelial to mesenchymal transition (EMT) process, known to accompany upregulation of brachyury during peri-gastrulation in mammalian species^113^. The second Epi subcluster (cluster 7) we termed as “committed epiblast” as it was marked by upregulation of ZIC2, ZEB2 and VIM lineage commitment marker expression, and absence of NANOG while maintaining OCT4 and SOX2 pluripotency markers **(Extended Data Fig. 20a, bottom row)**. Rare PGCs (POU5F1+PRDM1+SOX17+ cells) (n=27) could be also identified within epiblast clusters, most of them (n=19) in cluster 4 **(Fig. 6d)**. The fact that PGCs did not create their own subcluster likely results from their relative scarcity within SEMs, as was observed in mouse SEMs ^76, 114^. However, these markers are definitive for human PGC identity ^115^ and confirm our immunostaining based detection results for human PGCs in SEMs (**Fig. 3l**). Although we had technical problems in maintaining high trophoblast viability and recovery following SEM enzymatic dissociation, we still observed a well separated cluster expressing lineage markers of trophoblast identity (Tb-cluster 12) **(Extended Data Fig 20b)**. While amnion cells can share certain markers with Tb (like GATA3 and TFAP2A) ^87, 101^, they formed a separate cluster which also expressed amnion markers like ISL1, GABRP and VTCN1, that are specific to amnion but not to Tb (**Extended Data Fig. 20c**) ^27^. The latter is consistent with amnion immunostaining results in human SEMs shown herein (**Fig. 3g**). It has been hypothesized that human amnion might be a significant source of BMP4, analogous to the extra embryonic ectoderm in mice that serves as a signaling secreting source of Bmp4 and Furin secretion ^87, 103, 116^. We note that in our scRNA-seq data BMP4 and FURIN are expressed in a fraction of human SEM derived amnion cells (**Extended Data Fig. 20d**).

Among the three different yolk-sac (YS) clusters that commonly expressed GATA4, GATA6, PDGFRA and APOA1 (clusters 3,5 and 9), we found that recently identified markers of secondary yolk-sac (SYS) in marmoset ^90^, TTR, APOB and GSTA1, are expressed more specifically only in cluster 9, supporting its subclassification as SYS **(Extended Data Fig. 21a**, **Fig. 6a)**. STC1, LHX1 are absent in SYS (cluster 9), but not in primary yolk-sac (clusters 3 and 5) also consistent with findings in marmoset. ^90^ Lack of uniform HHEX and GSC among most cells in all YS clusters, is consistent with primitive, rather than definitive, endoderm identity for these three YS related clusters **(Extended Data Fig. 21a)** ^52^. The co-expression of DKK1 and LHX1 detected via scRNA-seq alongside CER1 expression among some of the SOX17+ cluster 3 YS/Hb cells, supports the validity of AVE cell identity in SEMs (**Extended Data Fig. 21a – black arrows in second row)**. Among the four identified ExEM clusters, based on FOXF1 and BST2 expression (**Fig. 6b**), HAND1, LEF1 and BMP4 were also expressed consistent with mesenchymal/mesodermal like identity **(Extended Data Fig. 21b)**. However, we noted that ExEM clusters 0 and 6, but not ExEM cluster 2 and 8, expressed significantly higher levels of extra cellular matrix related genes (COL3A, COL1A1, COL1A2, COL6A2 and FIBRONECTIN1 (FN1)) **(Extended Data Fig. 21b-c)** suggesting the possibility that that ExEM clusters 2 and 8 might represent a relatively earlier and less mature ExEM lineage identity. While ExEM cluster 8 cells express CER1, DKK1 and LHX1 AVE markers, they lack other key VE/AVE markers like SOX17 and APOA1 and thus were not annotated as AVE, but rather as EXEM cells as they predominantly express mesenchymal signature (**Extended Data Fig. 21a – second row; b-first row)**. The latter is consistent with CER1,DKK1 and LHX1 being co-expressed also in extra-embryonic mesodermal cells as detected in primate ^90^ and gastrulating human embryo datasets ^117^.

We next conducted a comparison analysis of the transcriptional profile of cells in the human SEMs dataset to an integrated human embryonic reference ^101^ consisting of 5 human embryonic data sets spanning in vitro cultured human blastocysts ^24, 118, 119^, 3D-in vitro cultured human blastocysts until pre-gastrulation stages ^120^, and a Carnegie Stage 7 (CS7) 16-19dpf human gastrula ^117^ (**Fig. 6e**). UMAP projection confirmed the annotated identity of the SEM clusters and validated their resemblance to the transcriptome and cell type composition of early post-implanted human embryos, including those derived following *in vitro* attached growth (**Fig. 6e; Extended Data Fig. 19d, e**). Projection of SEMs cells onto the human embryonic reference further highlighted that the transcriptional profile of cells differed from human pre-implantation embryos as expected, and all cells corresponded to a post-implanted lineage (Fig. 6e). This analysis validated the detection of post-implantation Epi, YS, ExEM and Tb, as well as emergence of other transcriptomic states, such as amnion, Hb/YS and posterior Epi (**Fig. 6e; Extended Data Fig. 19d, e**). Overall, despite of the known limitations of cross-dataset, cross-platform comparisons, the single cell transcriptomic analysis presented above supports the conclusion that human SEMs recapitulate lineage differentiation of the early human post-implantation embryo.

## Discussion

Studying early human post-implantation development is crucial for understanding the basis of normal human embryogenesis as well as developmental birth defects, since around a third of human pregnancies fail between implantation and the fourth week of gestation ^121^. Moreover, it is becoming increasingly clear that devising and optimizing protocols for 2D *in vitro* human ESC and iPSC differentiation into mature cell types, would greatly benefit from understanding the key mechanical, transcriptional, and signaling pathways active during early embryogenesis, to further improve the quality and efficiency of differentiation protocols ^122^. Such research endeavors would require large numbers of donated human embryo derived materials from early post-implantation stages but justified ethical barriers and the scarcity of such samples, make conducting the mechanistic human embryo-based research ethically and technically challenging. Given the capacity to generate stem cells from human embryonic and extraembryonic components, generating a human stem-cell derived model capable of mimicking the key developmental milestones occurring during the early post-implantation stages is becoming a necessary key element of a growing toolbox of SEMs^1, 2^.

Here, we readapted an equivalent approach to that recently described in mice ^27, 28^ and generated self-organizing human post-implantation SEMs exclusively from naive PSCs, but without the need to genetically modify or overexpress exogenous lineage factors for priming the naïve PSCs towards the three different extra-embryonic lineages prevalent at these developmental stages, contrary to what is currently still inevitable in mouse SEM derivation protocols ^27, 28^. The latter finding further underscores the self-organization capability of naïve pluripotent stem cells to generate both embryonic and all extra-embryonic compartments ^36, 50, 57^, including the ExEM found in human, but not mouse, early pre-gastrulation stages. The human SEMs generated herein mimic the 3D architecture and key developmental landmarks of *in utero* developed natural human embryos from day 7-8 to day 13-14 post-fertilization (Carnegie stage 5a-6a). We observed proper spatial allocation of cell lineages into defined embryonic and extra-embryonic compartments in the complete absence of fertilization or interaction with maternal tissues and without the need of providing external targeted signaling pathway induction during the self-organization of the aggregated cells.

The fact that human (and mouse) SEM formation bypasses the blastocyst-like stage and that the *ex vivo* protocols for growing natural human blastocysts into the authentic 14 dpf stage are still lacking, makes donated fetal materials from 7-14 dpf as the only relevant control for benchmarking the SEM describe herein. Obviously, obtaining new 7-14 dpf human embryo samples is not a viable option from both technical and ethical reasons. However, this limitation is partially mitigated by the available Carnegie and other embryo atlas collections ^82, 83^, as well as the ability to compare human development to NHP embryo models and datasets. Moreover, we emphasize that we are only trying to make a reproducible embryo <u>model</u> with recognizable recapitulation of the key milestones and the critical embryonic structures, which will be helpful for research purposes even if it does not entirely resemble the natural human embryo. At the level of structure, our human SEM resembles the situation in utero.

Naïve pluripotent stem cell growth conditions typically utilize FGF/MEK signaling inhibition, which leads to the loss of imprinting after extended passaging ^123, 124^, perturbing the developmental potential of such cells and thus might partially underlie the low efficiency yield and SEM quality. This risk may possibly be mitigated in the future by using naïve conditions with titrating down the concentration of FGF/MEK pathway inhibitors or using alternative naïve conditions that do not target this pathway^123, 125^. At this moment, the reduced efficiency and heterogeneity observed during the formation of human SEMs is a complicating factor that needs to be overcome to facilitate the use of such platforms as viable experimental set-ups. Nevertheless, the emergence of well-defined complete structures suggests that this will likely be possible. It is plausible that upon further experimentation and mechanistic understanding of self-organization, the efficiency and variability in SEM formation can be improved in the future. It is also likely that alternative *ex utero* culture platforms, aggregation strategies, or growth conditions will yield similar or enhanced results relative to the ones reported herein with human SEMs.

Testing of whether human SEMs described here can develop further towards completing gastrulation and initiating organogenesis as achieved with mouse SEMs, may be of critical experimental importance and may offer insights into previously inaccessible windows of early human development^1, 2^.

## Acknowledgments

J.H.H lab was funded by Pascal and Ilana Mantoux; Flight Attendant Medical Research Institute (FAMRI); MBZUAI-WIS Program, ISF; Minerva, Israel Cancer Research Fund (ICRF), Kimmel Stem Cell Research Center, and a sponsored research program by Renewal Bio Ltd. F.L. lab is supported by the Ming Wai Lau Center for Reparative Medicine, Ragnar Söderbergs Stiftelse, Wallenberg Academy Fellow, Center for Innovative Medicine and Karolinska Institutet SFO Stem Cells and Regenerative Medicine. S.P. lab is supported by the Swedish Research Council, Swedish Society for Medical Research. S.P. holds the Canada Research Chair in Functional Genomics of Reproduction and Development (950-233204),

## Author contributions

B.O and E.W established the SEM aggregation conditions and protocols for ex utero culture, designed and conducted most of the wet lab work and contributed on manuscript elaboration. B.O established the human stem cell conditions, and inductions. E.W and V.B conducted most embryo immunostaining and confocal imaging. V.B made light sheet microscopy analysis and helped writing the manuscript. B.O. generated cell lines with assistance from S.V and optimized the final SEM protocol. A.A.C conducted some aggregation experiments and roller culture adaptation for the system, immunostaining and microscopy and sample preparation for 10X scRNA-seq experiments. M.Y.C helped on human stem cell culture expansion and optimizations. C.Z conducted the scRNAseq comparative analysis to previous human and NHP datasets, under the supervision of F.L and S.P. S.T contributed on optimization of lineage inductions. R.C generated MEF and other critical reagents for stem cell maintenance. S.A and D.L conducted immunostainings and RT-PCR. F.R, C.J, M.R assisted in immunostainings. N.L assisted on lentivirus production and flow cytometry experiments. E.A supervised flow cytometry and sorting experiments. T.S helped in bioinformatic analysis. S.V generated plasmids. A.A.C, M.K. and H.K.S performed RNA library preparation and sequencing. S.A provided input on optimizing light sheet microscopy experimentation and analysis. N.N. – conducted and supervised bioinformatics analyses. J.H.H. conceived the idea for the project, supervised data analysis and manuscript writing.

## Extended Data Figure Legends

**Extended Data Figure 1.**
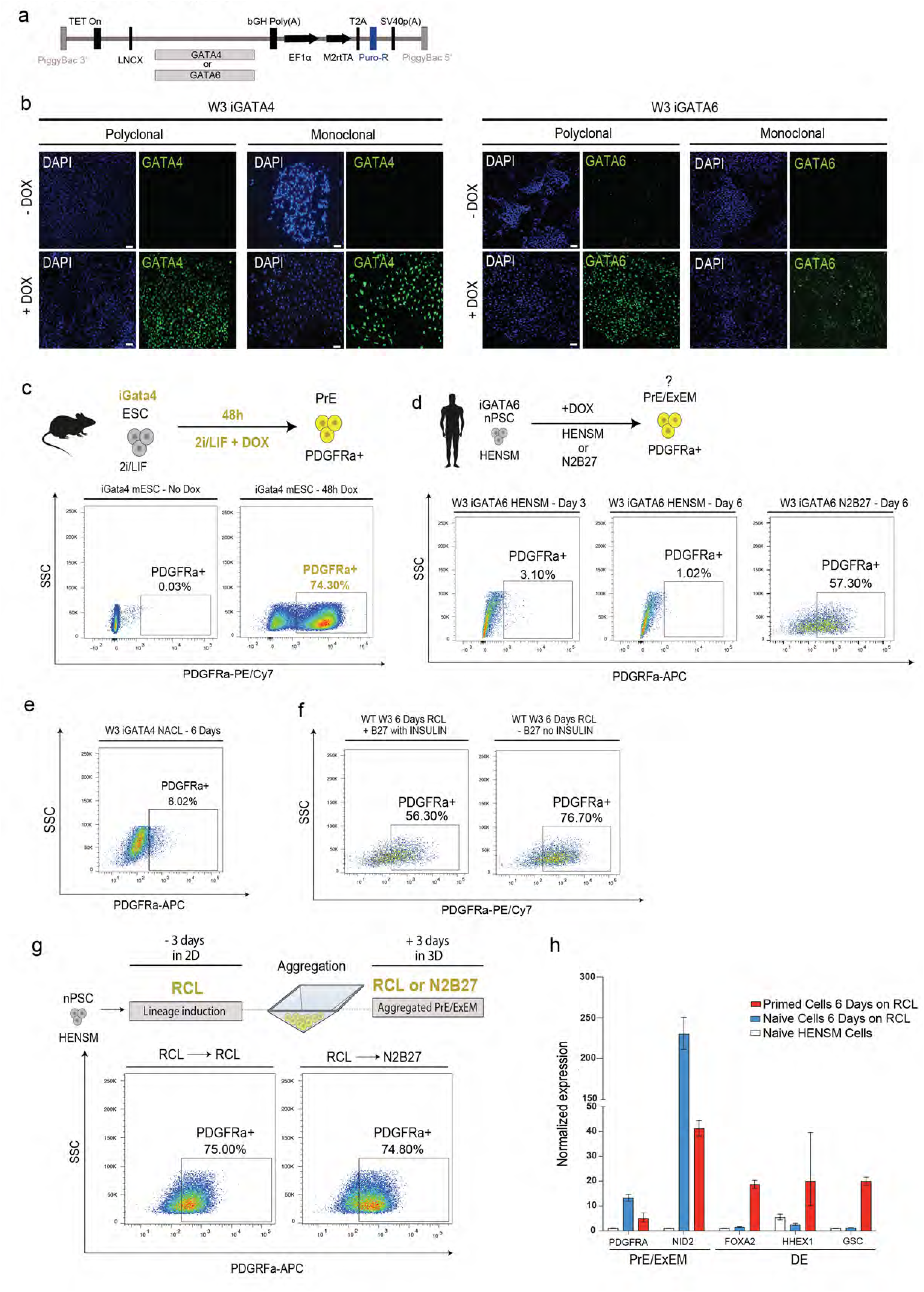
Optimization of extra embryonic lineage induction using transient overexpression of GATA4 and GATA6. **a,** scheme of the donor plasmid vector used for genomic integration of the DOX-inducible iGATA4 or iGATA6 overexpression in human PSCs. **b**, Representative immunofluorescence images of mono- and polyclonal iGATA4 (left) and iGATA6 (right) WIBR3 hESC (W3) clones, showing uniform expression of GATA4 (green) and GATA6 (green) only in response to DOX; nuclei (DAPI, blue). Scale bars, 50 µm. **c**, Flow Cytometry (FACS) plots of PDGFRa-PE/Cy7 marking primitive endoderm (PrE) induction from mouse embryonic stem cells (ESC) using iGata4 with DOX for 48 hours in 2i/Lif medium (right) versus the control condition (without DOX). **d**, FACS plots of PDGFRa-APC for putative PrE/ExEM induction from human naïve pluripotent stem cells (nPSCs) using iGATA6 with DOX in different media (N2B27 and HENSM) as indicated. **e**, Representative FACS plot of PDGFRa-APC for putative PrE/ExEM induction from iGATA4 nPSCs in NACL media. **f**, Representative FACS plots of PDGFRa-APC for putative PrE/ExEM PrE induction of wild type (WT, non-genetically modified) nPSCs in RCL media with B27 with insulin (left) or B27 without insulin (right). **g**, Representative FACS plots of PDGFRa-APC for putative PrE/ExEM induction from human nPSCs after three days in RCL medium (conventional 2D conditions) followed by three days of RCL in aggregation setting (3D -left) or N2B27 (right). **h**, qRT-PCR gene expression (normalized by GAPDH and ACTIN) of the endodermal markers for PrE/ExEM induction in RCL medium starting form naïve (blue) vs primed (red) human PSC. Naïve human PSCs maintained in HENSM (white) were used a reference control (set as 1). (DE) - definitive endoderm.

**Extended Data Figure 2.**
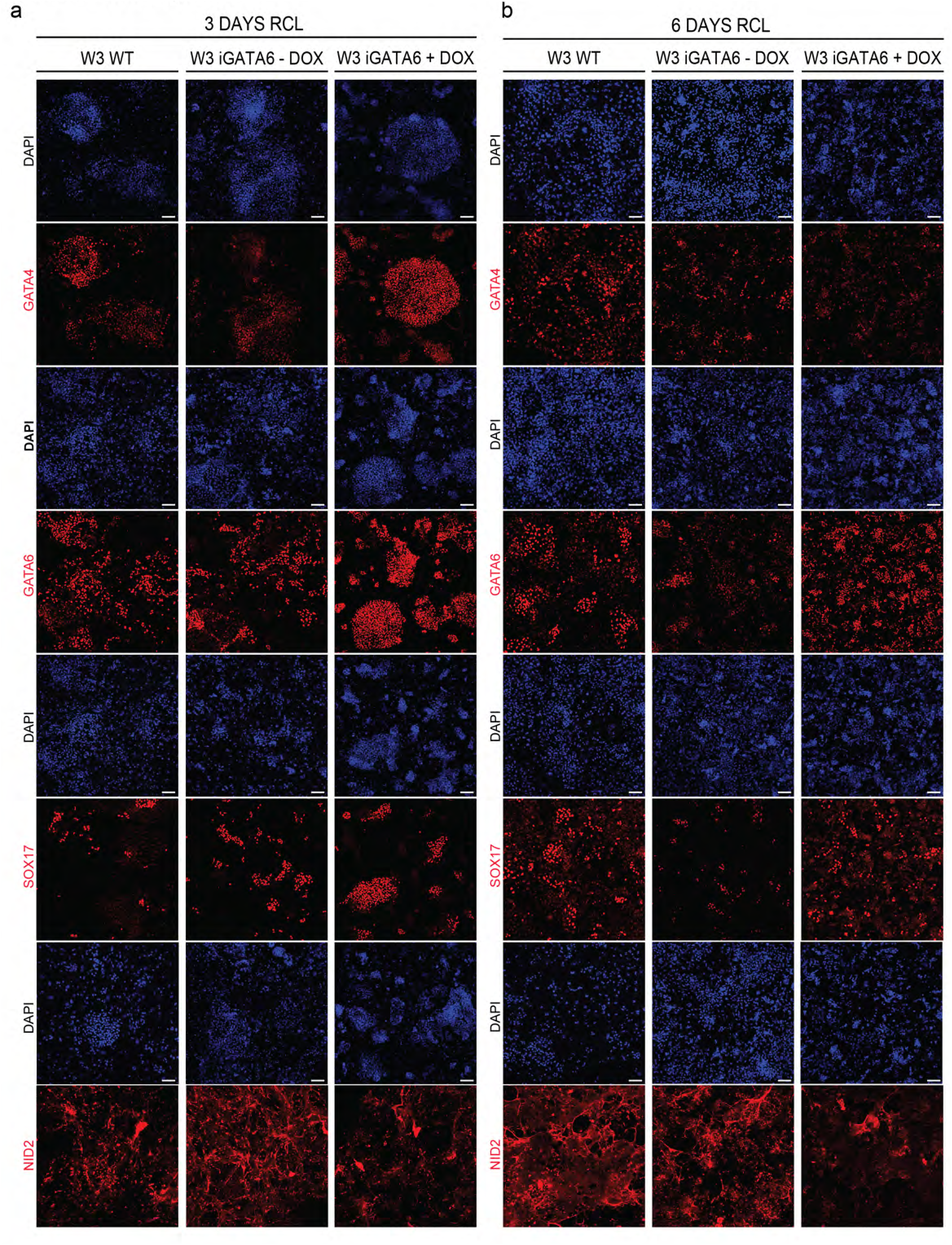
Immunostaining for PrE signature in naïve PSC derived cells in RCL conditions. **a, b,** immunofluorescence images of WT and iGATA6 naïve PCS cells with and without DOX induced in RCL media for 3 or 6 days as indicated (**a** and **b**, respectively), showing expression of the indicated PrE markers (red), including SOX17, and nuclei (DAPI, blue). Scale bars, 100 µm.

**Extended Data Figure 3.**
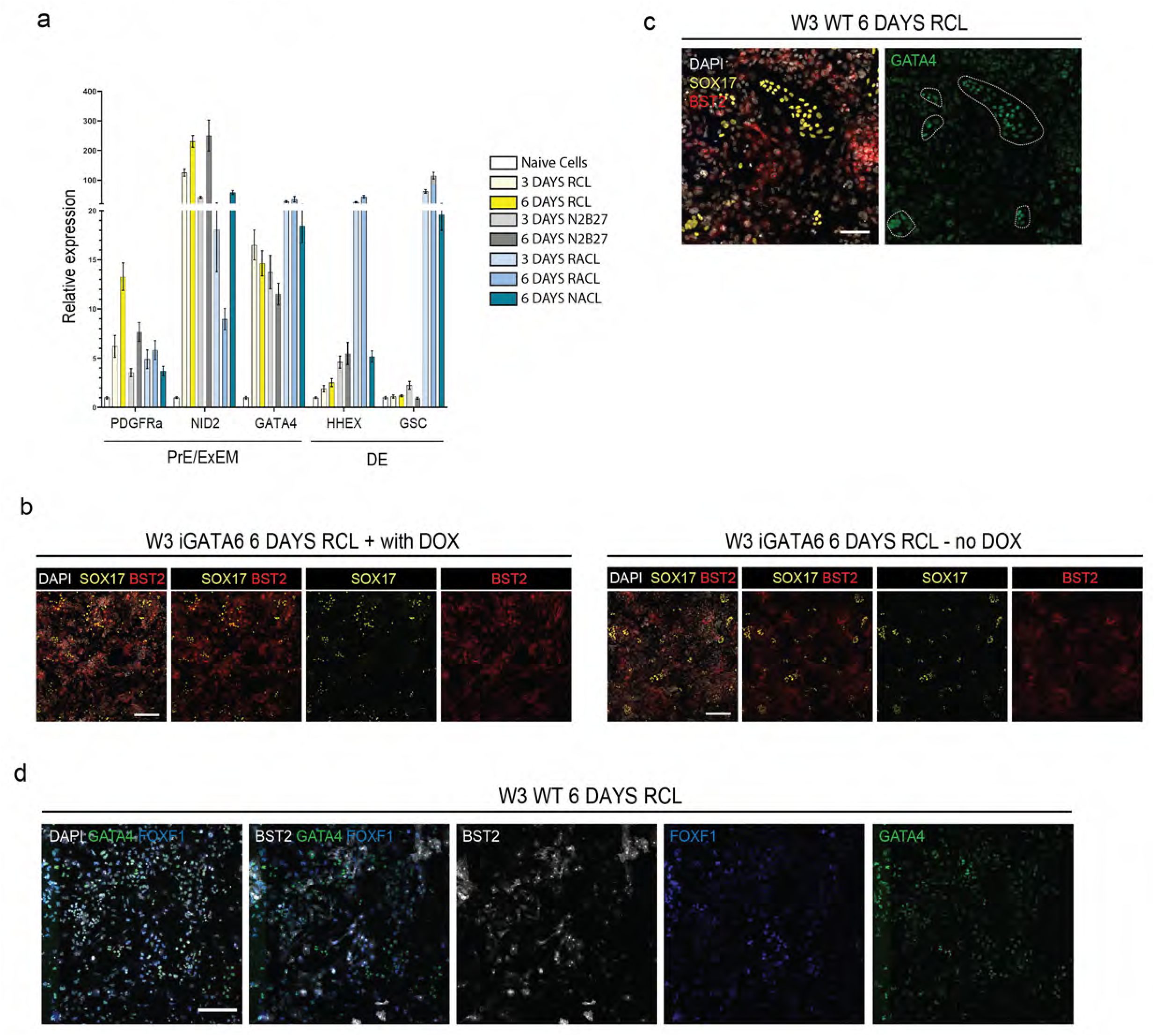
Optimization of PrE/ExEM induction protocol from human naïve PSCs. **a**, RT-qPCR gene expression (normalized by GAPDH and ACTIN) of the endodermal marker genes, in RCL (yellow), RACL (blue), NACL (dark blue), and N2B27 (grey) media conditions versus naïve PCSs used as reference control (white). PrE-/ExEM and definitive endoderm (DE)-specific genes are separately underlined. **b**, immunofluorescence images iGATA6 nPSCs induced in RCL media for 6 days (with or without DOX), showing expression of SOX17 (yellow), BST2 (red), and GATA4 (green) (see also Fig. 1e). In both set-ups, BST2-positive (ExEM) and SOX17 – positive (PrE) cell populations are mutually exclusive. Scale bars, 200 µm. **c**, immunofluorescence images of WT WIBR3 (W3) nPSCs induced in RCL media for 6 days, showing expression of GATA4 (green), FOXF1 (blue), and BST2 (white); nuclei (DAPI, white; left panel). As expected, GATA4 marks both SOX17+ PrE and BST2+ ExEM populations. Scale bars, 500 µm. **d**, left, immunofluorescence images of iGATA6 nPSCs induced with DOX in RCL media for 3 days (top) and 6 days (bottom). **d**, Immunofluorescence images of WT nPSCs in RCL media for 6 days (bottom). SOX17 (yellow), BST2 (red), nuclei (DAPI, white). Scale bars, 100 µm. Please note that the BST2+ cells are GATA4+/FOXF1+, excluding the possibility that they represent residual pluripotent cells in the RCL induced cultures.

**Extended Data Figure 4.**
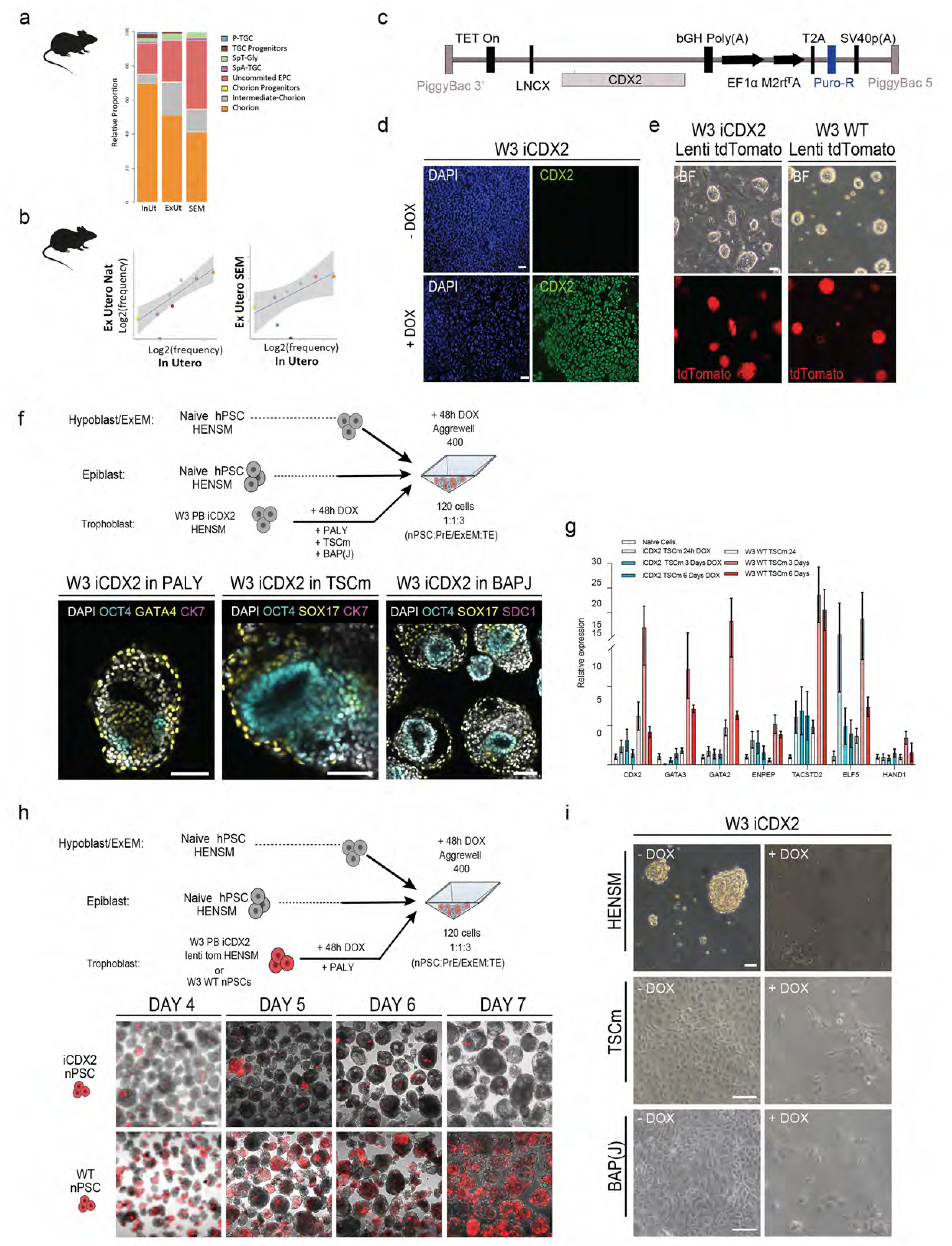
Testing Tb induction using transient overexpression of CDX2 in human naïve PSCs. **a,** relative proportion of the indicated cell types among extra-embryonic cells of mouse natural embryos grown in utero or ex utero and in day 8 mouse SEMs generated from iCdx2 mouse naïve ESCs (and not embryo derived mouse TSC lines)^76^. Three pooled samples are presented: *in utero* natural embryos (n = 2401 cells), *ex utero* natural embryos (n = 1382), and mouse iCdx2 day 8 SEMs (n = 6249). The cell types: Chorion, Intermediate-Chorion, Chorion Progenitors, Uncommitted Ectoplacental-Cone Cells (EPC), Trophoblast Giant Cells (TGC) progenitors, parietal trophoblast giant cells (pTGC), spiral artery associated trophoblast giant cells (SpA-TGC), and junctional zone spongiotrophoblast cells (SpT-Gly) based on previously published similar analysis and annotations ^77^. **b**, frequencies of the cell types presented in **(a)**, showing a significant reduction of TGC-progenitors and pTGCs in *ex utero* embryos (natural and SEM), compared to *in utero* embryos. Shaded area represents 95% confidence interval. This analysis confirmed that naïve PSCs derived TSC lineage following Cdx2 overexpression under optimized conditions developed in ^76^, can contribute to both the chorionic and ectoplacental cone lineages in mouse SEMs generated exclusively from mouse naïve PSCs. **c,** scheme of the donor plasmid vector used for genomic integration of the DOX-inducible CDX2 overexpression in human PSCs. **d**, immunofluorescence images of iCDX2 cells, showing CDX2 (green) expression only in response to DOX; nuclei (DAPI, blue). Scale bars, 50 µm. **e**, images of iCDX2 (left) and WT (right) human W3 ESCs in HENSM conditions, showing live fluorescence of tdTomato (red) after transfection with lentiviral particles carrying the fluorophore. Scale bars, 50 µm. **f**, top, the scheme of the experiment, where naïve PSCs and Hypoblast/ExEM induced cells were aggregated with iCDX2 cells, induced by DOX for 48h in three different media, PALY (N2B27 supplemented with **P**D0325901, **A**83-01, h**L**IF, and **Y**-27632) ^22^, TSCm ^70^, and BAP(J) (DMEM/F12 based medium with ALK4/5/7 inhibitor A83-01, FGF2 inhibitor **P**D0325901, and BMP4 substituted with JAK inhibitor I after 24h). **f**, bottom, immunofluorescence images showing no surrounding trophoblast in aggregates with iCDX2 cells, regardless of the media conditions; epiblast (OCT4, cyan), hypoblast (GATA4, SOX17, yellow), trophoblast (CK7, SDC1, magenta), nuclei (DAPI, blue). Scale bars (from left to right), 100 µm, 100 µm, 50 µm. **g**, RT-qPCR gene expression (normalized by GAPDH and ACTIN) of iCDX2 (blue) and WT (red) cells in TSCm conditions for one, three, and six days versus naïve PSCs maintained in HENSM conditions and used as a reference control (white). **h**, top, the scheme of the experiment, in which naïve PSCs and Hypoblast/ExEM induced cells were aggregated with WT or iCDX2 tdTomato-labelled nPSC induced towards trophectoderm (TE) in PALY media with or without DOX as indicated. **h,** bottom, brightfield and live fluorescence images of day 4 – 7 aggregates showing localization of the trophoblast. Scale bar, 200 µm. **i**, Phase contrast microscopy images showing iCDX2 cells induced for 72h in different media (HENSM, BAP(J), TSCm) with or without DOX, showing reduced viability upon iCDX2 transgene overexpression. In all conditions, 750,000 iCDX2 cells were seeded in 10 cm Matrigel coated plates, and DOX induction was started 24h after seeding. Scale bar, 100 µm.

**Extended Data Figure 5.**
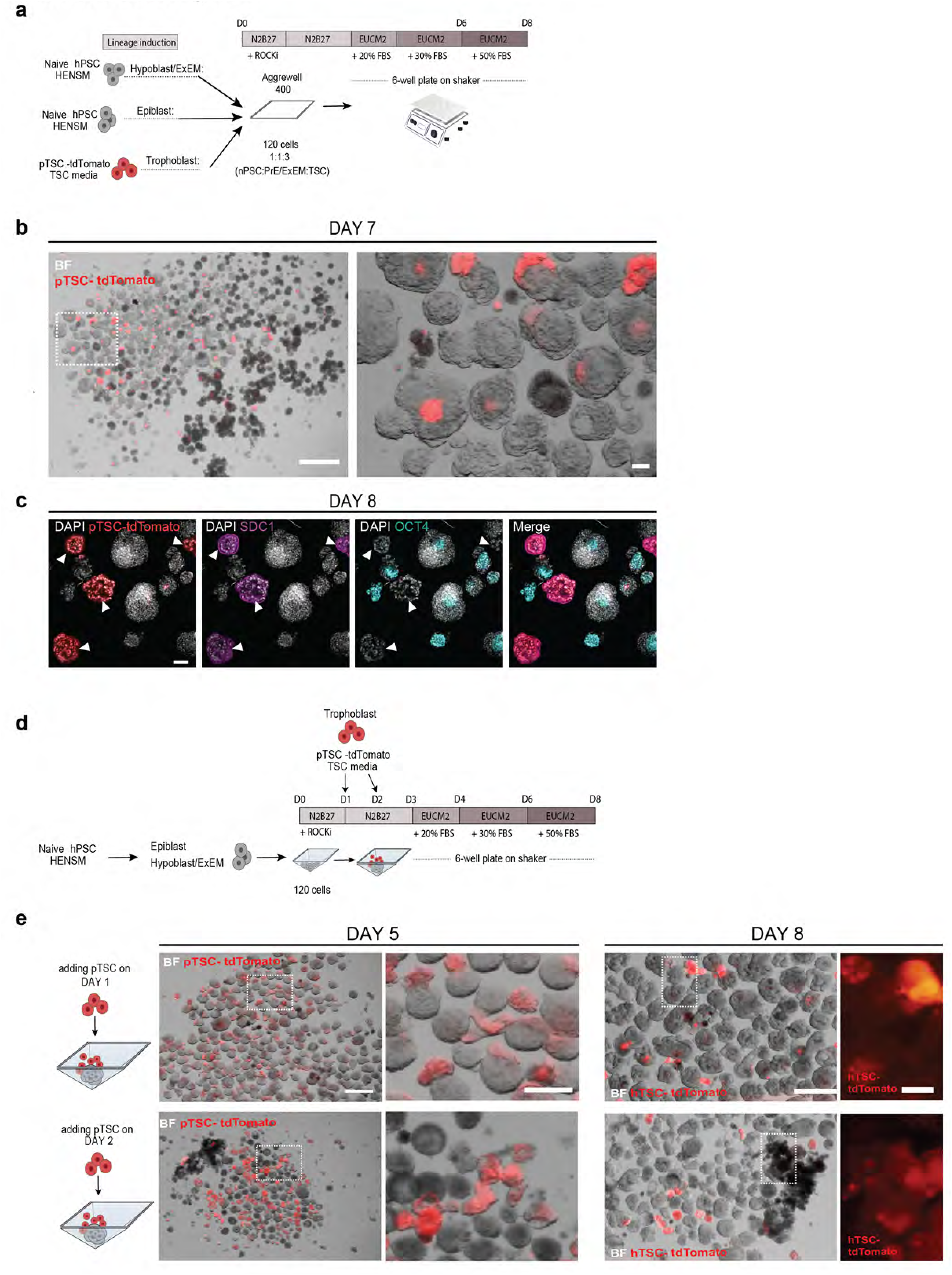
Testing SEM aggregations with human conventional trophoblast stem cell lines. **a**, scheme depicting aggregation protocol of epiblast (naïve hPSCs in HENSM media) and PrE/ExEM cells with tdTomato-expressing validated human TSC line derived from human primed ESCs (termed pTSC). **b**, brightfield images and live fluorescence of tdTomato (red) in aggregates with labeled TSCs; scale bar, 200 µm. Right, zoom into the several SEMs with tdTomato signal. Scale bar, 50 µm. The aggregates with TSCs do not form a uniformly surrounding TB layer, but rather remain as isolated clumps. **c**, immunofluorescence images of day 8 aggregates, showing OCT4 (cyan), SDC1 (magenta), and TSCs, labelled by tdTomato; nuclei (DAPI, white). Scale bar, 50 µm. White arrows highlight TSC isolated clumps. **d**, sequential aggregation of nPSCs (in HENSM) with tdTomato-expressing primed PSC derived TSC line on the first or second days of the aggregation protocol. **e**, brightfield images and live fluorescence of tdTomato (red) in day 5 and 8 aggregates with TSCs, added on the first (top) or second (bottom) days of aggregation; scale bar, 200 µm. Right, zoom into the several SEMs with tdTomato signal. Scale bar, 50 µm. The aggregates with TSCs do not form a uniformly surrounding TB layer, but rather remain as isolated clumps.

**Extended Data Figure 6.**
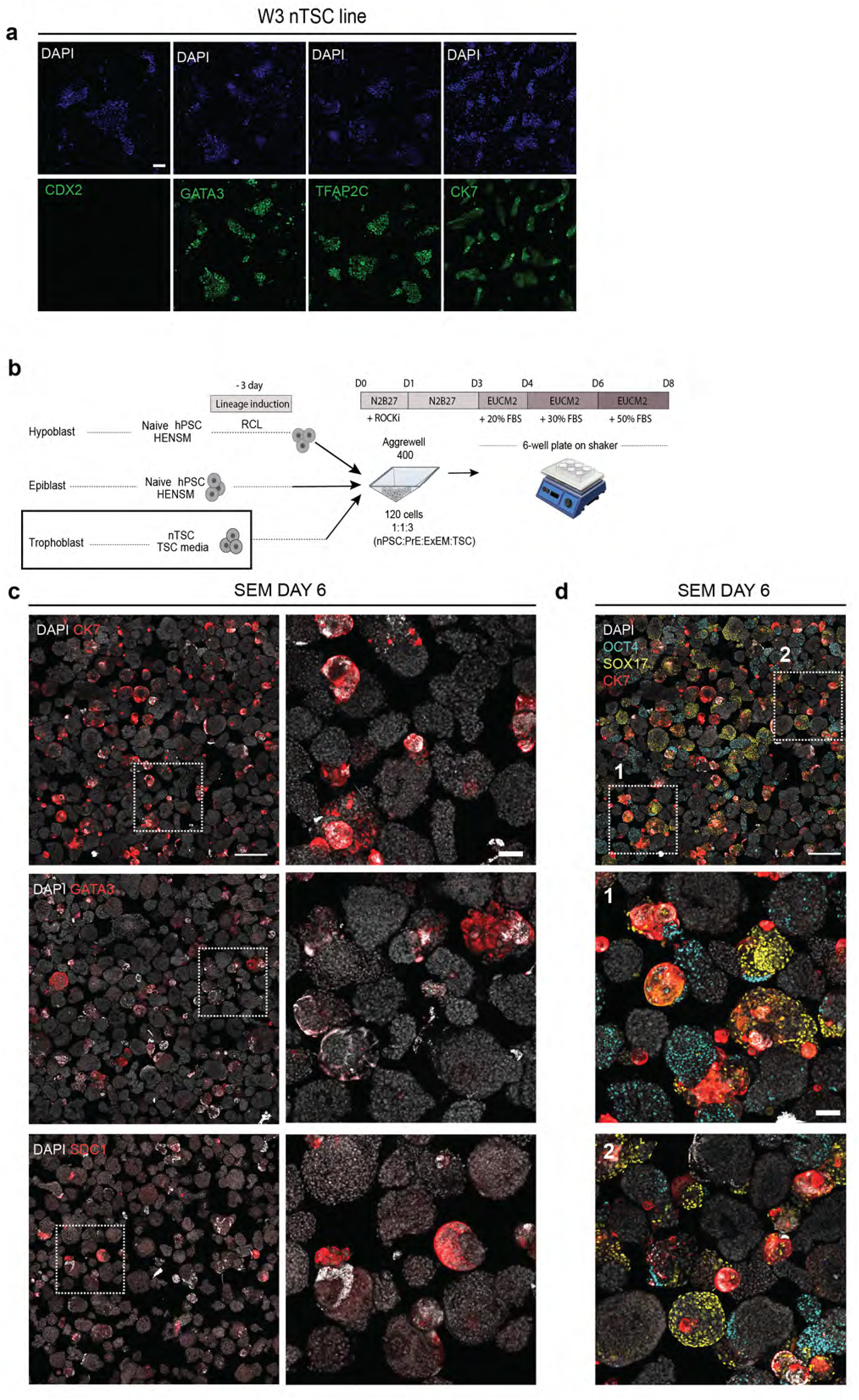
Testing SEM aggregations with naïve PSC derived human trophoblast stem cell lines. **a,** immunofluorescence images validating correct expression of TSC marker genes in colonies of a WIBR3 (W3) naïve hESC derived TSC line (termed nTSC). CK7, TFAP2C, GATA3, CDX2 (all in green); DAPI (blue). Scale bar, 100 µm. **b,** scheme for aggregation protocol of naïve pluripotent stem cells (nPSCs) in HENSM media, naïve-derived trophoblast stem cells (nTSCs), and nPSCs induced in RCL towards PrE/ExEM for 3 days. **c**, immunofluorescence images showing rare expression of CK7, GATA3, and SDC1 (red) trophoblast markers in the aggregates. nuclei (DAPI, white); Left column scale bar, 500 µm; Right, zoom into several aggregates with CK7 expression; scale bar, 100 µm. **d**, immunofluorescence image from **(upper left panel in c)**, showing Epi (OCT4, cyan) and PrE (SOX17, yellow) with CK7 (red); nuclei (DAPI, white). Zoom ins from the indicated regions are shown. Although some aggregates express lineage markers, they do not organize into embryoid-like structures and are not uniformly surrounded by the trophoblast compartment. Scale bar, 500 µm; bottom, 100 µm.

**Extended Data Figure 7.**
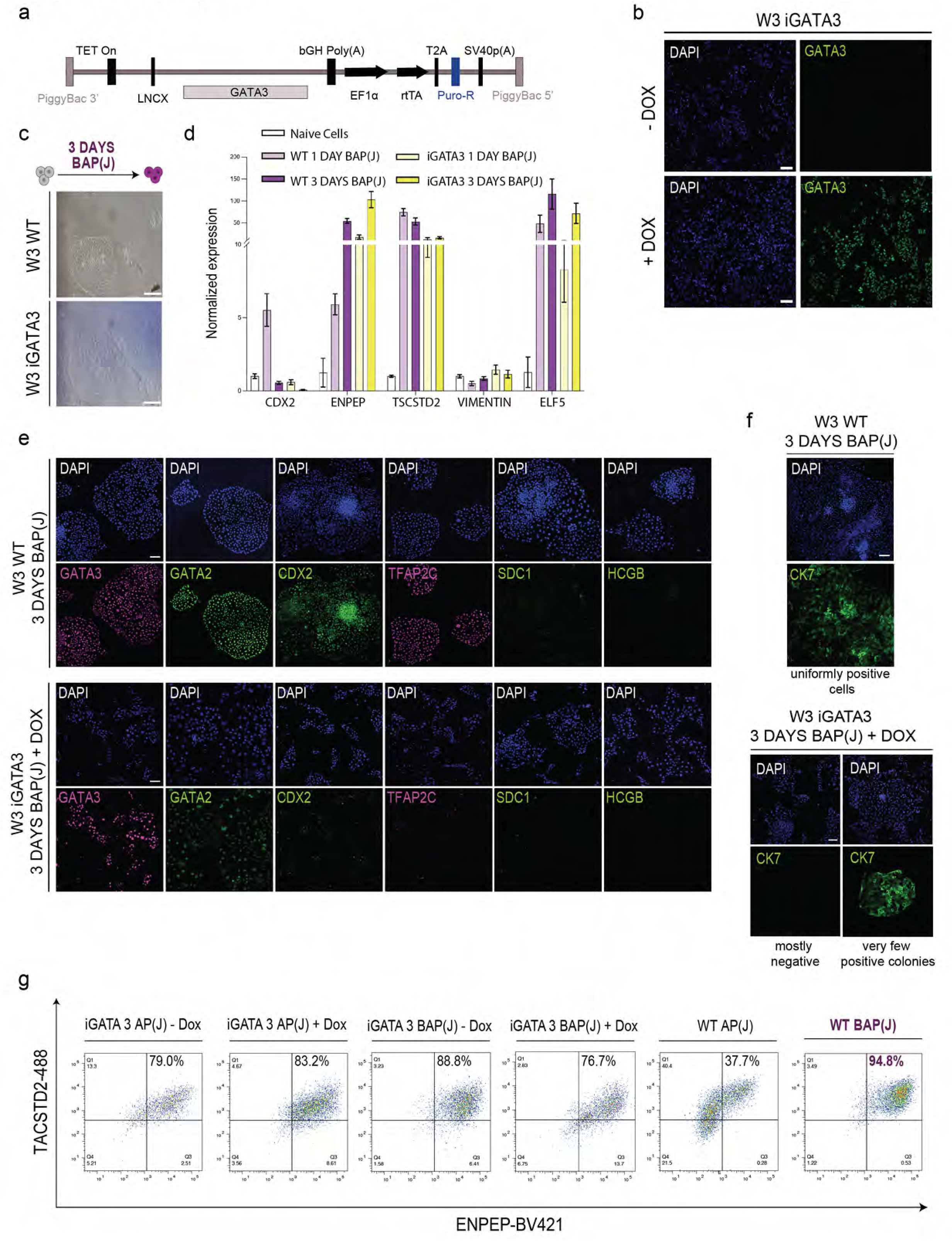
Evaluating trophoblast induction using transient overexpression of GATA3 in human naïve PSCs. **a,** scheme of the donor plasmid vector used for genomic integration of the DOX-inducible iGATA3 overexpression transgene. **b**, immunofluorescence images of iGATA3 cells, showing uniform GATA3 expression (green) only in response to DOX; nuclei (DAPI, blue). Scale bars, 100 µm. **c**, brightfield images of WT (top) and iGATA3 (bottom) cells after incubation in BAP(J) media for three days. Scale bar, 200 µm. **d**, qRT-PCR gene expression (normalized by GAPDH and ACTIN) of the trophoblast markers for WT naïve pluripotent stem cells (nPSCs) in BAP(J) media (purple) and iGATA3 nPSC cells induced by DOX in BAP(J) media (yellow), versus nPSCs maintained in HENSM media and used as a reference control (set as 1) (white). **e**, immunofluorescence images showing different patterns of expression of trophoblast marker genes in the wild type (WT) nPSCs incubated in BAP(J) media for three days (top) versus iGATA3 cells, induced by DOX in BAP(J) media (bottom). GATA3 (magenta), TFAP2C (magenta), GATA2, CDX2, SDC1, and HCGB (all in green), nuclei (DAPI, blue). Scale bar, 100 µm. **f**, immunofluorescence images showing uniform expression of CK7 in colonies of WT nPSCs incubated in BAP(J) media for three days (top), whereas colonies of iGATA3 cells induced by DOX in BAP(J) media express CK7 heterogeneously, with most of them being negative for CK7. Scale bar, 100 µm. **g**, from left to right, FACS plots of ENPEP versus TACSTD2 for trophoblast (Tb) priming using iGATA3 induction in different media (AP(J) with and without DOX, BAP(J) with and without DOX), and using WT nPSC priming to trophectoderm in AP(J) and BAP(J) regimens. Percentage of double positive population is indicated. This result shows that transient expression of the GATA3 transgene is not required for Tb differentiation and that it can be achieved with BAP(J) media using WT nPSCs as starting cells.

**Extended Data Figure 8.**
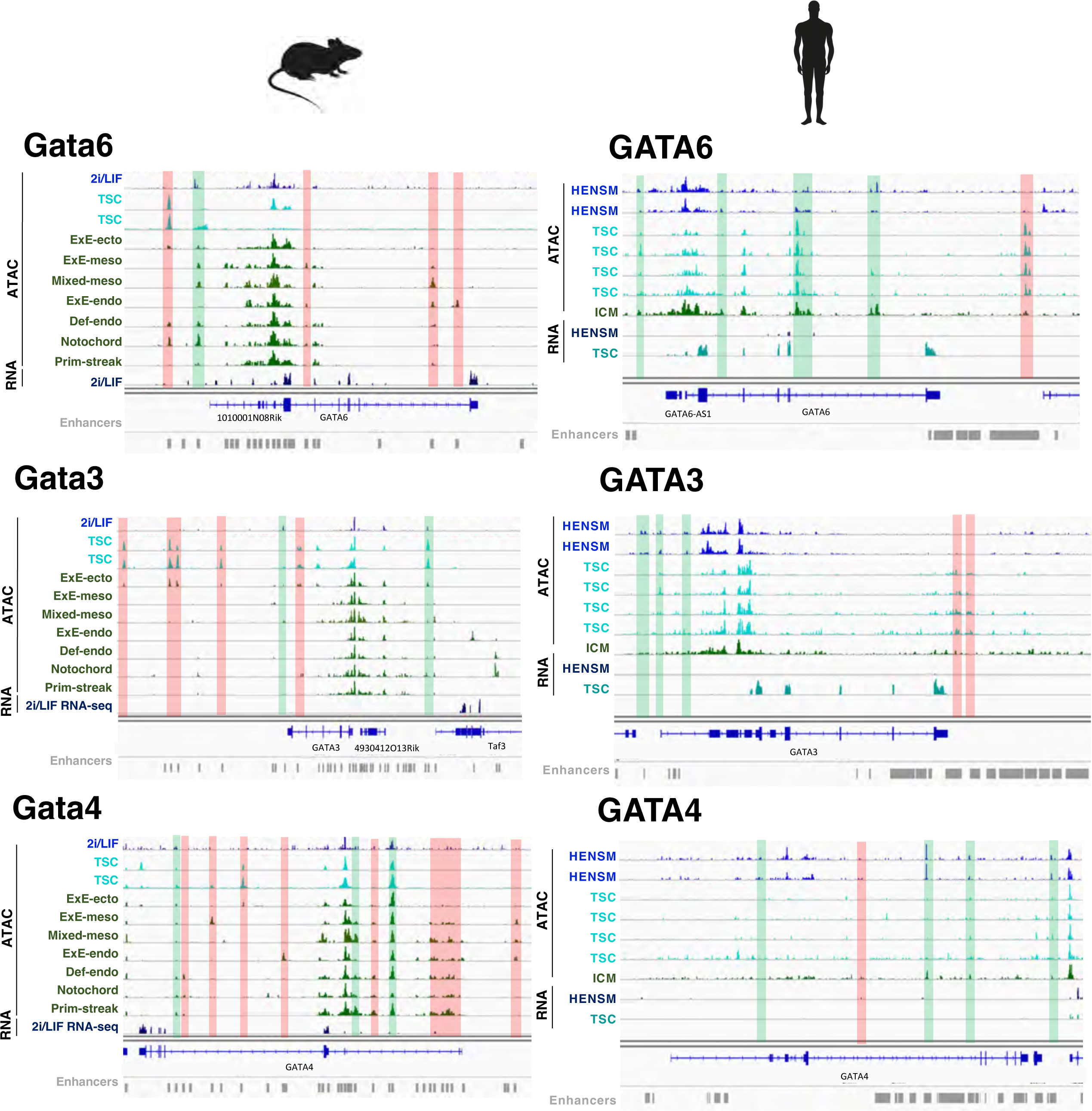
Enhancer accessibility of extra-embryonic lineage master regulators GATA3, GATA4 and GATA6 in human but not mouse naïve PSCs. ATAC-seq and RNA-seq of GATA3, GATA4 and GATA6 genes in human vs. mouse, as measured in naïve PSCs and in multiple differentiated cell types (as indicated). HENSM conditions were used for human naïve PSCs. Known enhancers are marked in grey bars at the bottom. Putative enhancers are marked in red or green: (1) green indicates potential enhancers that are already open in naïve stem cells, (2) red indicates enhancers that are closed in naïve pluripotent stem cells, but open in at least one differentiated cell state. Putative enhancers in the approximate regions of the genes were manually selected. Open regions that overlap with promoter or exon were excluded from the analysis. RNA-seq of the indicated genes in naïve PSCs are shown. The latter shows that while Gata6, Gat3 and Gata4 master regulators of extra-embryonic lineage differentiation^63, 127^ are not transcribed in mouse and human naïve PSCs, their enhancers are predominantly more accessible in human naïve PSCs, but not in mouse naïve PSCs.

**Extended Data Figure 9.**
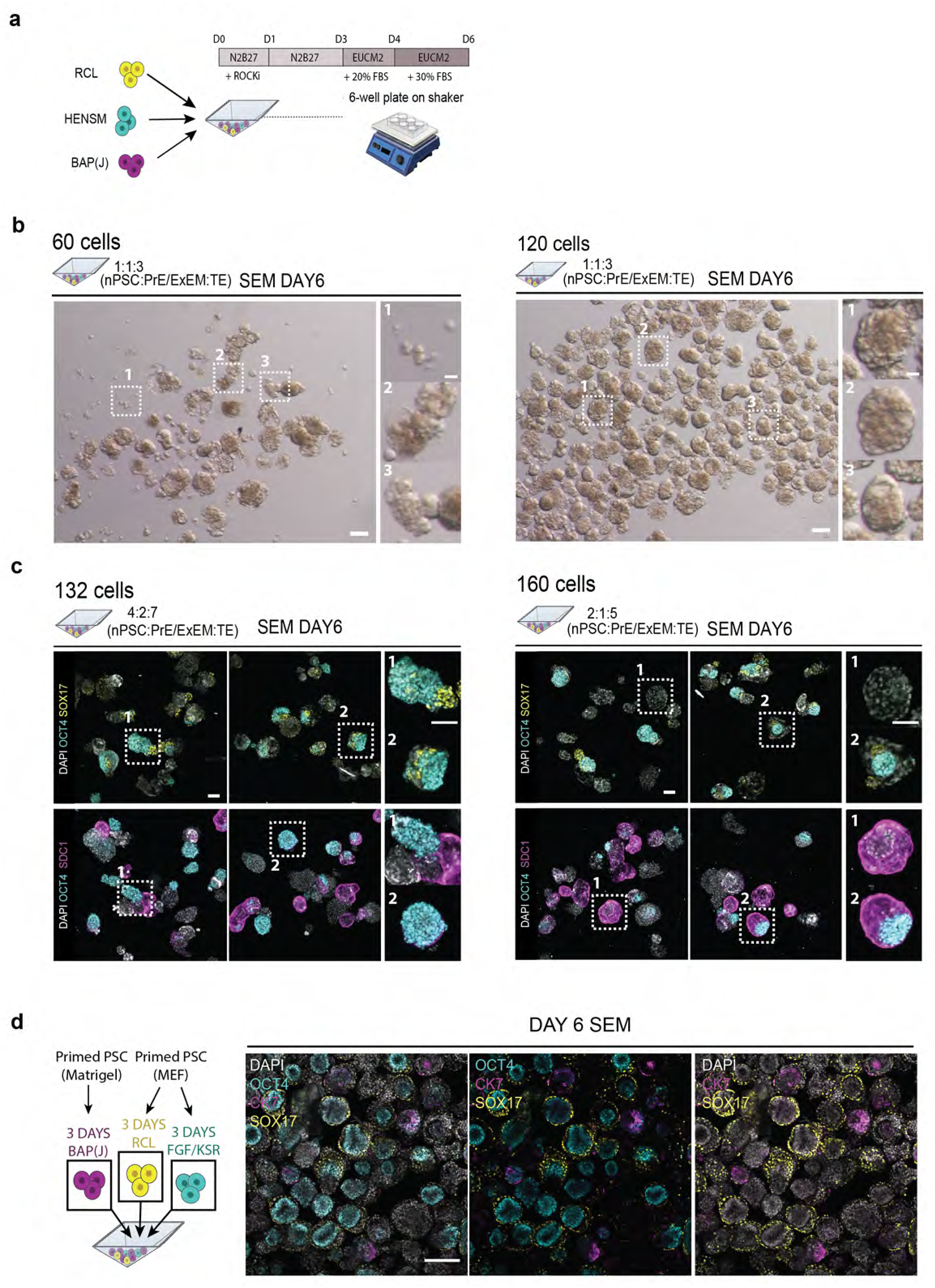
Optimization of the cell numbers co-aggregated for conducive human SEM formation exclusively from human naïve PSCs. **a**, scheme of the protocol: induced PrE/ExEM cells (yellow), WT naive pluripotent stem cells (nPSCs) maintained in HENSM medium (cyan), and WT nPSC induced towards trophoblast with BAP(J) medium (magenta) were aggregated on Aggrewell 400 platform at different ratios in N2B27 and grown as indicated in the scheme until day 6. **b**, left, brightfield images of day 6 SEMs, aggregated from total 60 cells at 1:1:3 (nPSC: PrE/ExEM: TE) cell ratio, showing mostly small and fragmented aggregates. Right, brightfield images of day 6 SEMs, aggregated from total 120 cells at 1:1:3 (nPSC: PrE/ExEM: TE) cell ratio, showing lower fragmentation tendency and more frequent formation of bigger aggregates (this condition was chosen for SEM generation). Scale bar, 1000 µm; zoom, 125 µm. **c**, immunofluorescence images of day 6 SEMs showing epiblast (OCT4, cyan), hypoblast (SOX17, yellow), and trophoblast (SDC1, magenta); nuclei (DAPI, white). Left, Day 6 SEMs aggregated from total 132 cells at 4:2:7 (nPSC: PrE/ExEM :TE) cell ratio, or, right, from total 160 cells at 2:1:5 (nPSC: PrE/ExEM :TE) cell ratio showing inadequate organization of SEMs, when compared to optimized conditions (120 cells at 1:1:3 nPSC: PrE/ExEM: TE) as shown in **Extended Data Figure 12c**. Scale bars, 100 µm. **d**, left, scheme of the experiment testing the capacity of cells, differentiated from human WIBR3 primed PSCs, to form equivalent SEMs to those obtained from isogenic naïve PSCs expanded in HENSM conditions. Right, immunofluorescence images of day 6 SEMs showing OCT4 (cyan), SOX17 (yellow), CK7 (magenta), and nuclei (DAPI, white). Scale bar, 200 µm. The resulting aggregates when starting with isogenic primed PSCs, did not present organization and maturation of the key embryonic and extra-embryonic compartments a seen when starting with naïve PSCs (please compare to Fig. 2).

**Extended Data Figure 10.**
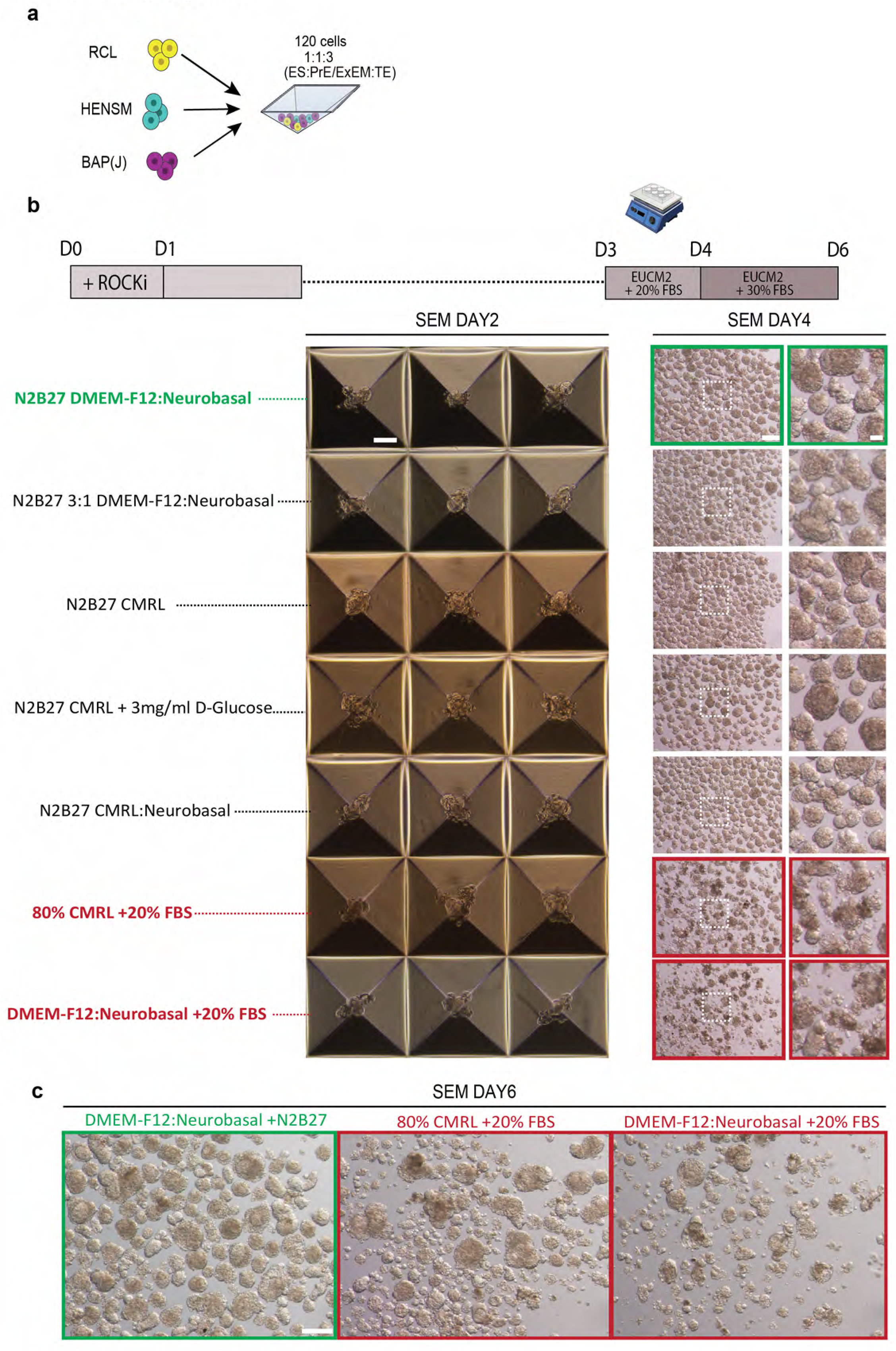
Optimization of the aggregation conditions for human SEM generation from naïve PSCs. **a**, scheme of the protocol: PrE/ExEM induced cells in RCL medium (yellow), WT naive pluripotent stem cells (nPSCs) maintained in HENSM medium (cyan), and iGATA3 trophectoderm cells induced with DOX in BAP(J) medium (magenta) were aggregated in different media in Aggrewell 400 plate at 1:1:3 (nPSC: PrE/ExEM: TE) ratio and cultured until day 6. **b**, from bottom to top, representative brightfield images of day 2 and 4 SEM, aggregated in DMEM-F12:Neurobasal (1:1) supplemented with 20% FBS or N2B27, CMRL 1066 base medium supplemented with 20% FBS or N2B27, CMRL supplemented with N2B27 and extra-added 3 mg/ml D-Glucose, and DMEM F12: Neurobasal (3:1). Although addition of FBS is beneficial for SEM growth, FBS impairs aggregation by day 4, and the best aggregation efficiency was consistently observed in DMEM-F12:neurobasal supplemented with N2B27 without FBS. **c**, representative brightfield images showing of SEM morphology at day 6 after aggregation in DMEM-F12:Neurobasal and CMRL supplemented with 20% FBS. Addition of FBS impaired formation of the aggregates while N2B27 conditions allowed better human PSC derived aggregate growth. Scale bars, 200 µm.

**Extended Data Figure 11.**
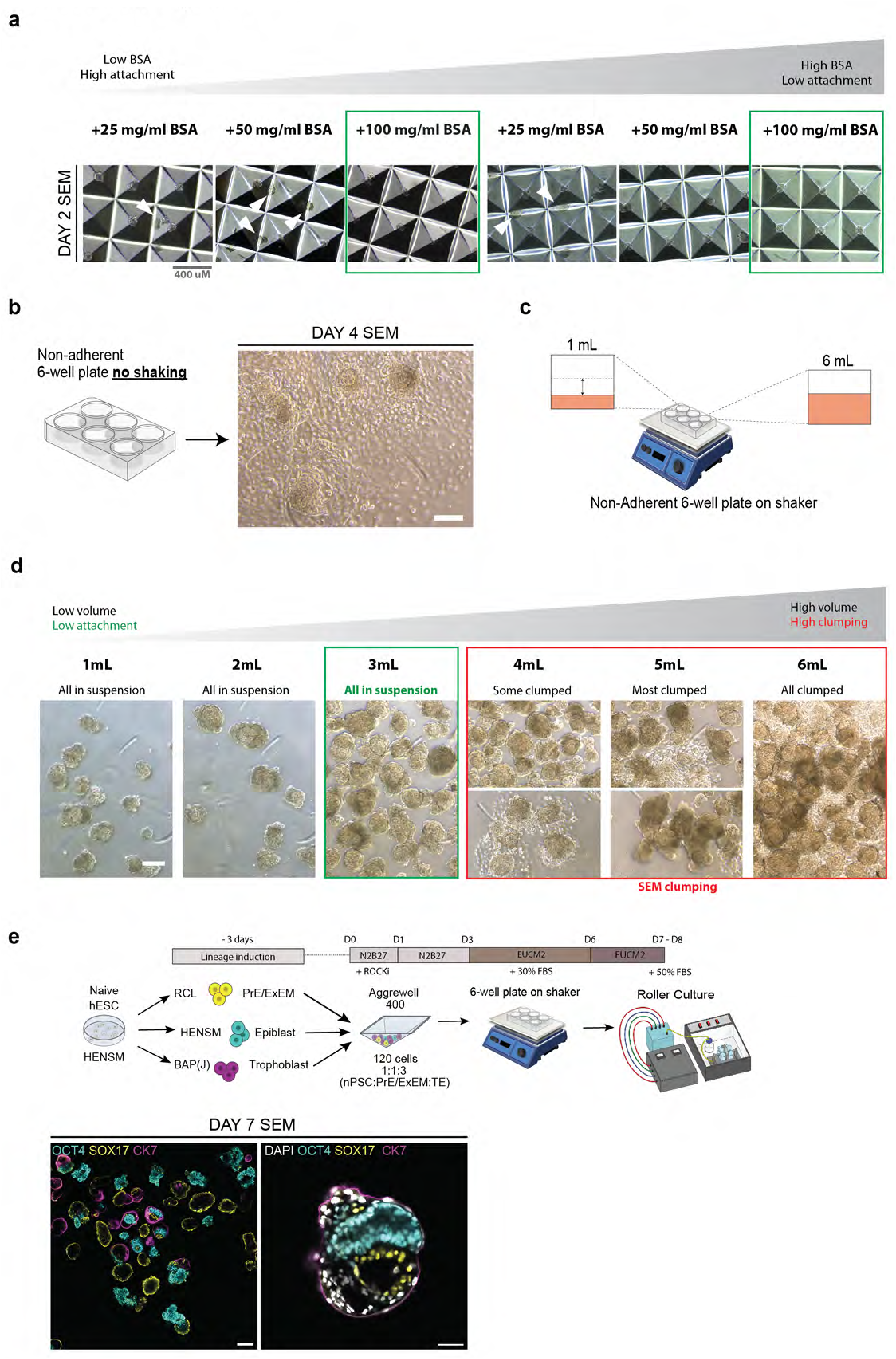
Optimization of human SEM culture conditions. **a**, brightfield images of day 2 SEM aggregates inside the Aggrewell 400 plate; addition of BSA to the aggregation medium prevents attachment of the aggregates to the plate edges (white arrows). Scale bar, 400 µm. Optimal BSA concentration chosen for further experimentation highlighted by a green box. **b**, growth of SEMs in 6-well non-adherent plates without shaking leads to their attachment to the plate and disruption of morphology. Scale bar, 200 µm. **c**, scheme of the experiment in which volume of the culture media on the non-adherent 6-well dish, using orbital shaking (day 4-8), was optimized. **d**, brightfield images of multiple SEMs showing that the rate of their clumping with each other is dependent on the media volume (1 – 6 ml). The optimal suspension condition is outlined in green (3ml). Scale bar, 200 µm. **e**, top, the scheme of roller culture test, where aggregates were generated as described (see Methods), cultured until day 6 on a shaker, followed by roller culture in EUCM2 50% FBS 20%O_2_ 5%CO2. Bottom, immunofluorescence images of day 7 SEMs in this regimen, showing Epi (OCT4, cyan), hypoblast (SOX17, yellow), and trophoblast (CK7, magenta); nuclei (DAPI, white), brightfield (BF); scale bar, 200 µm. Right, example of the SEM in which epiblast, hypoblast, and trophoblast are adequately compartmentalized; scale bar, 50 µm.

**Extended Data Figure 12.**
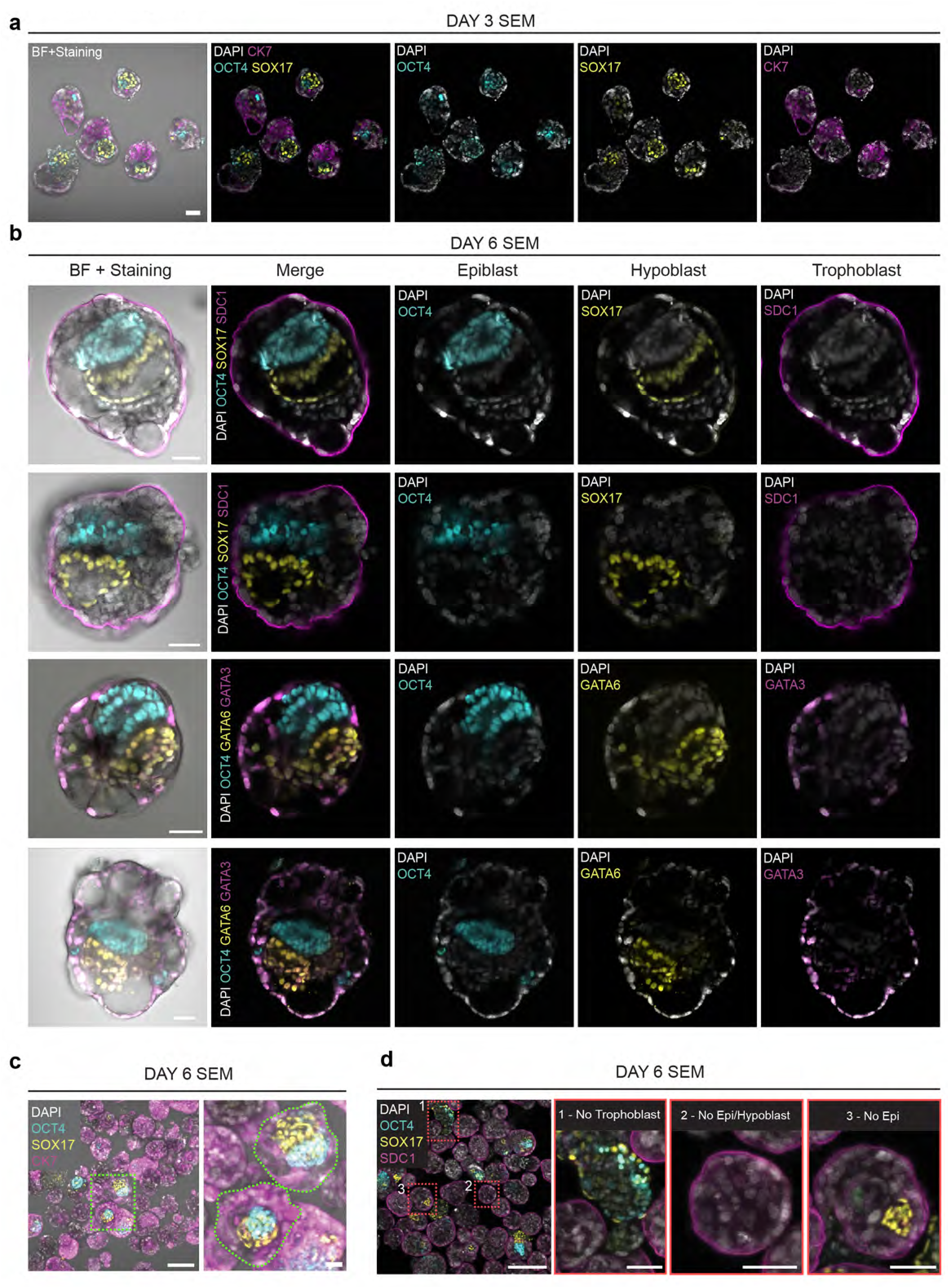
Characterization of epiblast, hypoblast, and trophoblast lineages in SEMs. **a**, from left to right, representative merged brightfield (BF) and immunofluorescence images of day 3 SEMs showing three lineages, epiblast (OCT4, cyan), hypoblast (SOX17, yellow), and trophoblast (SCD1, magenta) merged with a nuclei channel (DAPI, white). **b**, representative merged brightfield and immunofluorescence images of multiple day 6 SEMs, showing epiblast (OCT4, cyan), hypoblast (SOX17, yellow), and trophoblast (CK7, magenta) merged with a nuclei channel (DAPI, white). **c**, immunofluorescence images of day 6 SEMs with segregated Epi (OCT4, cyan) and hypoblast (SOX17, yellow), and surrounded by trophoblast (CK7, magenta, outlined). Left panel shows a wide field image to identify adequate structures (dotted square). Right panel zooms into the structure of interest **d**, immunofluorescence images of common examples of mis-developed day 6 SEMs with no trophoblast (1), or no epiblast and hypoblast (2), or no epiblast (3). Epiblast (OCT4, cyan), hypoblast (SOX17, yellow), and trophoblast (SDC1, magenta); nuclei (DAPI, white). Scale bars, 50 µm, 200 µm (c right, d right).

**Extended Data Figure 13.**
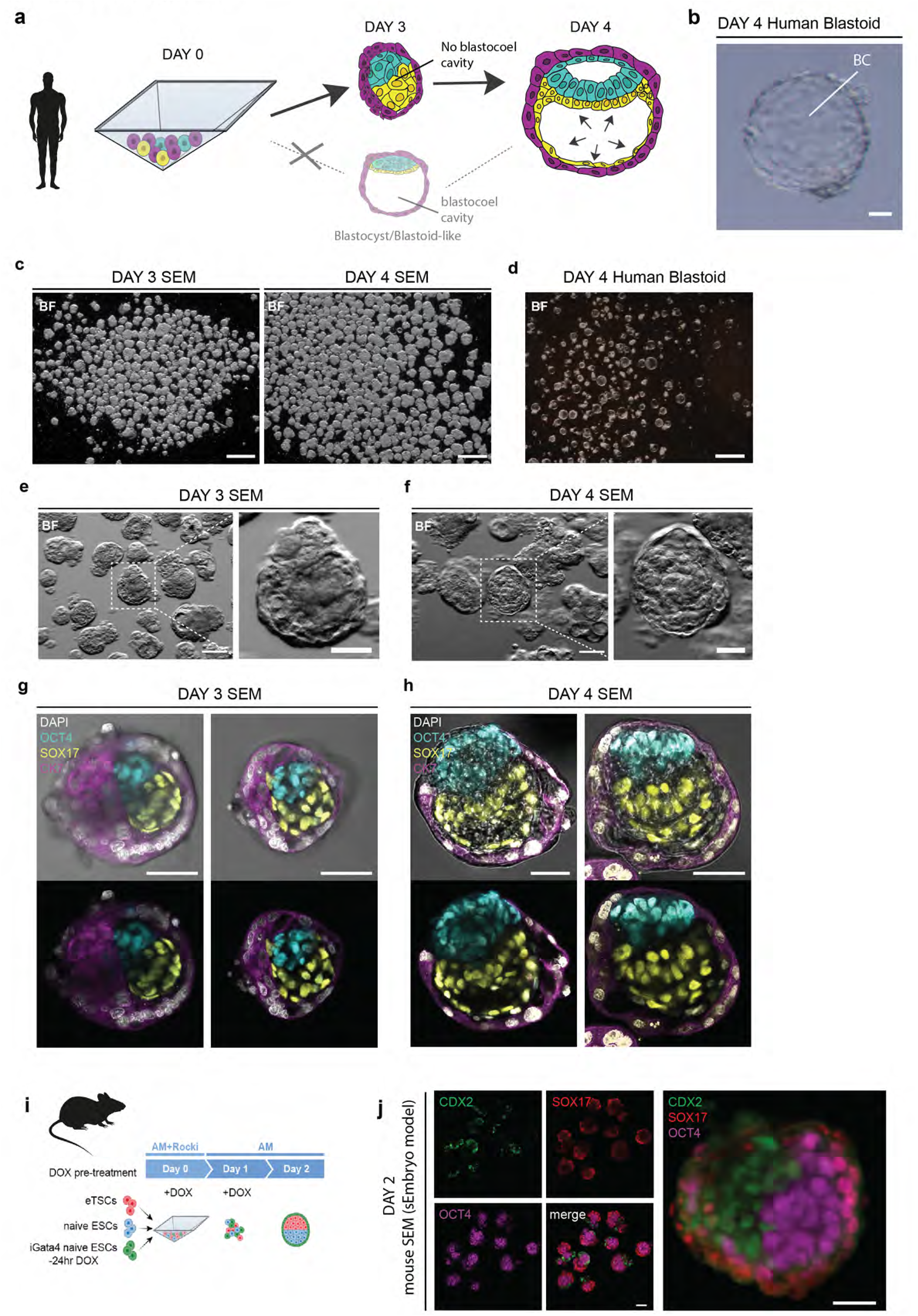
Human and mouse SEMs bypass blastocyst-like stage. **a,** scheme showing the distinction between human SEM (top) and human blastoid (bottom) aggregation protocols: as opposed to blastoids, SEMs do not form a blastocoel cavity. Epiblast (cyan), hypoblast (yellow), trophoblast (magenta). **b**, representative brightfield image of a blastoid with blastocoel cavity (BC). Scale bar, 100 µm. **c**, representative examples of brightfield images of day 3 and day 4 SEMs showing no blastocoel cavity. Scale bar, 200 µm. **d**, brightfield images of day 4 human blastoids with a BC cavity. Scale bar, 200 µm. **e-f**, enlarged brightfield (BF) images of the SEMs showing no blastocoel cavity at day 3 or day 4 human SEMs **(f).** Scale bar, 100 µm; zoom in, 50 µm. **g-h**, brightfield and immunofluorescence images of day 3 **(g)** and day 4 **(h)** human SEMs showing developmental progression of epiblast (OCT4, cyan), hypoblast (SOX17, yellow), and trophoblast (CK7, magenta); nuclei (DAPI, white). Scale bars, 50 µm **i**, scheme showing mouse SEM (also known as sEmbryo) aggregation with murine stem cells as described in ^27^. **j**, immunofluorescence images showing epiblast (OCT4, magenta), hypoblast (SOX17, red), and trophectoderm (CDX2, green); Like human SEMs, mouse SEMs (sEmbryo models) do not form a BC. Notably mouse SEMs are predominantly surrounded by the PrE, rather than TE by Day 2 of the protocol. Scale bars, 50 µm; zoom in, 20 µm.

**Extended Data Figure 14.**
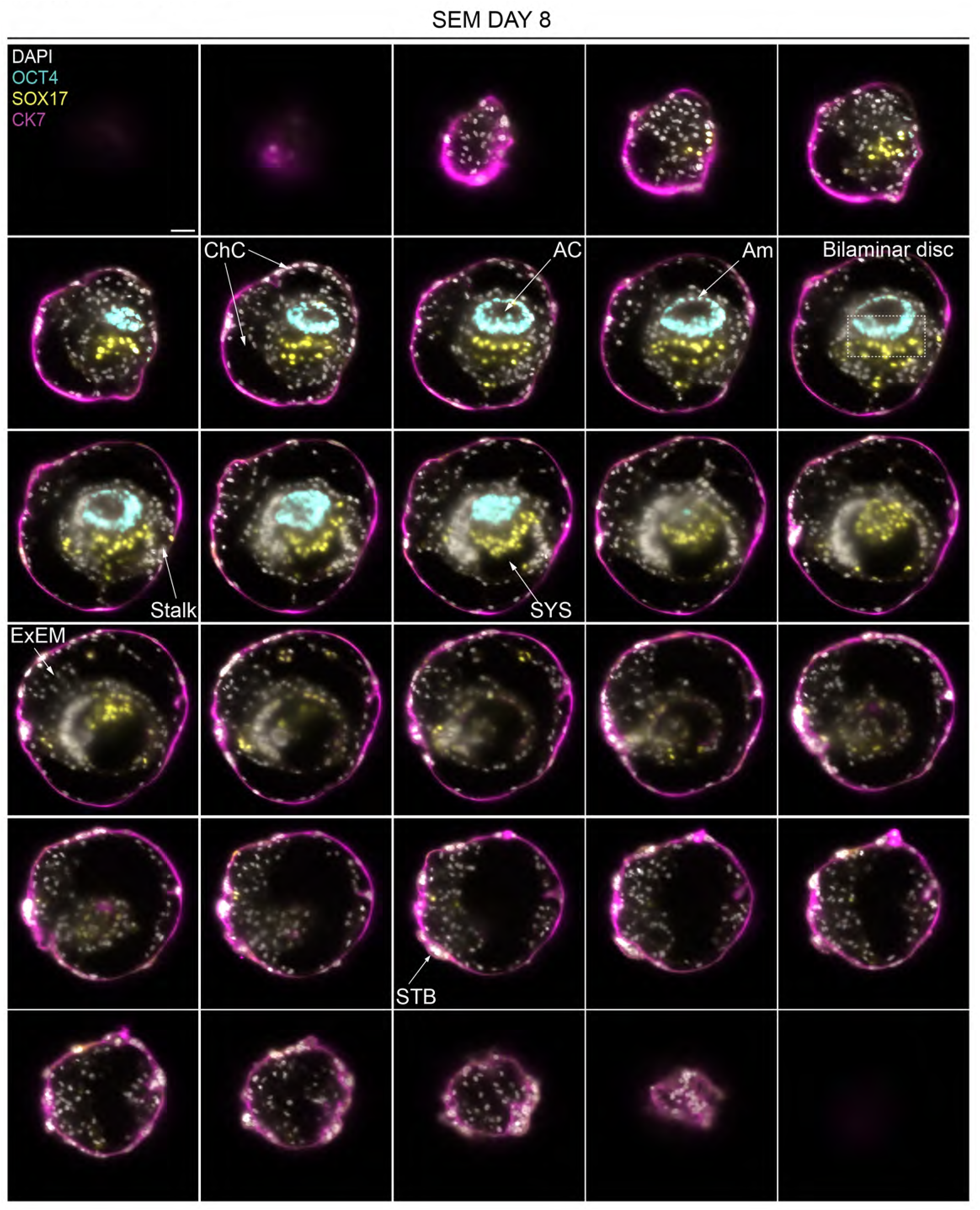
3D structure of a human SEM at day 8. **a**, individual Z-planes of the 3D immunofluorescence image of day 8 human SEM. Epiblast (OCT4, cyan), hypoblast (SOX17, yellow), and trophoblast (CK7, magenta); nuclei (DAPI, white). AC, amniotic cavity; Am, amnion; ExEM, extraembryonic mesoderm; SYS, secondary yolk sac; ChC, chorionic cavity; STB, syncytiotrophoblast. Z-step, 20 µm; scale bar, 50 µm. See also **Video S1**.

**Extended Data Figure 15.**
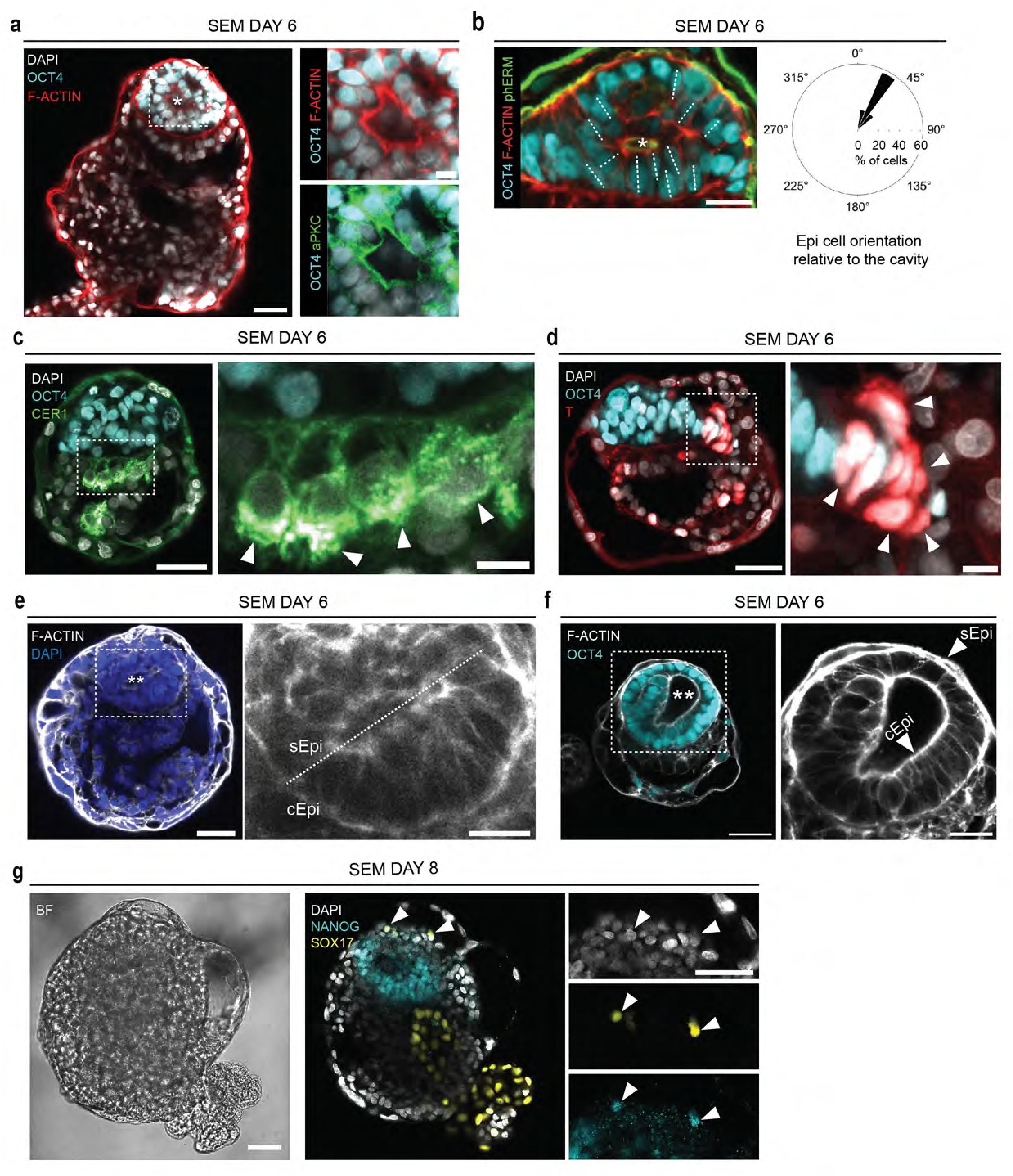
Characterization of the epiblast in human SEM. **a**, immunofluorescence image of day 6 SEM showing aPKC (green), OCT4 (cyan), F-ACTIN (red), nuclei (DAPI, white). Scale bars, 50 µm, 12.5 µm (zoom in, right). **b**, left, immunofluorescence image of day 6 SEM showing OCT4 (cyan), F-ACTIN (red), and phERM (green); alignment of epi cells in a single 2D plane is marked with dashed lines; asterisk, pro-amniotic cavity. Scale bar, 25 µm. **b**, right, the angle between the epiblast cell axis and the pro-amniotic cavity was quantified; the plot shows the radial histogram of the angle values. The latter shows predominant alignment of epi cells towards the center of the emerging cavity. **c**, immunofluorescence image of day 6 SEM showing CER1 (green) localization inside the intracellular vesicles; OCT4 (cyan), nuclei (DAPI, white). Right, zoom into hypoblast, arrows mark apical side of the visceral endoderm cells with CER1 vesicles. Scale bar, 50 µm, 12.5 µm (4x zoom in, right). **d**, immunofluorescence image of day 6 SEM showing T/Bra expression (red); OCT4 (cyan), nuclei (DAPI, white). Right, zoom into the posterior epiblast, arrows mark individual T/Bra-positive cells. Scale bar, 50 µm, 12.5 µm (4x zoom in, right). **e**, immunofluorescence image of day 6 SEM showing F-ACTIN (white) and nuclei (DAPI, blue). Right, zoom into epiblast with squamous cells (sEpi) in the top and cylindrical/columnar cells (cEpi) in the bottom parts of the epiblast. Scale bar, 50 µm; zoom in, 25 µm. **f**, immunofluorescence image of day 6 SEM showing F-ACTIN (white) and OCT4 (cyan). Right, zoom into epiblast with squamous cells in the top and cylindrical cells in the bottom parts of the epiblast. Scale bar, 50 µm; zoom in, 25 µm. **g**, brightfield and immunofluorescence images of day 8 SEM showing NANOG (cyan), SOX17 (yellow), and nuclei (DAPI, white). Right, zoom onto PGCs in day 8 human SEMs, co-expressing NANOG and SOX17 (cells marked with arrows). Scale bars, 50 µm.

**Extended Data Figure 16.**
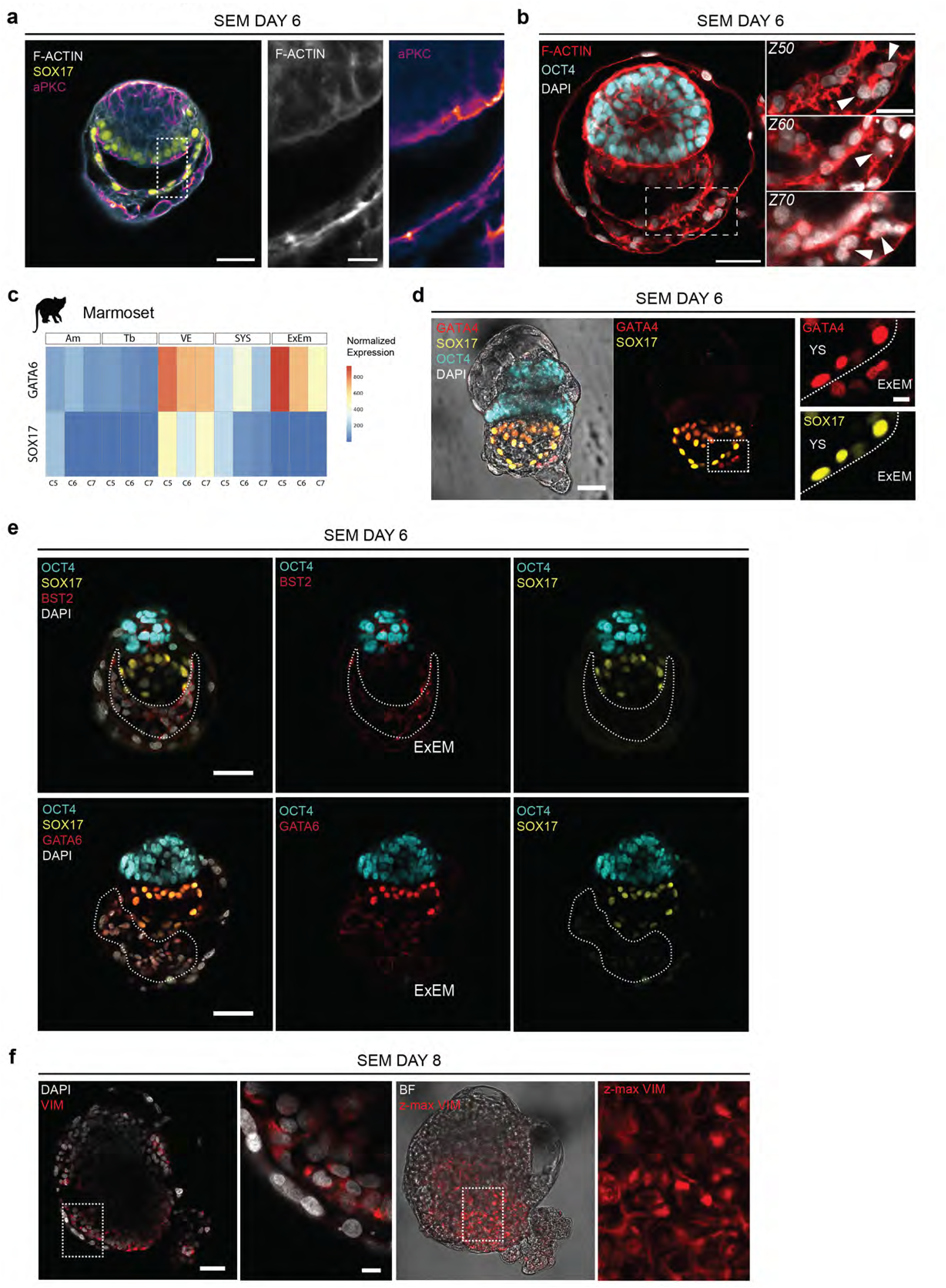
Characterization of hypoblast and extraembryonic mesoderm in human SEMs. **a**, immunofluorescence image of day 6 SEM showing apical polarity of the visceral and parietal hypoblast (SOX17, yellow); aPKC (heat gradient), F-ACTIN (white). Scale bar, 50 µm; zoom in, 10 µm. See also **Video S3**. **b**, immunofluorescence image of day 6 SEM showing mesenchymal-like cells underneath the yolk sac; OCT4 (cyan), F-ACTIN (red), and nuclei (DAPI, white). Right, zoom into the region underneath the yolk sac, shown in different Z-planes (number 50, 60, and 70). Arrows point at the cells between yolk sac and the trophoblast compartments. Scale bar, 50 µm; zoom in, 25 µm. See also **Video S2. c**, heatmap of GATA6 and SOX17 gene expression across extraembryonic tissues in marmoset, corresponding to the Carnegie stages (CS) 5-7), extracted from previously published study related gene expression dataset ^90^. Am, amnion; Tb, trophoblast; VE, visceral endoderm; SYS, secondary yolk sac; ExEM, extraembryonic mesoderm. **d**, immunofluorescence image of day 6 SEM showing OCT4 (cyan), GATA4 (red), SOX17 (yellow), and nuclei (DAPI). Right, zoom on ExEM cells expressing GATA4, but not SOX17. YS, yolk sac. Scale bar, 50 µm; zoom in, 10 µm. **e,** immunofluorescence images of day 6 SEMs showing chorionic cavity surrounded by ExEM (outlined), negative for SOX17 (yellow), but expressing BST2 (top, red) and GATA6 (bottom, red); OCT4 (cyan), nuclei (DAPI, white). Scale bar, 50 µm. **f**, left, immunofluorescence image of Z slice from day 8 SEM and the zoom into the ExEM region showing VIM (red) expression and nuclei (DAPI, white). Right, merged brightfield and maximum intensity projection showing VIM (red) expression. Scale bar, 50 µm; zoom in, 10 µm.

**Extended Data Figure 17.**
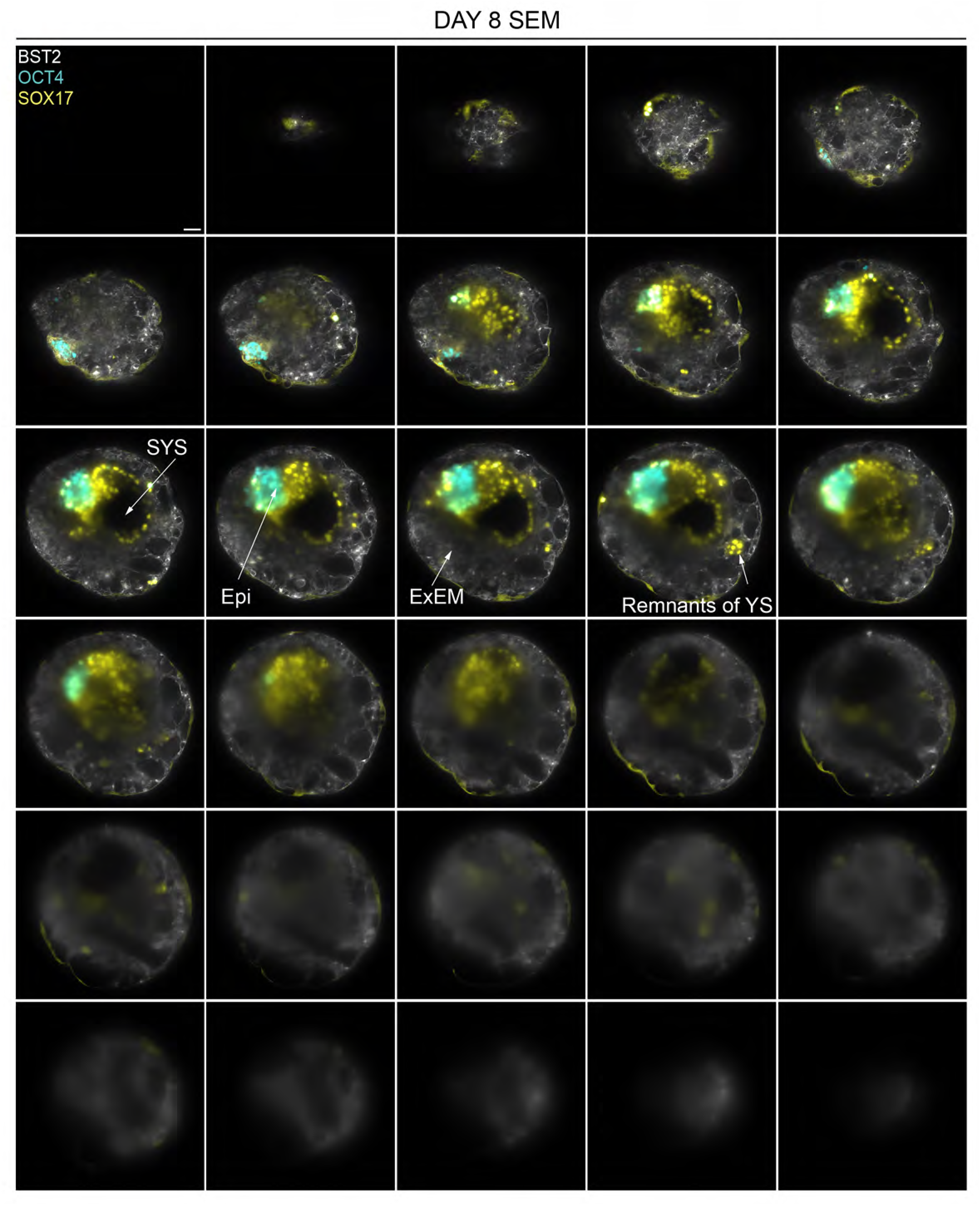
3D structure highlighting the ExEM in human SEM at day 8. **a**, individual Z-planes of the 3D immunofluorescence image of day 8 human SEM showing OCT4 (cyan), SOX17 (yellow), and BST2 (white). Epi, epiblast; YS, yolk sac; ExEM, extraembryonic mesoderm; SYS, secondary yolk sac; Z-step, 20 µm; scale bar, 50 µm.

**Extended Data Figure 18.**
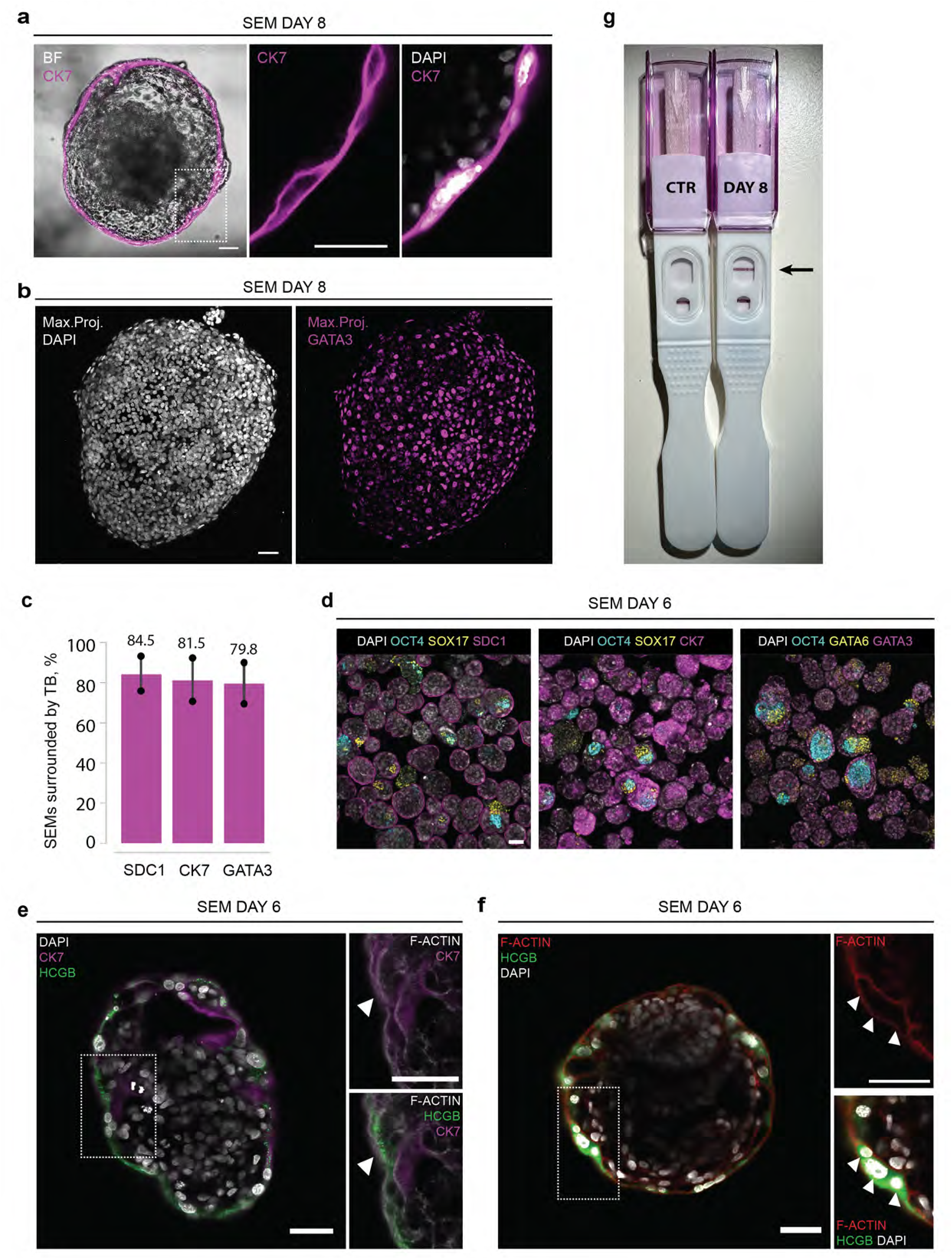
Characterization of the trophoblast compartment in human SEM. **a**, brightfield image overlayed with the immunofluorescence of day 8 SEM showing the outer trophoblast (CK7, magenta) surrounding the entire SEM. Right, zoom into the multinucleated trophoblast layer. Scale bars, 50 µm. **b**, the maximum intensity projection (Maxi. Proj.) of the immunofluorescence image of day 8 SEM showing trophoblast (GATA3, magenta) and nuclei, (DAPI, white). Scale bar, 50 µm. **c**, quantification of the percentage SEMs surrounded by trophoblast lineage (generated from WT naïve PSCs following 3 day BAP(J) protocol), as judged by the expression of SDC1, CK7, and GATA3 of day 6, across two independent experimental replicates. The total number of SEMs used for quantification are: SD1 (n1=525 and n2=769); CK7 (n1=299 and n2=466) and GATA3 (n1=150 and n2=337). **d**, representative immunofluorescence images of multiple day 6 SEMs showing epiblast (OCT4, cyan), hypoblast (SOX17 and GATA6, yellow), and trophoblast (SDC1 and GATA3, magenta) surrounding the SEMs; nuclei (DAPI, white), that were used for calculation presented in (**c**). Scale bar, 200 µm. **e**, immunofluorescence image of day 6 SEM showing HCGB (green) and CK7 (magenta) staining. HCGB positive syncytiotrophoblast cells (right, zoom) are highlighted. nuclei (DAPI, white), F-actin (white, right zoom). Arrows point at the outer syncytiotrophoblast cell surface. Scale bars, 50 µm. **f**, immunofluorescence image of a human SEM showing multinucleated HCGB-positive syncytiotrophoblast; HCGB (green), F-ACTIN (red), nuclei (DAPI, white). Arrows point at multiple nuclei inside the same single cell as validated following F-actin staining. Scale bars, 50 µm. **g**, pregnancy test run on spent medium of the day 8 SEMs (Day 8-right) compared to unspent medium as a negative control (CTR-left) which shows the secretion of HCGB from the syncytiotrophoblast of day7-8 human SEM to the culture medium.

**Extended Data Figure 19.**
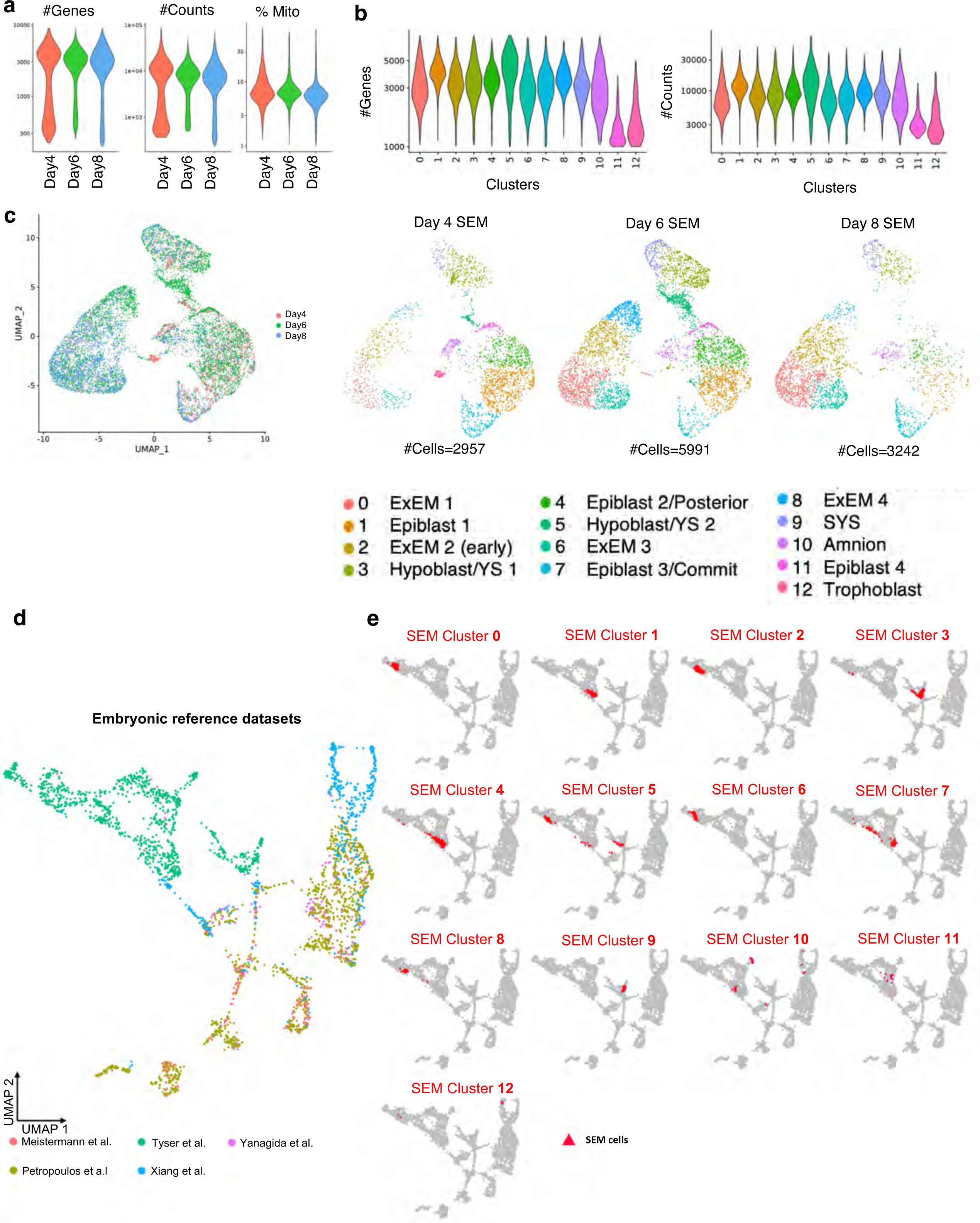
Human SEM UMAP quality assessment of scRNA-seq experiment. **a,** Violin plots indicating the number of genes, the unique molecular identifiers (UMIs), and the percentage of mitochondrial reads obtained per sample prior to filtering. **b,** Violin plots indicating the number of genes and the unique molecular identifiers (UMIs) obtained per cluster, after filtering cells with less than 1000 identified genes. **c**, UMAP plot displaying individual cells of the different SEM samples as indicated. Left: Colors indicate different samples. Right: Colors indicate the 13 identified clusters as indicated in Fig. 6a. The number of single cells in each sample is indicated. **d**, UMAP projection of the assembled human embryonic reference dataset with color indicating data source (related to Fig. 6e). **e**, Highlights the projected SEMs cells on the embryonic reference UMAP space stratified by the corresponding raw Seurat clusters. The grey data points represent embryonic reference cells or unselected neighborhood nodes, while each red triangle corresponds to the projection of representative neighborhood nodes from each SEM raw cluster onto the UMAP space (related to Fig. 6e).

**Extended Data Figure 20.**
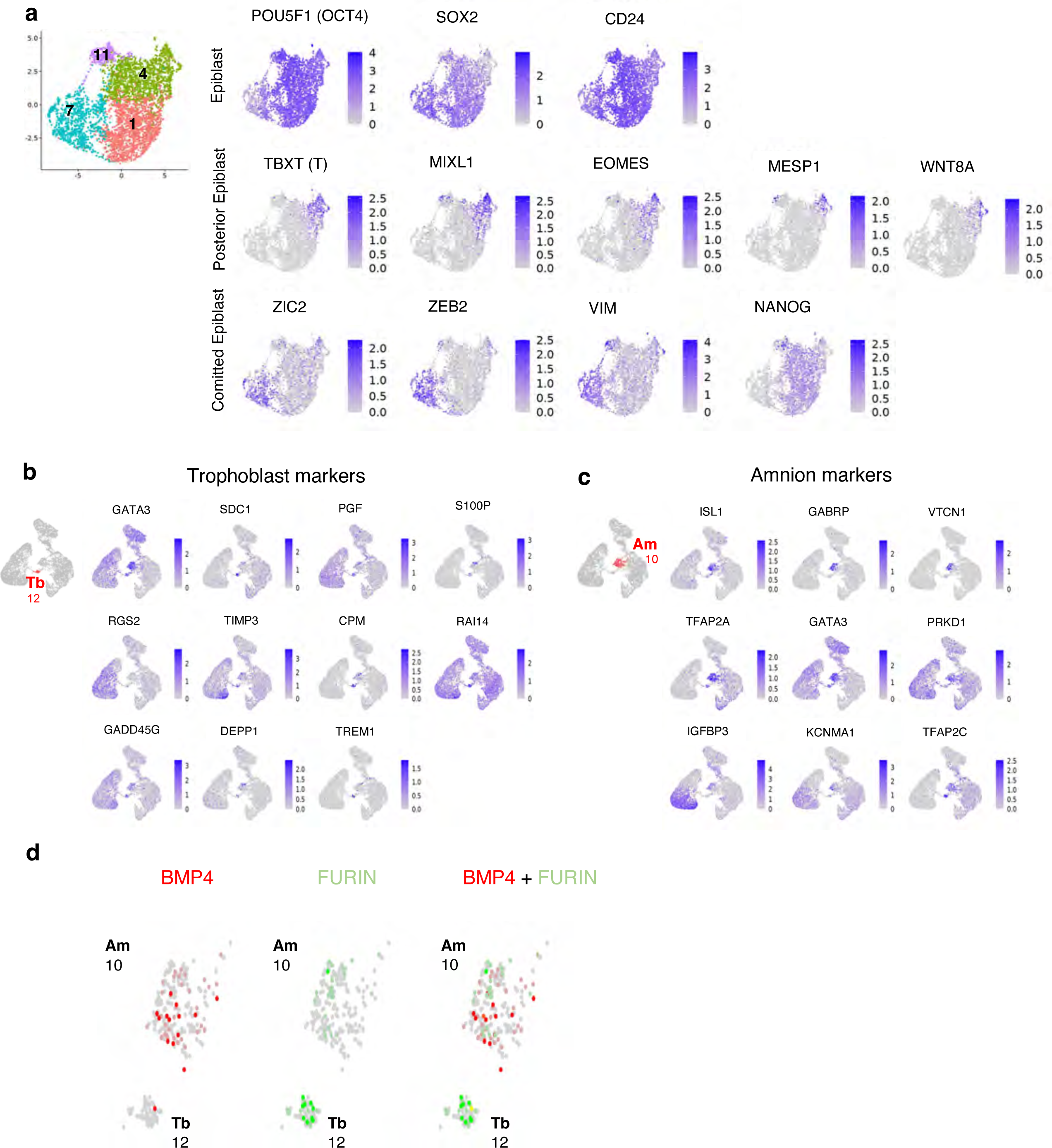
Identification and validation of specific cell sub-types in human SEMs. **a,** UMAP of the four annotated epiblast clusters (1,4,7 & 11) alongside normalized expression of key marker genes. From the four Epi clusters we subclassified 2 of them. The first we termed Posterior epiblast cluster (#4) was marked by upregulation of TBXT (Brachyury), MIXL1, EOMES, MESP1 and WNT8a which are markers of EMT. The second we termed as “committed epiblast” cluster (#7) and was marked by ZIC2, ZEB2, VIM lineage commitment marker expression and absence of NANOG while maintaining OCT4 and SOX2 expression. **b**, Normalized expression of key trophoblast marker genes projected on SEM UMAP. Trophoblast (Tb) cluster (number 12) is highlighted in red. **c**, Normalized expression of key amnion marker genes projected on human SEM related UMAP. Amnion cluster (number 10) is highlighted in red. **d**, Expression of BMP4 and/or FURIN in Amnion (Am) and Trophoblast (Tb) clusters.

**Extended Data Figure 21.**
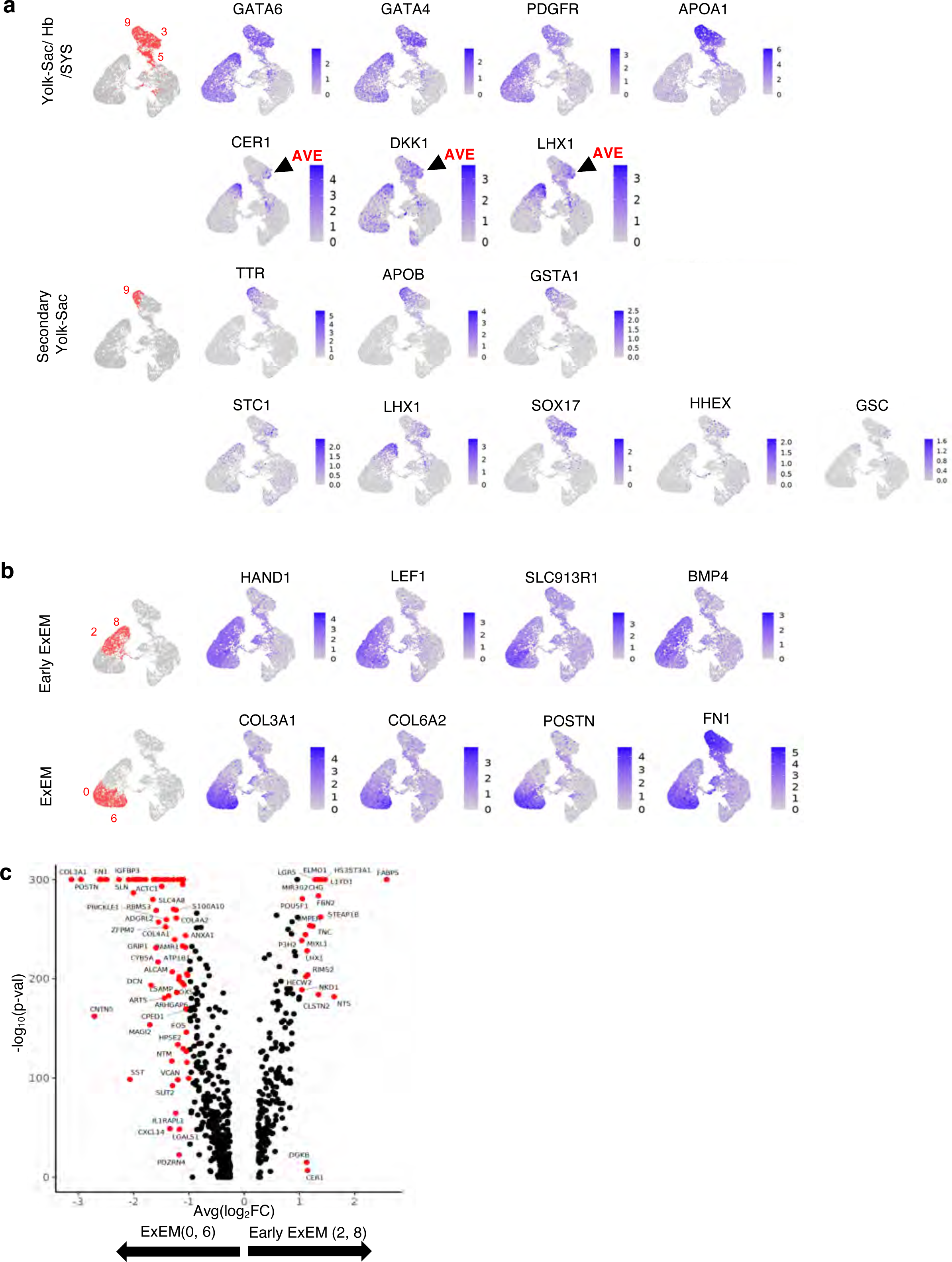
Identification of Primary and Secondary Yolk-sac, and Extra-Embryonic Mesoderm lineages. **a,** Normalized expression of key Yolk-Sac (YS) marker genes projected on SEM UMAP. GATA4, GATA6, PDGFRA & APOA1 are expressed both in primary and secondary yolk-sac. TTR, APOB & GSTA1 are expressed more specifically in the secondary Yolk-sac consistent with findings in marmoset ^90^. STC1, LHX1 are absent in secondary yolk-sac cluster, but not in primary yolk-sac consistent with findings in marmoset ^90^. The co-expression of DKK1 and LHX1 detected via scRNA-seq alongside CER1 expression among some of the cluster 9 SOX17+ YS cells (arrows) marks AVE cells. **b,** Normalized expression of key extra-embryonic mesodermal genes projected on SEM UMAP. Cells in all four ExEM clusters (0, 2, 6 and 8) express general mesodermal genes such as HAND1, LEF1 and BMP4, while only cells in clusters 0 and 6 express typical extra cellular matrix related genes such as collagen genes and POSTN. **c,** differentially expressed genes between extra-embryonic mesoderm (ExEM) clusters 2 and 8 vs. (ExEM clusters 0 and 6. Most significant differentially expressed genes (adjusted p-value < 0.05 & asb(log2Fold-change)>1) are marked in red. Clusters 2 and 8, termed Early ExEM cells, expressed much lower level of extra cellular matrix related genes and remodeling enzymes, indicative of their putative relatively less mature cellular state.

## Supplementary Spreadsheet Legends

**Supplementary Spreadsheet 1. Summary of the microscopy parameters used for imaging in this study.** The spreadsheet provides information of the type of the microscope, used detection objectives, laser lines, and the voxel size for acquisition of the images published herein.

**Supplementary Spreadsheet 2 - PCR primers used in this study.**

Primer names, DNA sequences from 5’ to 3’ end, and the cited reference (when relevant).

**Supplementary Spreadsheet 3. Human SEM scRNA-seq analysis related gene expression list.**

Top markers (average log2(Fold-change)>0.25) of each of the 13 cell clusters in human SEMs, as identified by Seurat package.

## Supplementary Video Legends

**Video S1. 3D reconstruction of the human SEM at day 8**.

The video shows 3D rendering of the embryonic and extraembryonic tissue structures comprising day 8 human SEM. Immunofluorescence for epiblast (OCT4, cyan), yolk sac (SOX17, yellow), trophoblast (CK7, magenta), and nuclei (DAPI, white). 0 – 8 sec, 3D view of the SEM shape and the outer trophoblast with enlarged multinuclear cells (also seen on the slices form 22 sec). 8 – 22 sec and 39 – 42 sec, 3D segmentation of the epiblast (cyan) and hypoblast (yellow) is shown with the DAPI immunofluorescence signal on the background. 22 sec – 38 sec, slicing through the 3D volume showing the inner structure of the SEM with bilaminar disk with amnion, connected to the outer trophoblast, the yolk sac with the cavity, and the surrounding connective tissues with extensive intercellular space. Immunofluorescence signal and tissue segmentation are outlined. The image was acquired with Zeiss Z7 light-sheet microscope (see Methods) and processed with Imaris v10.0.0.

**Video S2. 3D reconstruction of the human SEM at day 6 demonstrating pro-amniotic cavity formation within the epiblast**.

3D reconstruction of the human SEM at day 6 shows 3D shape of the day 6 SEM (0 – 12 sec) and early formation of the proamniotic cavity in the center of the epiblast (12 – 18 sec). Immunofluorescence for OCT4 (epiblast, cyan), F-actin (red), and nuclei (DAPI, white). The image was acquired with Zeiss LSM 800 microscope (see Methods) and processed with Imaris v10.0.0.

**Video S3. 3D reconstruction of the human SEM at day 8 shows embryonic disk and amnion formation**.

3D reconstruction of the human SEM at the day 8, showing embryonic disk (SOX2, cyan) and amnion (TFAP2C, magenta). The segmentation of epiblast (cyan) and amnion (pink) is denoted as the semi-transparent outline together with the immunofluorescence signal. 17 – 21 sec, human SEM epiblast has a disk shape. The image was acquired with Zeiss Z7 light-sheet microscope (see Methods) and processed with Imaris v10.0.0.

**Video S4. 3D reconstruction of the human SEM at day 6 shows yolk sac morphology.**

3D reconstruction of the human day 6 SEM showing yolk sac development (marked by SOX17, yellow). 5 – 18 sec, zoom into the visceral and parietal yolk sac cells having columnar and squamous cell shape, respectively, and apical cell polarity (aPKC, green). F-actin (red), nuclei (DAPI, white). The image was acquired with Zeiss LSM 700 microscope (see Methods) and processed with Imaris v10.0.0.

**Video S5. 3D reconstruction of the human SEM at day 8 shows extraembryonic mesoderm cells integration underneath the yolk sac.**

3D reconstruction of the human SEM at the day 8 showing extraembryonic mesoderm cells located underneath the yolk sac (yellow) and marked by expression of VIM (red). OCT4 (epiblast, cyan), nuclei (DAPI, grey). The image was acquired with Zeiss LSM 700 microscope (see Methods) and processed with Imaris v10.0.0.

**Video S6. 3D reconstruction of human SEM at day 8 shows development of the syncytial trophoblast and lacunae formation.**

3D reconstruction of the human SEM at the day 8 showing development of the syncytial trophoblast, expressing both SDC1 (magenta) and HCGB (green) with multiple lacunae. F-actin (red), nuclei (DAPI, grey). 11 – 19 sec, slicing through the 3D volume showing inner structure of the multiple trophoblast lacunae. The image was acquired with Zeiss LSM 700 microscope (see Methods) and processed with Imaris v10.0.0.

## Methods

### Ethics

All experiments reported herein involving human PSCs were conducted following obtaining Weizmann Institute IRB approval (1868-2) to generate human **S**tem cell-derived **E**mbryo **M**odels (termed “SEMs” or “Stembroids”) from WIBR1, WIBR2 and WIBR3 human ESC lines ^128^ and from iPSC lines following obtaining donor informed consent to make genetically unmodified iPS lines from donated blood (Weizmann Institute IRB approval – 1871-2). All the experiments reported herein follow the latest ISSCR guidelines released in 2021^126^. This study does not involve derivation of new human ESC lines, does not use any newly obtained samples from fetal abortions, and does not use any newly donated human blastocysts. Further, this study does not involve in utero transfer of any human SEMs into any other species, consistent with ISSCR guidelines and Israeli legislation. Finally, all the human SEMs described herein do not correspond to developmental stages beyond 14 dpf.

### Data reporting

No statistical methods were used to predetermine sample size. Samples were randomly allocated when placed in the different growth conditions. Other experiments were not randomized. The investigators were not blinded to allocation during experiments and outcome assessment since there was no relevant scientific reason to do so.

### Stem cell lines

The following already established human ESC lines were used: WIBR3 (WT female), WIBR2 (WT female) and WIBR1 (WT male) human ES lines ^128^. Genetically unmodified JH33 and JH22 human induced pluripotent stem cell (iPSC) lines made from a healthy adult Caucasian/Middle Eastern male were also used to generate human SEMs. Previously established mouse Tet-ON iGata4 KH2-WT ESC line was used for comparing induction efficiency of PDGFRa^+^ cells ^27^. All cell lines were routinely checked for Mycoplasma contaminations every month (Lonza–MycoAlert), and all samples analyzed in this study were not contaminated.

### Human naïve ESC in vitro culture conditions

Golden stocks of human ESCs were cultured on feeder layer of irradiated mouse embryonic fibroblast (MEFs) and maintained in conventional human FGF/KSR primed conditions. FGF/KSR conditions on MEF substrate: DMEM-F12 (Invitrogen 10829) supplemented with 20% Knockout Serum Replacement (Invitrogen 10828-028), 1mM GlutaMAX (Gibco 35050061), 1% nonessential amino acids (BI 01-340-1B), 1% Sodium-pyruvate (BI 03-042-1B), 1% Penicillin-Streptomycin (BI 03-031-1B) and 8 ng/mL bFGF (Peprotech 100-18B-1MG).

To reprogram primed PSCs to a naïve state, human PSCs were maintained and expanded as in Bayerl et al ^35^, in serum-free HENSM on plates coated with 1% Matrigel (corning 356231) or MEF/gelatin-coated plates. At least three passages in HENSM conditions were applied before cells were used for experiments. Human naïve ESC lines were used for up to 10 passages in HENSM conditions. For maintenance of ESCs in naïve HENSM conditions, cells were passaged every 3-5 days using TryplE (Gibco 12604054). Naïve and primed hPSCs were expanded and induced into different lineages in a 5% CO_2_ incubator at 5% O_2_ at 37C.

### Generation of iGATA4, iGATA6, iCDX2 and iGATA3 human ESCs clones

To generate Tet-ON inducible lines, we employed a PiggyBac plasmid expressing cDNA insert adba transposase vector of choice under the control of a doxycycline-inducible promotor (a kind gift from Volker Busskamp, Addgene plasmid #104454). The donor vector carries M2Rtta and a site for cDNA insert of transcription factor of interest. We used this vector to generate 4 different DOX inducible lines in WIBR3 WT human female ESCs: human iCDX2 or human iGATA3 (to promote human PSC differentiation towards trophectoderm) and human iGATA4 or human iGATA6 (to promote human primitive endoderm/ extra embryonic mesoderm priming from human PSCs). Puromycin selection was applied for approximately 6-8 days. Resistant clones were picked and cultured for downstream characterization. Insertion was validated by immunostaining after DOX (2µg/ml) induction of the gene of interest. Transgene expression was verified to be specifically detected only after DOX addition in the corresponding lines. Detailed generation, characterization and validation of these lines can be found on (Extended Data Figure 1, 3, 7). Generation of fluorescent labeled WIBR3 iCDX2 line was made after transduction with lentivirus constitutively expressing tdTomato protein. For lentivirus generation, HEK293T cells were plated on 10 cm dishes filled with 10 ml DMEM 10% FBS and Pen/Strep, at a density of 5.5 million cells per plate. On the next day, cells were transfected with Second-generation lentiviral vectors (Addgene 8455 and 8455), using X-tremeGENE 9 transfection reagent, along with 16 μg of the target plasmid for the transduced fluorescent protein (tdTomato). The supernatant containing the virus was collected 48hr following transfection, filtered using 0.45μm filter and concentrated by ultracentrifugation. Human PSCs were plated in mTESR medium on Matrigel coated 6-well plates at low density, next day they were transduced with lentivirus in the presence of protamine sulfate (10 µg/ml) for 6 hours, afterwards medium was exchanged. After 2 days, the infected human ESCs were expanded for 1 passage and the positive population was sorted using FACS and further expanded for experimentation.

### Derivation of a stable human TSC line from human ESCs

TSC lines were produced from Naïve (HENSM) (nTSC) and primed conditions (pTSC) according to Okae et al and Viukov et al, respectively ^70, 71^. Briefly, human naïve WIBR3 ESCs were expanded at least 3 passages in naïve or primed conditions and then transferred into TSC media (TSCm) on 1% Matrigel (corning 356231)-coated plates. After 3 passages, stable TSC lines could be established and could be passaged 70 times ^129, 130^. Cells were expanded in a 5% CO_2_ incubator at 5% O_2_. For maintenance, human TSC were passaged with TrypLE (Gibco 12604054) when reached 70%-80% confluency. Immunostaining was performed to confirm human TSC identity: human TSC cells were negative for CDX2, and positive for Cytokeratin7 (CK7) GATA3 and TFAP2C, consistent with previous reports. Only confirmed lines were used for SEM experiments. Human TSC media (TSCm) used herein was previously described in ^70^ with slight modifications: 470 ml DMEM/F12 (Invitrogen 21331), 5 ml Commercial N2 supplement (Invitrogen 17502048), 10 ml B27 supplement (Invitrogen 17504-044), 5 mL Sodium Pyruvate (Biological Industries 03-042-1B), 5 mL Penicillin/Streptomycin (Biological Industries 03-033-1B) 5ml (Biological Industries 03-033-1B), 5 mL GlutaMAX (Invitrogen 35050061), 5 mL NEAA (Biological Industries 01-340-1B), 50 μg/ml L-ascorbic acid 2-phosphate (Sigma A8960), 50ng/ml Human EGF (Peprotech AF-100-15), 0.75-1 µM TGFRi A83-01 (Axon 1421), 2 µM GSK3i CHIR99021 (Axon Medchem 1386), and 5 µM ROCKi Y27632 (Axon 1683).

### Derivation of human trophectoderm (TE) from human naïve PSCs for SEM generation

Human TE cells were obtained from human naïve PSCs expanded in human HENSM conditions for at least 3 passages, 24 hours before the induced cells were plated in HENSM on 1% Matrigel (corning 356231)-coated plates supplemented with 10 µM ROCKi Y27632 (Axon 1683), next day the protocol was started. The 3-day protocol for human TE induction was previously described and is adapted from Io et al^36^ as follows: BAP(J) media was used for 72 hours. This medium consisted on 2 µM TGFRi A83-01 and 2 µM MEKi/ERKi PD0325901 base, which was complemented the first 24h with 10ng/ml Human recombinant BMP4 (Peprotech) and then substituted with for 1µM JAK inhibitor I (Calbiochem 420099) on day 2 and 3. The base medium consisted on: 470 ml of 1:1 mix of Neurobasal (Invitrogen 21103-049) and DMEM/F12 (Invitrogen 21331), 5 ml penicillin-streptomycin (Biological Industries 03-033-1B), 5 ml GlutaMAX (Invitrogen 35050061), 5 ml NEAA (Biological Industries 01-340-1B), 5 ml Sodium Pyruvate (Biological Industries 03-042-1B), 10 ml B27 supplement (Gibco 17504-044), 5 ml N2 supplement (Invitrogen 17502048), 2 µM TGFRi A83-01 (Axon Medchem A83-01), 2 µM MEKi/ERKi PD0325901 (Axon Medchem 1408).WT human naïve PSCs or iCDX2/iGATA3 cell lines were tested for human TE induction in the presence or absence of DOX as indicated. All the process was incubated in a 37C incubator with 5% O_2_ and 5% CO2.

### Primitive Endoderm (PrE) and Extra-Embryonic Mesoderm (ExEM) induction from human naïve PSCs

Pre/ExEM cells were induced from human naïve PSCs expanded in HENSM conditions for at least 3 passages as described above. For induction HENSM cells were plated on gelatin-MEF coated plates the day before induction with 10 µM ROCKi Y27632, next day medium was changed to RCL for 72h^131^. RCL is composed of: 480 ml RPMI media (GIBCO 21875-03), 10 ml B27 minus insulin supplement (Invitrogen A18956-01), 1 mM GlutaMAX (Invitrogen), 1 % penicillin-streptomycin (Invitrogen), 3 µM CHIR (Axon Medchem 1386) and 10 ng/ml LIF (Peprotech 300-05). RCL medium contains the same composition as RACL but without adding recombinant Activin. WT human naïve PSCs or iGATA4/iGATA6 cell lines were employed for induction in the presence or absence of DOX as indicated. All the process was incubated in a 37C incubator with 5% O_2_ and 5% CO_2_.

### Generation of human stem cell embryo models (SEMs)

A step-by-step detailed protocol will be accompanying this work upon final publication. Briefly,To generate SEMs from human naïve PSCs, three starting cell mixtures were co-aggregated using AggreWell 24-well plate 400 (STEMCELL Technologies 34415):

1. Naïve PSC (nPSC) WT cells cultured in HENSM medium in a 5% CO_2_ incubator at 5% O_2_ at 37C.
2. For the primitive endoderm and extra-embryonic mesoderm compartments (PrE/ExEM), naïve WT cells were plated on irradiated MEF (mouse embryonic fibroblast conditions)/Gelatin coated plates in HENSM supplemented with ROCKi 10 µM (Axon Medchem 1683). The next day, cells were washed with PBS twice (without harvesting), and HENSM was replaced by RCL medium. RCL was kept for 72h with 24h medium exchanges in a CO_2_ incubator at 5% O_2_ at 37C.
3. For the trophectoderm lineage, naïve WT cells were plated on feeder free conditions (Matrigel) in HENSM supplemented with ROCKi 10 µM. The next day, cells were washed with PBS twice (without harvesting), and HENSM was replaced with BAP medium for 24 hours, following by replacement with APJ for another 48 hours in a 5% CO_2_ incubator at 5% O_2_ at 37C (termed BAP(J) protocol).

Co-aggregation was defined as time point 0 of the protocol. 12-24h before aggregation all donor cells were supplemented with ROCKi 10 µM (Axon Medchem 1683). At the day of aggregation (day 0), AggreWell 400 24-well plate preparation was done according to manufacturer instructions. Briefly, 500 µl of anti-adherence rinsing solution (STEMCELL Technologies 07010) was added to each well, the plate was centrifuged at 2,000g for 5 minutes and incubated 30 min at room temperature. Subsequently, rinsing solution was removed and the plate was washed with PBS. Each well was filled with 500 µL of aggregation medium and kept at 37° C for medium equilibration. Aggregation medium (BSA supplemented N2B@7 media) consisted in 500ml 1:1 mix of Neurobasal (Invitrogen 21103-049) and DMEM/F12 (Invitrogen 21331), 5 ml penicillin-streptomycin (Biological Industries 03-033-1B), 5 ml GlutaMAX (Invitrogen 35050061), 5 ml NEAA (Biological Industries 01-340-1B), 5 ml Sodium Pyruvate (Biological Industries 03-042-1B), 10 ml B27 supplement (Invitrogen 17504-044), 5 ml N2 supplement (Invitrogen 17502048), 1 ml β-mercaptoethanol 50mM (Gibco 31350-010), 2.25ml of BSA solution 35% (Merck 9048-46-8).

The three cell populations were collected with TrypLE (Thermo Fisher 12604054) (3 minutes for the HENSM and RCL-induced cell populations, and 5 minutes for the BAP(J)-induced cells) at 37° C, Afterwards TrypLE was removed with vacuum and the cells were incubated for two minutes at room temperature and cells were subsequently collected with PBS. Cells were centrifuged at 1300 rpm for 3-5 minutes and resuspended in aggregation medium. Next, RCL-induced cells were plated on gelatinized tissue culture plates on MEF medium consisting on 500ml DMEM (Gibco 41965-039) 20% FBS (Sigma, F7524-500ml), 5 ml penicillin-streptomycin (Biological Industries 03-033-1B), 5 ml GlutaMAX (Invitrogen 35050061), 5 ml NEAA (Biological Industries 01-340-1B), 5 ml Sodium Pyruvate (Biological Industries 03-042-1B), for MEF depletion for 30 minutes at 37° C. At the end of MEF depletion, the supernatant was collected and passed through a 70uM cell strainer, and all three cell types were centrifuged separately and resuspended and passed through a 70 µM cell strainer in N2B27 medium. The three cell fractions were counted and combined as follows in an Aggrewell 400 plate: Ratio of 1:1:3 (HENSM: RCL: BAPJ) or (Epi: PrE/ExEM: TE) = 28800 Epi (HENSM) cells, 28800 Pre/ExEM (RCL) cells, and 86400 TE (BAPJ) cells per 24-well. Cell number per single microwell = 120 cells. Cells were prepared on a 2x concentration with 20 µM ROCKi, 500 µl of cell-mix suspension was gently added drop wise to each well of the AggreWell plate (1ml final volume 10uM final ROCKi concentration). The plate was centrifuged at 100g for 3 minutes and incubated at 37 °C in hypoxic conditions (5%O_2_ and 5% CO_2_)

Next day (day 1), 900 µL of medium were gently removed from each well and replaced with 1 ml of pre-equilibrated aggregation (N2B27-BSA) medium. The same volume of medium was replaced at day 2. At aggregation day 3, aggregates were gently transferred to 6-well cell suspension culture plates (Greiner, 657185) filled with 3 ml of pre-equilibrated EUCM2 (20% FBS) per well and placed on an orbital shaker rotating at 60 rpm (Thermo Scientific 88881102 + 88881123) located inside a 5% CO_2_ incubator in 20% O_2_. On day 4, 2 ml of medium were gently removed per well and were replaced with 2 ml of pre-heated EUCM 2 (with 30% FBS). Same procedure was repeated on day 5, refreshing with 2 ml of EUCM2. After 6 days, 2 ml of medium were gently removed per six-well and were replaced with 2 ml of pre-heated EUCM2 with 50% FBS. Same procedure was repeated at day 7 and cultures were finished at day 8 post-aggregation. Alternatively, human SEMs can be cultured from day 6 to 8 using the roller culture platform adapted to an electronic gas regulator module using EUCM2 50% FBS with similar outcome. EUCM2 is formulated as follows: Advanced DMEM/F12 (GIBCO 21331-020), extra added 1 mM Sodium pyruvate (Sigma-Aldrich, S8636), 0.5% CMRL media (GIBCO 11530037), extra added 1 mg/mL D(+)-Glucose Monohydrate (J.T. Baker - 0113) (e.g. add 500mg per 500mL media), 1 mM GlutaMAX (GIBCO, 35050061), 1% penicillin streptomycin (Biological Industries – Sartorius 03-031-1B), 1x of ITS-X supplement (Thermo Fisher Scientific 51500-056), 8 nM B-estradiol (Sigma-Aldrich, E8875), 200 ng/ml progesterone (Sigma-Aldrich, P0130), 25 µM N-acetyl-L-cysteine (Sigma-Aldrich, A7250), 20-50% FBS (Sigma Aldrich F7524 – heat inactivated and filtered) as indicated in **Fig. 2b**. Culture media was pre-heated for at least an hour by placing it inside a CO_2_ incubator at 37°C.

### Human blastoid generation and PALLY/PALY conditions

Human blastoids were generated according to Kagawa et al ^22^ with few modifications. WIBR3 hESCs were grown in HENSM for at least 3 passages on feeder free conditions (1% Matrigel coated plates) and were used for generating human blastoids. After 3 days of growth of naïve PSC were harvested and counted and 55 cells were seeded per microwell (total of 66,000 cells were seeded per 24-well in 1 ml of medium) AggreWell 400 24-well (Stemcell Technologies cat 34415) in N2B27-BSA supplemented with 10 µM ROCKi Y27632. The next day medium was changed to PALLY consisting of N2B27 base with 1 µM MEKi/ERKi PD0325901 (Axon Medchem 1408), 1 µM TGFRi A83-01 (Axon Medchem A83-01), 1 µM LPA (Tocris, 3854), LIF 10ng/ml and 10 µM ROCKi Y2763. This medium was repeated on day 2, but on day 3 medium was changed for LY (1 µM LPA (Tocris, 3854) and 10 µM ROCKi Y27632) for another 48h. Afterwards human blastoids were manually selected and collected for further analysis. The en tire was conducted in 5% O2 and 5% CO2 conditions.

### Mouse SEM generation

Mouse SEM aggregations were made according to Tarazi et al ^27^. Mouse animal experiments pertained only to mouse SEM and were performed according to the Animal Protection Guidelines of Weizmann Institute of Science and approved by the following Weizmann Institute IACUC (#01390120-1, 01330120-2, 33520117-2). Mouse aggregation medium was also tested for human SEMs but was found inappropriate. Mouse aggregation medium (mouse AM) consisted of 1x DMEM (GIBCO-41965) supplemented with 20% FBS (Sigma), 1 mM GlutaMAX (GIBCO, 35050061), 1% penicillin streptomycin (Biological Industries – Sartorius 03-031-1B), 1% Sodium Pyruvate (Biological Industries – Sartorius 03-042-1B), 1% non-essential amino acids (Biological Industries – Sartorius 01-340-1B) and 0.1 mM β-mercaptoethanol (Thermo 31350010).

### Morphological evaluation of human early development and efficiency calculations

Assessment of appropriate human development was performed by careful analysis of available *in-utero* histological embryo collections (predominantly Carnegie collection), taking in account different epithelium morphology and structure organization through different stages of development. Furthermore, available work on primate development was employed as a reference for anatomical structure ^132–134^, specific markers of each of the compartments was inferred from previous ex-utero human development works ^135, 136^, in-vitro differentiation protocols and primate existing databases^132^.

Most of human histological descriptions and figures used for this paper are mentioned in the virtual human embryo website (https://www.ehd.org/virtual-human-embryo/) and available human embryology textbooks (Langman, Larsen, Carlson).

All human in utero data and figures included in this study were made only from Carnegie collections after obtaining the appropriate copyright approvals (in process). Only SEMs presenting all the previously defined features were considered as properly developed. Percentage of human SEMs generation is calculated based on the number of properly developed structures observed per random fields of view at a specific time point on independent experiments, while always relying on immunofluorescence to corroborate the 3-lineage contribution.

### Quantification and Statistical Analysis

Statistical analyses of real time PCRs were performed in QuantStudio software v1.3 and visualized in GraphPad Prism 7. Visualization and statistical analyses of the cell numbers and SEM efficiencies were performed with Python 3.8.5 software using scipy and seaborn libraries. Boxplot graphs indicate medians with interquartile ranges, the whiskers mark distribution range. The barplots show average values plus s.d. The dots mark individual numerical values used for visualization of the data distributions and analyses. Significant difference between two samples was evaluated by the two-sided Mann-Whitney test for non-normally distributed data. p < 0.05 was considered as statistically significant.

### Immunofluorescence

Cells were fixed in 4% paraformaldehyde in PBS at RT for 10 minutes. Samples were then washed 3 times in PBS, permeabilized in PBST (PBS with 0.1% Triton X-100) for 10 min, blocked in PBS/0.05% Tween/5% fetal bovine serum/1% bovine serum albumin for 1h and incubated with primary antibodies diluted in blocking solution at 4°C overnight. Subsequently, cells were washed in PBS/0.05% (three times, 5 min each) and incubated with Alexa Fluor (488, 568 and/or 647)-conjugated secondary antibodies (Jackson ImmunoResearch) diluted in blocking solution (1:200). Samples were counterstained with 1 μg/ml DAPI for 10 min at RT, washed with PBS three times (5 min each) and mounted with Shandon Immuno-Mount (Thermo Scientific). The antibodies and dilutions employed for cell immunofluorescence were the following: Rabbit polyclonal anti-Cdx2 (Cell Signaling Cat# 3977) 1:200; Mouse monoclonal anti-Cdx2 (Biogenex Cat# MU392A-UC) 1:200; Rabbit polyclonal anti Gata4 (Abcam Cat# Ab84593) 1:120; Rabbit monoclonal anti-Foxa2 (Abcam Cat# Ab108422) 1:100; Goat polyclonal anti-Sox17 (R&D Cat# AF1924) 1:200; Mouse monoclonal anti-Oct4 (clone C-10) (Santa Cruz Cat# SC-5279) 1:200; Rabbit monoclonal Cdx2 (Abcam Cat# ab76541) 1:200; Goat Monoclonal Tfap2c (R&D Cat# AF5059) 1:200; Rabbit monoclonal Cytokeratin 7 (Abcam Cat# ab181598) 1:200; Rabbit monoclonal Cytokeratin 7 (Abcam Cat# ab68459) 1:200; Rabbit monoclonal Nanog (Abcam Cat# ab109250) 1:200; Mouse monoclonal Gata3 (Invitrogen Cat# MA1-028) 1:200; Goat polyclonal Gata3 (R&D Cat# AF2605) 1:200; Mouse monoclonal HCG-Beta (Abcam Cat# ab9582) 1:200; Rabbit monoclonal Gata6 (Cell signaling Cat# 5951) 1:200; Goat polyclonal Oct3/4 (R&D Cat# AF1759) 1:200; Goat polyclonal Gata6 (R&D Cat# AF1700) 1:200; Rabbit monoclonal Syndecan1 (Abcam Cat# ab128936) 1:500; Rabbit monoclonal PDGFR-A (Abcam Cat# ab134123) 1:100; Goat polyclonal Nidogen2 (R&D Cat# AF3385) 1:100.

### Flow cytometry

Flow cytometry analysis were done on a BD FACS-Aria III. Cells were harvested with TrypLE and washed once with PBS afterwards they incubated for half an hour with conjugated primary antibodies (5ul) on 100ul PBS/0.5% BSA. The primary antibodies used are as follows: Mouse monoclonal TROP2-488 labeled (R&D Cat# FAB650G); Mouse monoclonal TROP2-PE labeled (R&D Cat# FAB60P); Mouse monoclonal CD249 (ENPEP)-BV421 labeled (BD Cat# 744872); Rat monoclonal anti mouse CD140a (PDFGR-a)-PE/Cy7 labeled (BioLegend Cat# 135912); Mouse monoclonal anti human CD140a (PDFGR-a)-PE/Cy7 labeled (BioLegend Cat# 323508); Mouse monoclonal anti human CD140a (PDFGR-a)-APC labeled (BioLegend Cat# 323512). FSC and SSC singlets were gated, and only single cells were considering for all analyses. An unstained control was employed to determine the negative/positive populations for all antibodies, ensuring that 100% of the unstained population was allocated on the negative area of the histogram/dot plot.

### Confocal microscopy

The immunofluorescence images were acquired using a Zeiss LSM 700, as well as a LSM 800 inverted confocal microscopes (Zeiss), both equipped with 405 nm, 488 nm, 555 nm and 635 nm solid state lasers, using a Plan-Apochromat 20× air objective (numerical aperture 0.8) or an EC Plan Neofluar 10× air objective (numerical aperture 0.3). For a detailed description of the imaging parameters, see **Table S1**. Images and maximum intensity projections were processed using Fiji ^137^, Zen 2 blue edition software 2011 (Zeiss), and Adobe Illustrator CC.

### Light-sheet microscopy

The immunofluorescence images were acquired using Zeiss Z7 light-sheet microscope, equipped with 405 nm, 488 nm, 561 nm, and 638 nm lasers, using a single water 20x Plan-Apochromat (numerical aperture 1.0) detection objective (Zeiss) and two air 10x Plan-Apochromat (numerical aperture 0.2) illumination objectives (Zeiss). The single sample was imaged at a time, mounted in 1% low-melting temperature agarose inside the imaging chamber filled with PBS. Light-sheet volumes along the Z-axis were acquired in a dual scanning mode, using a pivot scan. Light-sheet thickness was set to 3.77 µm, and laser power in the 1 – 50% range was applied. Frame size, 1920×1920 px, exposure time, 50 msec. The light-sheets for left- and right-side illuminations were adjusted independently inside the sample volume for each channel based on the signal intensity in the focal plane of the detection lens. See also **Table S1**.

### Electronically controlled ex utero roller culture platform

Human SEMs can be kept in the ex utero electronically controlled roller culture platform after day 6 ^138^ which provides continuous flow of oxygenating gas. The system consists of an electronic gas modulating unit (Arad Technologies) adapted to the roller culture unit from B.T.C. Engineering, – Cullum Starr Precision Engineering Ltd - UK)^139^, as previously described ^138^. On day 7, all human SEMs from one well of the 6-well plate were picked and transferred to glass culture bottles (50-100 structures per bottle) containing 4 mL of fresh EUCM2 50% FBS. The bottles were placed on the rolling culture system, rotating at 30 revolutions per minute at 37°C, and continuously gassed with an atmosphere of 21% O_2_, 5% CO_2_ at 6.5-8 pounds per square inch (psi). Bottles were kept inside glass culture bottles rotating on a spinning wheel allocated inside a “precision” incubator system (BTC01 model with gas bubbler kit - by B.T.C. Engineering, – Cullum Starr Precision Engineering Ltd - UK). (BTC 04). Gas flows from the gas mix box through the inlet into the humidifier water bottle, and then to the inside of the bottles in the rotating drum. The rate of bubbles created inside an outlet-test tube filled with water is used to monitor speed of the gas flow. The bubble rate was adjusted using the valve on the lid of the water bottle to the first point where continuous bubbling is observed, which generally corresponds to 0.06-0.1 psi and 50-75ml/min gas flow. A black cloth was used cover the incubator to provide protection against phototoxicity. 2 ml of EUCM2 with 50% FBS was used in the roller culture for human SEMs at these stages.

### RNA extraction & RT-PCR analysis

Total RNA was isolated using RNeasy mini kit (Qiagen) following manufacturer instructions. After, 1 μg of total RNA was reverse transcribed using a High-Capacity Reverse Transcription Kit (Applied Biosystems). RT-PCR was performed in triplicate using SYBR Green PCR Master Mix (Qiagen) and run on Viia7 platform (Applied Biosystems). Values were normalization to Actin and/or Gapdh across all experiments. Data are presented as fold difference compared to reference sample, which is set as 1. RT-PCR primer list used listed in **Table S2.**

### Chromium 10X single cell RNA sequencing

Human SEMs grown ex utero were manually selected and harvested for single cell RNA sequencing (**Table S3**) using the Chromium Next GEM Single Cell 3’ platform (V3.1). All human SEMs analyzed by scRNA-seq were generated by co-aggregating WT WIBR3 naïve ESC lines grown in HENSM with RCL-induced or BAPJ-induced WT cells. All SEMs samples were processed including extraembryonic compartments without any dissection. SEMs were dissociated with Trypsin-EDTA solution C (0.05%) for 10 minutes (Biological Industries; 030501B). Trypsin was neutralized using media with 10% FBS, and cells were washed and resuspended in 1x PBS with 400 µg/ml BSA. Cell suspension was filtered with a 100 µm cell strainer to remove cell clumps. Cell viability of at least 90% was determined by trypan blue staining for all samples. Cells were diluted at a final concentration of 1000 cells/µL in 1x PBS with 400 µg/ml BSA. scRNA-seq libraries were generated using the 10x Genomics Chromium v3.1 Dual Index system (5000 cell target cell recovery) and sequenced using Illumina NovaSeq 6000 platform according to the manufacturer’s instructions.

### 10X Single cell RNA-seq analysis

10x Genomics data analysis was performed using Cell Ranger 7.1.0 software (10x Genomics) for pre-processing of raw sequencing data, and Seurat 4.3.0 for downstream analysis. To filter out low-expressing single cells, possible doublets produced during the 10x sample processing, or single cells with extensive mitochondrial expression, we filtered out cells with under 1000 expressing genes, over 8,000 expressing genes and over 15% mitochondrial gene expression. Seurat integrated analysis and anchoring of all individual samples was performed and then normalized by log-normalization using a scale-factor of 10,000. The top 2,000 variable genes were identified by the variance stabilizing transformation method, and subsequently scaled and centered. Principal components analysis was performed for dimensional examination using the ‘elbow’ method. The first 10 dimensions showed the majority of data variability. Therefore, UMAP dimensional reduction was performed on the first 10 dimensions in all samples. Clusters were detected using Seurat Find Clusters function, with resolution parameter =0.5. Dot-plot describing expression and prevalence of specific genes was generated using Seurat DotPlot() function. Projection of selected genes on SEM UMAP was generated with Seurat FeaturePlot() function.

### Projection on Human Embryo Reference

The human embryo reference was built by integrating previously published datasets consisting of 5 human embryonic data sets spanning in vitro cultured human blastocysts ^24, 118, 119^, 3D-in vitro cultured human blastocysts until pre-gastrulation stages ^120^, and a Carnegie Stage 7 (CS7) 16-19dpf human gastrula ^117^, as previously described ^101^ (manuscript in final preparation). The raw counts for cells of human SEMs were aggregated within neighborhood nodes as calculated by Milo (Dann et al 2022) resulting in 945 nodes, followed by projecting the summed counts matrix onto the assembled Human embryo reference (**Fig. 6e**).

### Mouse Extra-embryonic annotation analysis

Rhox5 positive cells (>1 counts) were chosen for the analysis of extra-embryonic tissue. The cells were annotated based on marker genes as previously conducted in ^77^, such that if at least 4 markers (3 in the case of SpA-TGC and SpT-Gly) were expressed (>0 counts), the cell was annotated in that category. 26% of the annotated cells, were annotated by multiple categories. The markers are as following: Chorion (Irx4, Esx1, Id1, Id3, Phlda2, Klhl13), Chorion progenitors (Sox3, Dusp6, Nat8l, Bmp4, Sox2, Esrrb, Eomes), Intermediate Chorion (Ascl2, Fgfr2, Cited1, Gjb3, Ndrg1, Irx2, Irx3), uncommitted EPC (Chsy1, Gjb3, Krt19, Lgals1, Cald1, Ctsl), SpA-TGC (Ctla2a, Pecam1, Ramp3, Igfbp7, Nos3), SpT-Gly (Dlx3, Car2, Ncam1, Pcdh12, Tpbpa), TGC-progenitors (Adm, Fosl1, Hand1, Trpm5, Maged2, Prl5a1), p-TGC (Star, Serpinb9d, Hsd3b6, Rhox6, Cts7) as in ^77^. R ggplot was used to generated scatter plot, along with geom_smooth(method=“lm”).

### Mouse and Human IGV Analysis

Bulk ATAC-seq and RNA-seq profiles in the proximity of selected genes (GATA3, GATA4, GATA6) are presented using IGV genome browser. Mouse datasets were taken from published datasets ^140–142^ and https://www.ncbi.nlm.nih.gov/geo/query/acc.cgi?acc=GSE181053. and so were human datasets ^125, 143, 144^. Mouse enhancers were taken from ^141^, human enhancers were taken from GeneHancer ^145^. Open regions that overlap with promoter or exon were excluded from the analysis. Potential enhancers in these regions were manually curated.

### Data and code availability

All scRNA-seq data are deposited under GEO: XXXXXXXXX

Any other information required to reanalyze the data reported in this work is available upon request from the corresponding authors upon request authors.

